# Identification of Complex Multidimensional Patterns in Microbial Communities

**DOI:** 10.1101/540815

**Authors:** Golovko George, Khanipov Kamil, Albayrak Levent, Fofanov Yuriy

**Affiliations:** Department of Pharmacology and Toxicology, University of Texas Medical Branch–Galveston, Galveston, TX 77555-0144, USA; Sealy Center for Structural Biology and Molecular Biophysics, University of Texas Medical Branch–Galveston, Galveston, TX 77555-0144, USA

## Abstract

**Motivation:** Identification of complex relationships within members of microbial communities is key to understand and guide microbial transplantation and provide personalized anti-microbial and probiotic treatments. Since members of a given microbial community can be simultaneously involved in multiple relations that altogether will determine their abundance, not all significant relations between organisms are expected to be manifested as visually uninterrupted patterns and be detected using traditional correlation nor mutual information coefficient based approaches.

**Results:** This manuscript proposes a pattern specific way to quantify the strength and estimate the statistical significance of two-dimensional *co-presence*, *co-exclusion,* and *one-way relations* patterns between abundance profiles of two organisms which can be extended to three or more dimensional patterns. Presented approach can also be extended by including a variety of physical (pH, temperature, oxygen concentration) and biochemical (antimicrobial susceptibility, nutrient and metabolite concentration) variables into the search for multidimensional patterns. The presented approach has been tested using 2,380 microbiome samples from the Human Microbiome Project resulting in body-site specific networks of statistically significant 2D patterns. We also were able to demonstrate the presence of several 3D patterns in the Human Microbiome Project data.

**Availability:** C++ source code for two and three-dimensional patterns, as well as executable files for the Pickle pipeline, are in the attached supplementary materials.

## 1. Introduction

Understanding the nature of microbial variation within environments and how this variation is altered in response to stimuli remains among the key challenges in the field at present. The same environment can be colonized by significantly different microbiomes, however, shifting microbial communities between “enterotypes” requires the identification of interaction patterns. Elucidation of complex multidimensional relations defining the shape and functionality of MC opens new opportunities for “personalized microbiome treatment”.

### 1.1 Non-continuous multidimensional patterns

To a certain extent, complex relations between microorganisms within microbial communities (MC) can be recovered by observing their abundances as well as monitoring how such abundances change in response to internal and external perturbations. While relations between some microorganisms can be manifested as continuous visually uninterrupted patterns and can be identified using mutual information based approaches such as MIC (Cover and Thomas, 1991), Pearson or Spearman correlation (Fieller, et al., 1957; Pearson, 1895), etc., not every significant relation in MC is expected to be expressed as a visually uninterrupted pattern. It seems obvious that some members of MC can be simultaneously involved in multiple relations which altogether will determine their abundance. In many environmental microbial communities for example, vital to whole community functions (e.g. nitrogen fixation) are often performed by single species so the abundance pattern between these organisms and other members of MC will not be represented by correlation, but rather is expected to exhibit a Boolean pattern (Saito, et al., 2011). A Boolean *one-way relation* pattern is exhibited where the presence of “dependent” microorganisms requires the presence of “provider”, but not vice versa (Figure 1b). Similarly, other pairwise relations such as *co-presence* and *co-exclusion* may be represented as non-continuous patterns (Figure 1a,c). In fact, out of 2^4^ possible combinations of the presence/absence profiles between two organisms, only four may be interpreted as possible relations: *co-presence*, *co-exclusion*, and two *one-way relations* (organism 1 needs organism 2 to survive and vice versa). It is also important to keep in mind that if the cooperation of several organisms is required to maintain a single metabolic pathway, their abundances will fit into multidimensional Boolean patterns, such as *co-presence* (Figure 1g).

**Figure 1.**
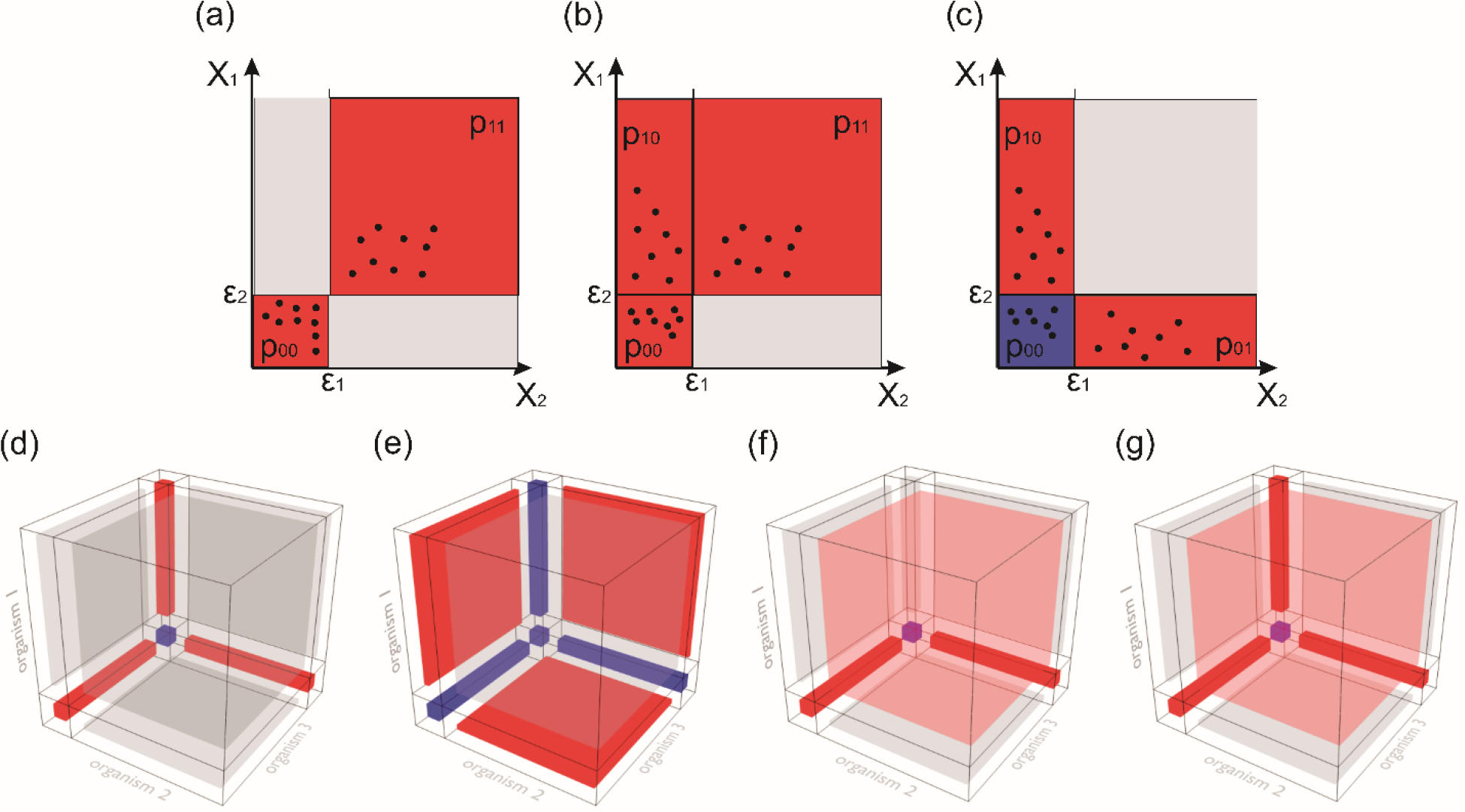
Examples of non-continuous two- and three-dimensional patterns. Two-dimensional patterns: (a) *co-presence*; (b) *one-way* ; and (c) *co-exclusion* patterns. X1 and X2 are abundances of microorganisms; *ɛ*_1_ and *ɛ*_2_ represent the presence/absence threshold; *p*_00_, *p*_01_, *p*_10_, and *p*_11_ the proportion of points (observation) located in each partition. Three-dimensional patterns: (d) Type 1 *co-exclusion*; (e) Type 2 *co-exclusion*; (f) a pattern when the presence of organism *X*_2_ changed patterns between *X*_1_ and *X*_3_ from *co-presence* to *co-exclusion*; and (g) represents the case where three organisms can be present only all together on one-by-one. Red color represents quadrants requiring the proportion of observation to exceed the minimal threshold. Red and blue quadrants are areas contributing to the pattern score.

### 1.2 Complications of using Mutual Information as a score for non-continuous patterns

In Boolean patterns, two microorganism’s abundances can vary without affecting the pattern (e.g., variation in abundance within the same quadrant), the pattern strength can be defined based on the fraction of observations located in four quadrants of two-dimensional space: *p*_00_, *p*_01_, *p*_01_, *p*_11_ (Figure 1a-c). However, it is important to mention that since different roles played by microorganisms in MC may require different minimal abundance, the appropriate calculation of the *p*_*ij*_ will require identification of the microorganisms specific thresholds so *p*_*ij*_ becomes a function of four variables: *p*_*ij*_ (*X*_1_, *X*_2_, *ɛ*_1_, *ɛ*_2_), where *X*_1_ and *X*_2_ are abundance profiles of two microorganisms under consideration and *ɛ*_1_ and *ɛ*_2_ are corresponding presence/absence thresholds.

The most obvious choice to define the strength of a Boolean pattern would be by using a mutual information score (MIS):

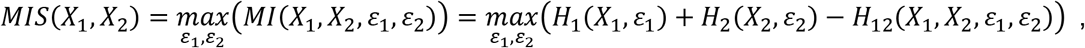

where *H*_*i*_(*X*_*i*_, *ɛ*_*i*_) and *H*_*ij*_(*X*_*i*_, *X*_*j*_, *ɛ*_*i*_, *ɛ*_*j*_) correspond to one- and two-dimensional entropies.

The use of mutual information to identify such patterns, however, has several significant disadvantages. Firstly, the best possible (maximal) MIS value is not the same for different pattern types. For example, while MIS value for the “ideal” *co-presence* and *co-exclusion* patterns is 0.693 (Figure 2a,b), the score for the “ideal” *one-way* pattern is only 0.174 (Figure 2c). Moreover, a small disbalance between fractions of points located in four partitions *p*_00_, …, *p*_11_ can significantly affect the MIS value. For example, while two *co-exclusion* patterns may be intuitively obvious (Figure 2a and 2d), a disbalance between *p*_00_ and *p*_11_ can cause signa ificant drop in the MIS value. Similar observation can be made for *co-exclusion* (Figure 2b and 2e). The use of MIS value can be especially misleading in case of *one-way* relation patterns. An MIS value may be extremely low (0.055) for patterns which can be clearly interpreted as *one-way* relation (Figure 2f). These observations suggest that the use of MIS to identify non-continuous Boolean patterns may result in missing certain intuitively obvious patterns. The presented work is an attempt to introduce an alternative, pattern-specific approach, to estimate the strength and statistical significance of two- and higher dimensional patterns between members of microbial communities.

**Figure 2.**
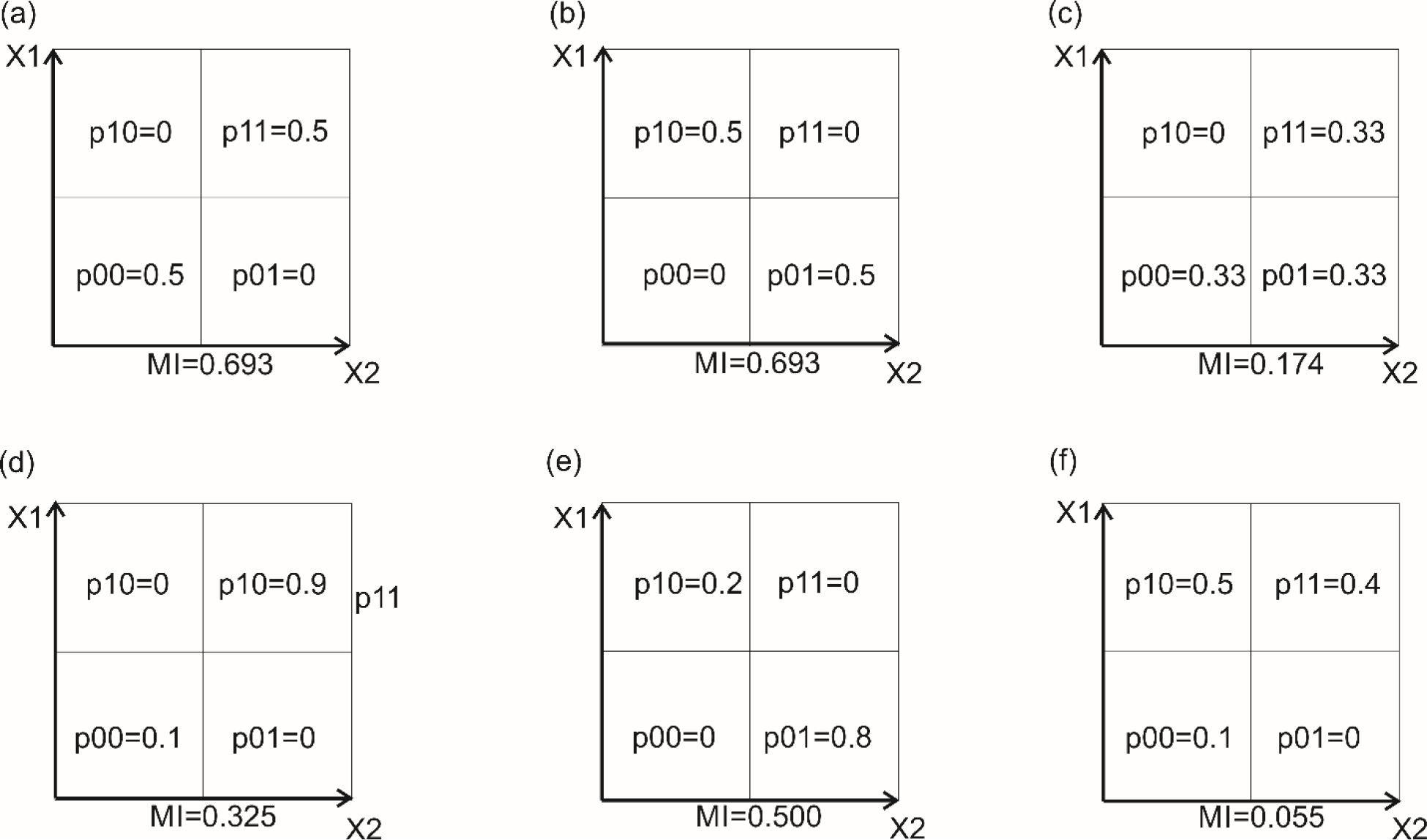
MIS values for “ideal” (a) *co-presence*; (b) *co-exclusion*; and (c) *one-way* patterns. Effect of disbalance on (d) *co-presence*; (e) *co-exclusion*; and (f) *one-way* relations on MIS values.

### 1.3 Pattern-Specific Strength Score

The basic idea of the proposed approach is to estimate the pattern score by simply counting the fraction of observations belonging to the pattern under investigation. Assuming that *p*_00_ + *p*_01_ + *p*_10_ + *p*_11_ = 1 and the presence/absence threshold can be microorganism specific, the strength of each pattern can be defined as following:

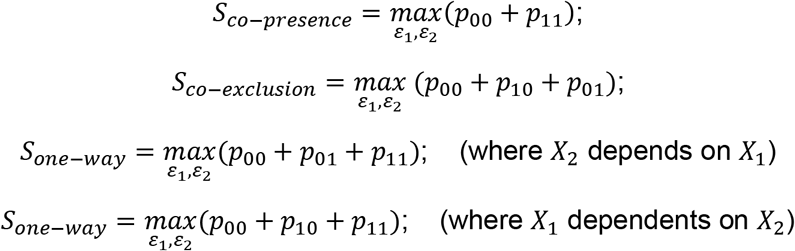

It is important to mention that the presence of *p*_00_ is required to distinguish *co-presence* patterns from cases when both organisms and simply present in all samples. *Co-presence* patterns require the existence of *co-absence* between the microorganisms in the sample set. Additionally, *co-exclusion* and *one-way* relation pattern scores include *p*_00_, because mutual absence does not contradict the pattern.

While presence/absence threshold optimization allows to consider that different microorganisms may have various minimal abundance thresholds to interact with the MC, this approach also can produce misleading results. For example, a perfect *co-presence* score may be achieved by increasing the presence/absence threshold to the point where all the observations will be counted as absent: *ɛ*_1_ ≥ (*X*_1_) and *ɛ*_2_ ≥ (*X*_2_).

This effect can be minimized by requiring a proportion of experimental observations in quadrants contributing to the pattern under consideration to be above a pre-defined minimal threshold (*m*):

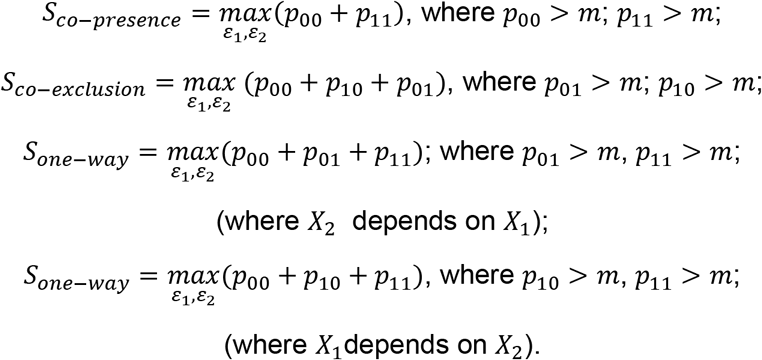

### 1.4 Non-trivial multidimensional patterns

The proposed approach can be further extended to identify more complex multidimensional patterns. For example, in some 3D patterns, the presence or absence of one organism may define what kind of 2D pattern will be exhibited between two other organisms. Figure 1f shows a case when organisms two and three will be *co-present* if organism one is present and *co-exclude* if this organism is absent):

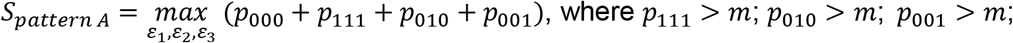

Similar to 2D patterns, not all combinations of *p*_*ijk*_ values can be interpreted as possible relations between microorganisms. Some 3D patterns can be the direct result of 3 pairwise 2D patterns: for example, the pairwise *co-exclusion* pattern between three organisms will unambiguously lead to a 3D *co-exclusion* pattern (Figure 1d):

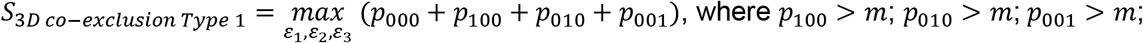

Is direct consequence of its 2D patterns:

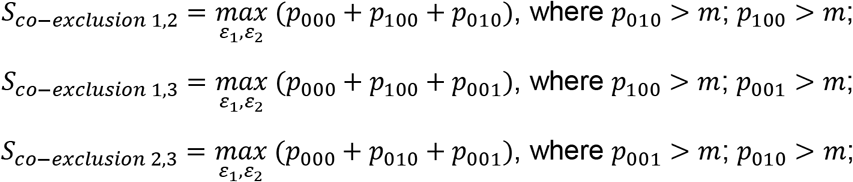

Three-dimensional *co-exclusion* patterns may additionally be observed in a very different way where each pair if organisms *co-present* only if the third one is absent (Figure 1e):

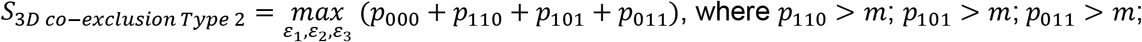

In fact, every 2D pattern has at least one non-trivial 3D analog which can be interpreted as the relation between organisms and not derived directly from any 2D combination:

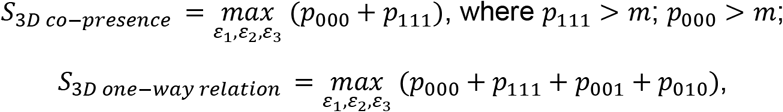

where *p*_110_ > *m*; *p*_001_ > *m*; *p*_010_ > *m*; (organism 1 requires two others to be present)

Additionally, some 3D patterns can reflect interesting new relations which exist only in higher dimensions. Figure 1f presents a case where two microorganisms (*X*_1_ and *X*_2_) follow *co-exclusion* patterns in presence of the third one (*X*_3_), as well as *co-presence* in its absence; Figure 1g shows a case where three organisms can be present only all together or one-by-one.

### 1.5 Statistical significance and Type 1 error

It is important to keep in mind that arbitrary choice of the presence threshold (*m*) and minimal score (*S*_*min*_) above which patterns are considered to be present, can significantly affect the results of the analysis in both: number of detected patterns and their statistical significance (e.g., Type 1 error). Lowering these thresholds increases chances for patterns to appear randomly, this can be detected by comparing the results produced by real data against a randomized (shuffled) dataset. Table 1 provides an example of the number of two-dimensional *one-way* relation patterns identified in original and shuffled (and renormalized) datasets from the Human Microbiome Project (Genus level, Mid vagina samples) (Consortium, 2012; Consortium, 2012).

**Table 1.** The number of two-dimensional *one-way* relation patterns identified in shuffled and real (original) mid vagina samples.

| Population Threshold\Score | Number of Patterns in Real Data/Number of Patterns in Shuffled Data |  |  |  |  |  |  |  |  |  |  |
| --- | --- | --- | --- | --- | --- | --- | --- | --- | --- | --- | --- |
|  | 0.90 | 0.91 | 0.92 | 0.93 | 0.94 | 0.95 | 0.96 | 0.97 | 0.98 | 0.99 | 1 |
| 0.05 | 268/166 | 257/144 | 234/124 | 234/124 | 220/110 | 189/78 | 166/59 | 135/35 | 98/19 | 57/10 | 57/10 |
| 0.075 | 149/63 | 140/55 | 120/46 | 120/46 | 104/35 | 86/26 | 75/20 | 62/13 | 48/4 | 30/2 | 30/2 |
| 0.1 | 83/31 | 78/25 | 68/21 | 68/21 | 65/18 | 51/10 | 41/8 | 35/4 | 23/2 | 15/0 | 15/0 |
| 0.125 | 54/13 | 50/10 | 47/9 | 47/9 | 43/6 | 36/4 | 31/1 | 24/1 | 18/1 | 13/0 | 13/0 |
| 0.15 | 34/8 | 31/5 | 27/3 | 27/3 | 26/1 | 21/1 | 18/1 | 14/1 | 12/1 | 11/0 | 11/0 |
| 0.175 | 18/7 | 16/2 | 16/1 | 16/1 | 15/1 | 10/0 | 10/0 | 8/0 | 6/0 | 3/0 | 3/0 |
| 0.2 | 13/3 | 12/1 | 12/1 | 12/1 | 12/0 | 9/0 | 7/0 | 7/0 | 6/0 | 2/0 | 2/0 |
| 0.225 | 7/1 | 7/0 | 7/0 | 7/0 | 6/0 | 5/0 | 4/0 | 4/0 | 4/0 | 2/0 | 2/0 |
| 0.25 | 5/0 | 4/0 | 4/0 | 4/0 | 4/0 | 4/0 | 4/0 | 4/0 | 4/0 | 2/0 | 2/0 |
| 0.275 | 4/0 | 4/0 | 4/0 | 4/0 | 4/0 | 4/0 | 4/0 | 4/0 | 4/0 | 2/0 | 2/0 |
| 0.3 | 4/0 | 4/0 | 4/0 | 4/0 | 4/0 | 4/0 | 4/0 | 4/0 | 4/0 | 1/0 | 1/0 |
| 0.325 | 1/0 | 1/0 | 1/0 | 1/0 | 1/0 | 1/0 | 1/0 | 1/0 | 1/0 | 0/0 | 0/0 |
| 0.35 | 0/0 | 0/0 | 0/0 | 0/0 | 0/0 | 0/0 | 0/0 | 0/0 | 0/0 | 0/0 | 0/0 |

The choice of the shuffling method reflects the underlying assumption about what would be considered as the random alternative to the observed dataset (Zero Model) (Faust and Raes, 2016; Friedman and Alm, 2012). For example, the shuffling of the abundance values across the whole dataset reflects the assumption of the total randomness of the appearances of all the values across samples and organisms. While this model preserves the overall distribution of the abundance values, it does not take into consideration that some organisms may always be present in low abundance and others can become highly dominated species in the community. In order to reflect this property on microbial abundance data shuffling across individual OTU profiles has been implemented in the presented software and used in all the examples shown in this manuscript.

The shuffling approach has been implemented as part of all pattern specific computational pipelines (see Materials and Methods section) to make sure that the search for the patterns in real data is performed only for the presence (*m*) and minimal score (*S*_*min*_) thresholds for which the number of specific patterns in shuffled data is equal to zero. The presented software however allows a variety of modifications including less strict Type 1 error requirements. The next versions of the software will include the ability to perform multiple shuffling types as well as the ability to perform shuffling multiple times.

## 2. Materials and Methods

### 2.1 Implementation

The presented software is able to identify three types of 2D patterns (*co-presence*, *co-exclusion,* and *one-way* relations) as well as three types of 3D patterns shown on Figure 1e-g, and was implemented in C++ using object-oriented approach. Executable files and source code are available in the Supplementary Materials section.

In order to improve performance in the proposed implementation, the patterns in shuffled data for all the combinations of presence threshold (*m*) and minimal score (*S*_*min*_) are calculated during the first step of the analysis so search for patterns in real data can be performed in a limited search space where zero patterns are detected in randomized (shuffled).

To evaluate performance, the presented source code was compiled using a GCC compiler version 6.3.1 under Linux CentOS 6.7. Sixteen HMP OTU files were used for the identification of 2D and 3D patterns on 4x AMD Opteron 8 core processors, 512 GB RAM, and 30 TB of storage system. Search for the 2-dimensional *co-presence*, *co-exclusion*, and *one-way relation* patterns for all tested samples took between one to three minutes, and the memory footprint did not exceed 50 megabytes of RAM. However, search for the 3D patterns may take 100’s of hours and requires a higher level of parallelization and a high-performance computing environment.

### 2.2 Data acquisition

The microbial community compositions used for this analysis originated from the NIH Human Microbiome Project (Consortium, 2012; Consortium, 2012) and contained 18 datasets associated with 16 body sites. Microbial profiles for 2910 samples have been downloaded from the project website as of December 2016 in text format (HMQCP–QIIME community Profiling v13 OTU table). Samples representing significantly low (less than 2,000) and significantly high (over 50,000) number of sequencing reads were excluded from the analysis. The microbial profiles of the remaining 2,380 samples, varying from 67 for Posterior Fornix to 200 for Antecubital Fossa, have been normalized against the total number of reads in each sample and transformed into relative abundance profiles merged to Genus taxonomy level for each body site resulting in 619 profiles. Analysis has been performed for each body site individually. For each body site under consideration, micro-organism profiles present in less than 5% of samples have been excluded from the analysis.

## 3. Results

All three types of 2D patterns have been identified in virtually every type of sample of the Human Microbiome Project data (see Supplementary Materials document). The largest number of patterns (all types included) has been detected in Supragingival plaque, Tongue dorsum, Stool, and Subgingival plaque datasets (see Figure 3a-d and Supplementary Materials document). No apparent correlation has been observed between the number of patterns and the total number of samples nor the number of OTUs in the datasets. It is important to mention that while all of the observed 2D patterns pass statistical significance criteria, the overall size and complexity of the resulting networks depends on the pattern score threshold (Figure 3e). Some 3D patterns have also been observed in Buccal mucosa, Supragingival plaque, and Merged Retroauricular crease datasets (see example in Table 2).

**Figure 3.**
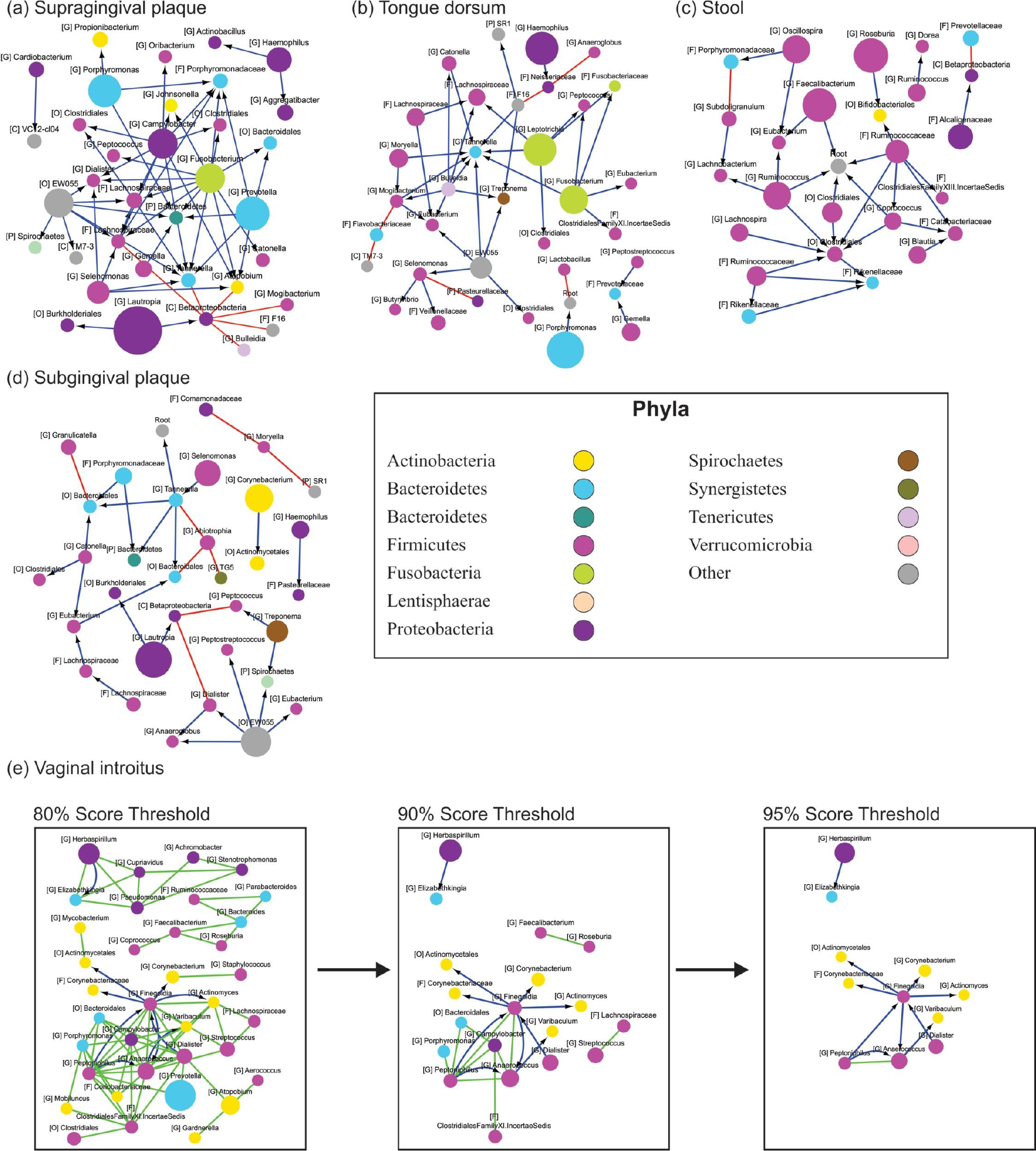
2D networks for (a) Supragingival plaque, (b) Tongue dorsum, (c) Stool, (d) Subgingival plaque and (e) Vaginal introitus samples at the genus level. Example of the effects of the patterns’ score threshold on the network’s complexity(e). Node colors reflect different taxonomy assignments at Phylum level and node sizes are proportional to the average relative abundance of the microorganism across samples. Capital letters inside square brackets represent the lowest taxonomy level identified for each OTU: G (Genus), F (Family), O (Order), C (Class) and P (Phyla). The color of edges indicates relationship type: blue with a black arrow (*one-way* relations), red (*co-exclusion*), and light green (*co-presence*).

**Table 2.** Example of 3D patterns where organisms 2 and 3 are *co-present* if organism 1 is present and *co-excluded* if Organism 1 is absent identified in Anterior nares samples (genus level). Calculations have been performed with the minimum population threshold set to 0.1.

| Organism 1 | Organism 2 | Organism 3 | Pattern Score | p<sub>000</sub> | p<sub>001</sub> | p<sub>010</sub> | p<sub>011</sub> | p<sub>100</sub> | p<sub>101</sub> | p<sub>110</sub> | p<sub>111</sub> |
| --- | --- | --- | --- | --- | --- | --- | --- | --- | --- | --- | --- |
| [G] Actinomyces | [G] Rothia | [G] Neisseria | 0.94 | 0.48 | 0.12 | 0.18 | 0.05 | 0.00 | 0.00 | 0.02 | 0.16 |
| [F] Ruminococcaceae | [G] Bacteroides | [F] Lachnospiraceae | 0.94 | 0.54 | 0.12 | 0.17 | 0.03 | 0.02 | 0.00 | 0.00 | 0.11 |
| [G] Faecalibacterium | [G] Bacteroides | [F] Lachnospiraceae | 0.94 | 0.57 | 0.12 | 0.16 | 0.04 | 0.00 | 0.00 | 0.02 | 0.10 |
| [G] Oscillospira | [G] Bacteroides | [F] Lachnospiraceae | 0.94 | 0.60 | 0.12 | 0.12 | 0.03 | 0.01 | 0.01 | 0.02 | 0.10 |
| [G] Oscillospira | [F] Lachnospiraceae | [G] Faecalibacterium | 0.94 | 0.57 | 0.10 | 0.17 | 0.04 | 0.01 | 0.02 | 0.00 | 0.10 |
| [G] Neisseria | [G] Rothia | [G] Prevotella | 0.94 | 0.56 | 0.15 | 0.12 | 0.02 | 0.01 | 0.01 | 0.02 | 0.11 |
| [G] Neisseria | [G] Rothia | [G] Veillonella | 0.94 | 0.48 | 0.23 | 0.10 | 0.04 | 0.01 | 0.01 | 0.00 | 0.13 |

## 4. Discussion

Identification of interaction patterns in microbial communities is essential to further our understanding of relationships in microbial communities. Knowledge of the interactions between specific organisms can help transition microbiomes between enterotypes and better predict microbial responses due to perturbations (e.g., targeted antimicrobials, probiotics, prebiotics). Ability to manipulate microbial communities in terms of community members and their functions will open new opportunities for precision medicine and personalized treatments. Thus, the development of systematic and statistically-sound methods for interaction pattern identification is a necessary step to understand the structure of microbiomes and the processes by which they evolve.

Pattern visualization involving multiple organisms, however, remains a significant challenge. Traditionally, the graph (network) representation of the patterns between organisms in microbial communities represents each OTU as the node and pairwise relationships as edges (Deng, et al., 2012; Faust and Raes, 2016; Friedman and Alm, 2012; Reshef, et al., 2011). We believe that one of the possible ways to visualize 2D, 3D, and higher dimensional patterns could be by using a Multi-Layer Network (Multi-Layer Graph) which in contrast with traditional graphs (networks) can simultaneously include nodes of different types (representing both OTUs and patterns). Figure 4 shows an example of such a representation for two- and three-dimensional patterns in Attached keratinized gingiva samples from the Human Microbiome Project.

**Figure 4.**
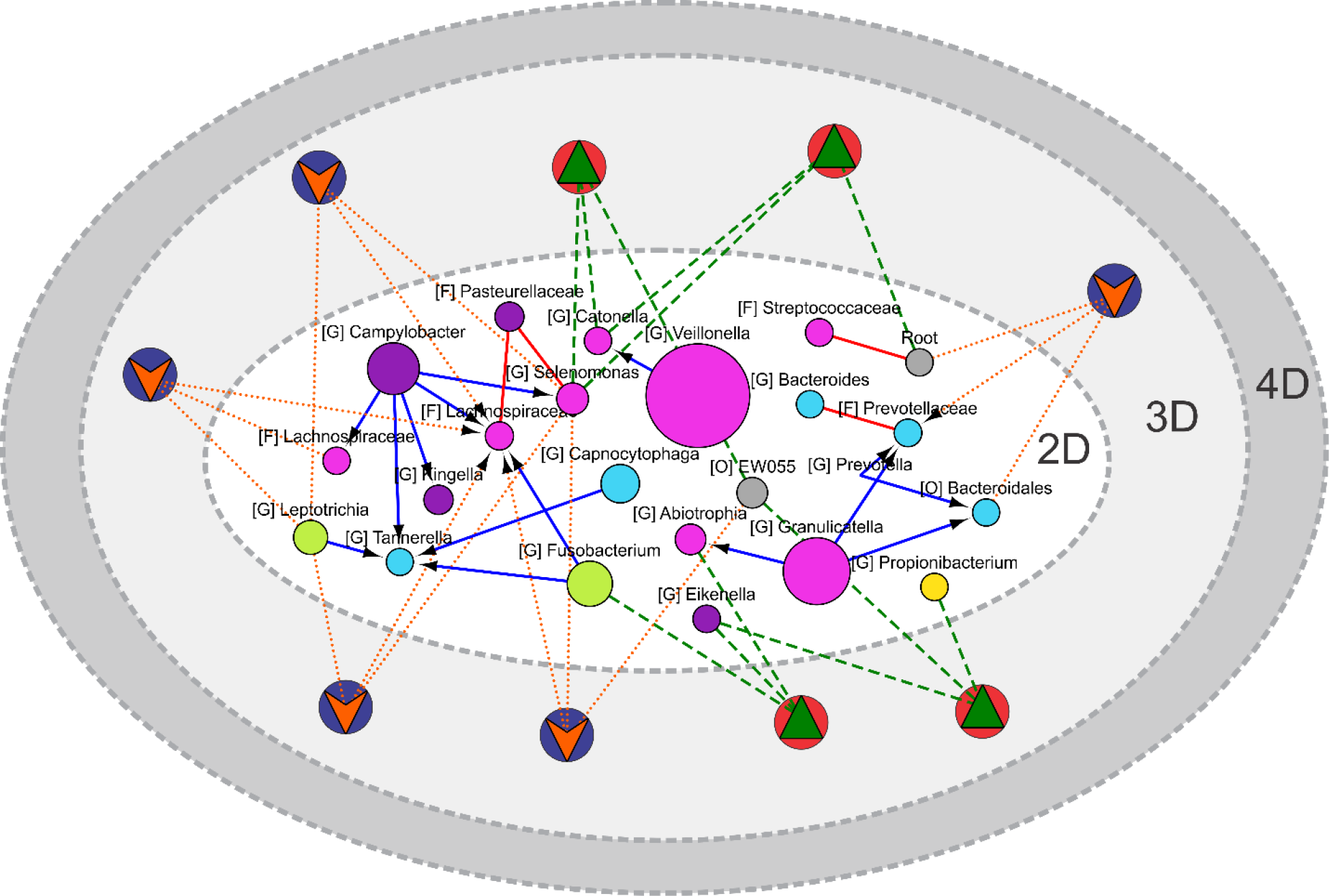
Example of Multi-Layer Network simultaneous visualization of two- and three-dimensional patterns in Attached keratinized gingiva samples. The network contains two types of nodes representing OTUs (circular) and three-dimensional patterns (circular with a triangle). Node colors reflect different taxonomy assignments at Phylum level and node sizes are proportional to the average relative abundance of the microorganism across samples. Capital letters inside square brackets represent the lowest taxonomy level identified for each OTU: G (Genus), F (Family), O (Order), C (Class) and P (Phyla). The color of edges indicates relationship type: blue with a black arrow (*one-way* relations), red (*co-exclusion*), light green (*co-presence*), dark green (3D *co-presence*), and orange (Type 2 *co-exclusion*).

Future work will be focused on expanding the presented approach to be applied beyond microbiome abundance tables to include physical (pH, temperature, oxygen concentration) and biochemical (antibiotics, nutrients and metabolites concentrations) variables. It can also be extended to the simultaneous analysis of multi-omics data, such as protein and mRNA expression in both microbial communities and mammalian host.

