## Supplementary material for "Identification of Complex Multidimensional Patterns in Microbial Communities": Pickle User Manual.pdf

### Pickle 1.0

#### User Manual

February 2019

Software presented in Identification of Complex Multidimensional Patterns in Microbial Communities

#### 1. Introduction

**Pickle** is the software designed to search for two- and three-dimensional patterns in tabulated OTU data files. There are four pipelines available for the specific pattern identification:

1. 2D_patterns_main
   Performs search for two dimensional co-exclusion, co-presence, and one-way relation patterns
2. 3D_All_together_or_alone
   Perform search for three dimensional all together or alone pattern
3. 3D_Switch_Pattern
   Perform search for three dimensional X23 present if X1 present pattern
4. 3D_Type2_Coexclusion
   Perform search for three dimensional type2 co-exclusion pattern

Questions, suggestions, and concerns related to the performance and functionality of the software should be addressed via

#### 2. Compile Pickle

In order to compile C++ source code please use the following command:

g++ -O3 2D_Patterns_Main.cpp –w –o 2D_Patterns

g++ -O3 3D_All_Together_Or_Alone.cpp –w –o 3D_All_Together_Or_Alone

g++ -O3 3D_Switch_Pattern.cpp –w –o 3D_Switch_Pattern

g++ -O3 3D_Type2_Coexclusion.cpp –w –o 3D_Type2_Coexclusion

*Note:* it is possible to use any other C++ compiler

#### 3. Run Pickle

There are three groups of parameters (nine values total) which can affect results of the search for both 2D and 3D patterns and can be defined using the command line.

Presence threshold optimization parameters:
-n min_presence_threshold (default value 0.0);
-m max_presence_threshold (default value 0.005 or 0.5%);
-s threshold_presence_step (default value 0.0001 or 0.01%).
These parameters are used to identify if the organism considered to be present or absent in the sample.

The population threshold optimization parameters are defining the minimal fraction of experimental observations required to be present in each of quarters (in multidimensional cases sectors) defining each particular pattern:
-p min_population_threshold (default value 0.1 or 10%);
-t max_population_threshold (default value 0.2 or 20%);
-r population_threshold_step (default value 0.01 or 1%).

The patterns score threshold parameters are used to filter which combination of the of the population threshold/pattern score will be presented in the output file.
-z min_score (default value 0.9 or 90%);
-x max_score (default value 1 or 100%);
-c score_step (default value 0.01 or 1%).

It is important to mention that while the presence and population threshold parameters are contributing to the performance of the application (time complexity), the patterns score threshold parameters affect only the total number of the identified patterns and as result only affect output file size.

**Command line example**

To run any of the 2D or 3D Pickle pipelines in a Linux environment, the user must provide the command line with 2 parameters separated by a single space:

Example of command with default parameters: 2D_patterns_main -i ./Anterior_nares_genus_example.txt -o ./Anterior_nares_genus_manual_

Example of full command: 2D_patterns_main -i ./Anterior_nares_genus_example.txt -o ./Anterior_nares_genus_manual_ -n 0.0 -m 0.005 -s 0.0001 -p 0.1 -t 0.2 -r 0.01 -z 0.9 -x 1 -c 0.001

#### 5. Input File Format

Input file for each of the available pattern search pipelines is a tabulated OTU file. The first line of the input file is a header with a list of samples separated by tab. Starting from the second line, file should contain OTU name and corresponding normalized values separated by tabs. Each line should end with taxonomical lineage associated to the OTU.
