## Supplementary material for "Identification of Complex Multidimensional Patterns in Microbial Communities": 2D_Patterns_Main.cpp

```
/*
THE SOFTWARE IS PROVIDED "AS IS", WITHOUT WARRANTY OF ANY KIND, EXPRESS OR IMPLIED, INCLUDING BUT NOT LIMITED TO THE WARRANTIES OF
MERCHANTABILITY, FITNESS FOR A PARTICULAR PURPOSE AND NONINFRINGEMENT. IN NO EVENT SHALL THE AUTHORS OR COPYRIGHT HOLDERS BE LIABLE FOR
ANY CLAIM, DAMAGES OR OTHER LIABILITY, WHETHER IN AN ACTION OF CONTRACT, TORT OR OTHERWISE, ARISING FROM, OUT OF OR IN CONNECTION WITH
THE SOFTWARE OR THE USE OR OTHER DEALINGS IN THE SOFTWARE.
*/
# include <iostream>
# include <fstream>


#include "Shuffle.h"
#include "Stat_and_Sort.h"

#include "OTU_Table.h"
#include "OTU_Profile.h"

#include "Two_D_Pattern.h"
#include "Array_of_Two_D_Patterns.h"

#include "Three_D_Pattern.h"
#include "Array_of_Three_D_Patterns.h"


#include "Two_D_Patterns_Pipelines.h"
#include "Three_D_Patterns_Pipelines.h"


using namespace std;

//2D Patterns Seeking Pipeline

int main(int argc, char **argv)

{


//==================== PARAMETERS
/*
 *
-i  input file name
-o  prefix for the output files

Presence threshold optimization parameters:
-n 	min_presence_threshold (default value 0.0);
-m	max_presence_threshold (default value 0.005 or 0.5%);
-s	threshold_presence_step (default value 0.0001 or 0.01%).
These parameters are used to identify if organism considered to be present or absent in the sample.

The population threshold optimization parameters define the minimal fraction of experimental observations required to be present in each of quartiles (in multidimensional cases sectors) defining each particular pattern:
-p 	min_population_threshold (default value 0.1 or 10%);
-t 	max_population_threshold (default value 0.2 or 20%);
-r 	population_threshold_step (default value 0.01 or 1%).


The patterns score threshold parameters are used to filter which combination of the population threshold/pattern score will be presented in the output file.
-z 	min_score (default value 0.9 or 90%);
-x 	max_score (default value 1 or 100%);
-c 	score_step (default value 0.01 or 1%).


*/

    double min_presence_threshold = 0.;	// these parameters will affect the time of calculations
	double max_presence_threshold = 0.005;
	double threshold_presence_step = 0.0001;

    double min_population_threshold = 0.1; // these parameters will affect the time of calculations
	double max_population_threshold = 0.2;
	double population_threshold_step = 0.01;

   double min_score = 0.9; // Does not affects performance. min_score affect changes how many patterns will be identified, lower threshold results in more weaker patterns
    double max_score = 1.;
	double score_step = 0.01;

    string output_prefix="";
    string OTU_file_name ="";

    if(argc<2) 
    {
    	cout<<"Please try again using following parameters:"<<endl;
    	cout<<"*******************************************"<<endl;
    	cout<<"Mandatory parameters:"<<endl;
    	cout<<"-i  input file name"<<endl;
    	cout<<"-o  prefix for the output files"<<endl;
    	cout<<"*******************************************"<<endl;
    	cout<<"Optional parameters:"<<endl;
		cout<<"Presence threshold optimization parameters:"<<endl;
		cout<<"-n 	min_presence_threshold (default value 0.0);"<<endl;
		cout<<"-m	max_presence_threshold (default value 0.005 or 0.5%);"<<endl;
		cout<<"-s	threshold_presence_step (default value 0.0001 or 0.01%)."<<endl;
		cout<<"These parameters are used to identify if organism considered to be present or absent in the sample."<<endl;
		cout<<"*******************************************"<<endl;
		cout<<"The population threshold optimization parameters define the minimal fraction of experimental observations required to be present in each of quartiles (in multidimensional cases sectors) defining each particular pattern:"<<endl;
		cout<<"-p 	min_population_threshold (default value 0.1 or 10%);"<<endl;
		cout<<"-t 	max_population_threshold (default value 0.2 or 20%);"<<endl;
		cout<<"-r 	population_threshold_step (default value 0.01 or 1%)."<<endl;
		cout<<"*******************************************"<<endl;
		cout<<"The patterns score threshold parameters are used to filter which combination of the population threshold/pattern score will be presented in the output file."<<endl;
		cout<<"-z 	min_score (default value 0.9 or 90%);"<<endl;
		cout<<"-x 	max_score (default value 1 or 100%);"<<endl;
		cout<<"-c 	score_step (default value 0.01 or 1%)."<<endl;

    	return 1;

    }


//========Parsing"<< input parameters"<<
    for (unsigned int argn= 1; argn < argc-1 ; argn++) 
    {

    	if(argv[argn][0] != '-') continue; 

        switch (argv[argn][1]) 
        {
        case 'n': min_presence_threshold = atof(argv[argn+1]); break;
        case 'm': max_presence_threshold= atof(argv[argn+1]); break;
        case 's': threshold_presence_step= atof(argv[argn+1]); break;
        case 'p': min_population_threshold= atof(argv[argn+1]); break;
        case 't': max_population_threshold= atof(argv[argn+1]); break;
        case 'r': population_threshold_step= atof(argv[argn+1]); break;
        case 'z': min_score= atof(argv[argn+1]); break;
        case 'x': max_score= atof(argv[argn+1]); break;       
        case 'c': score_step= atof(argv[argn+1]); break;
        case 'o': output_prefix=argv[argn+1];  break;
        case 'i': OTU_file_name=argv[argn+1]; break;

        default:
           cout<<"Parameter that you enered: -"<<argv[argn]<<" was not recognized!" <<endl;
            exit(EXIT_FAILURE);
        }
    }


//========Show List of The Parameters for Execution
    cout<<"Pickle performs search for two dimensional patterns using following the parameters:"<<endl;

    cout<<"Output files prefix: "<< output_prefix<<endl;
    cout<<"Input file: "<< OTU_file_name<<endl;

    cout<<"Minimum presence threshold: "<< min_presence_threshold <<endl;	// these parameters will affect the time of calculations
    cout<<"Maximum presence threshold: "<< max_presence_threshold <<endl;
    cout<<"Presence threshold step: "<< threshold_presence_step<<endl;

    cout<<"Minimum population threshold: "<< min_population_threshold<<endl; // these parameters will affect the time of calculations
    cout<<"Maximum population threshold: "<< max_population_threshold<<endl;
    cout<<"Population threshold step: "<< population_threshold_step<<endl;

    cout<<"Minimum score: "<< min_score<<endl;
    cout<<"Maximum score: "<< max_score<<endl;
    cout<<"Score step: "<< score_step<<endl;


//Pipelines


    string out_file_name_co_presence =output_prefix;
    out_file_name_co_presence=out_file_name_co_presence+"_co_presence_short.txt";
    string out_file_name_co_presence_long =output_prefix;
    out_file_name_co_presence_long=out_file_name_co_presence_long+"_co_presence_long.txt";

//Co-Presence
	Two_D_Patterns_Pipelines::find_2D_Co_Presence_patterns(&OTU_file_name[0], &out_file_name_co_presence[0], &out_file_name_co_presence_long[0],
														min_presence_threshold, max_presence_threshold, threshold_presence_step,
														min_population_threshold, max_population_threshold, population_threshold_step,
														min_score, max_score, score_step);

//Co-Exclusion

    string out_file_name_co_exclusion = output_prefix;
    out_file_name_co_exclusion =out_file_name_co_exclusion +"_co_exclusion_short.txt";
    string out_file_name_co_exclusion_long = output_prefix;
    out_file_name_co_exclusion_long= out_file_name_co_exclusion_long +"_co_exclusion_long.txt";

	Two_D_Patterns_Pipelines::find_2D_Co_Exclusion_patterns(&OTU_file_name[0], &out_file_name_co_exclusion[0], &out_file_name_co_exclusion_long[0],
														min_presence_threshold, max_presence_threshold, threshold_presence_step,
														min_population_threshold, max_population_threshold, population_threshold_step,
														min_score, max_score, score_step);

//One-way relationship
    string out_file_name_One_way = output_prefix;
    out_file_name_One_way =out_file_name_One_way +"_one_way_short.txt";
    string out_file_name_One_way_long = output_prefix;
    out_file_name_One_way_long =out_file_name_One_way_long+"_one_way_long.txt";

	Two_D_Patterns_Pipelines::find_2D_One_Way_Relation_patterns(&OTU_file_name[0], &out_file_name_One_way[0], &out_file_name_One_way_long[0],
																min_presence_threshold, max_presence_threshold, threshold_presence_step,
																min_population_threshold, max_population_threshold, population_threshold_step,
																min_score, max_score, score_step);


return 1;
}


//====================================================================================================================
//====================================================================================================================
//====================================================================================================================
```
