## Supplementary material for "Identification of Complex Multidimensional Patterns in Microbial Communities": Array_of_Three_D_Patterns.h

```
/*
THE SOFTWARE IS PROVIDED "AS IS", WITHOUT WARRANTY OF ANY KIND, EXPRESS OR IMPLIED, INCLUDING BUT NOT LIMITED TO THE WARRANTIES OF
MERCHANTABILITY, FITNESS FOR A PARTICULAR PURPOSE AND NONINFRINGEMENT. IN NO EVENT SHALL THE AUTHORS OR COPYRIGHT HOLDERS BE LIABLE FOR
ANY CLAIM, DAMAGES OR OTHER LIABILITY, WHETHER IN AN ACTION OF CONTRACT, TORT OR OTHERWISE, ARISING FROM, OUT OF OR IN CONNECTION WITH
THE SOFTWARE OR THE USE OR OTHER DEALINGS IN THE SOFTWARE.
*/
#ifndef _ARRAY_OF_THREE_D_PATTERNS_H_
#define _ARRAY_OF_THREE_D_PATTERNS_H_


#include <iostream>
#include <fstream>
#include <stdlib.h>
#include <random>
#include <time.h>

#include "Three_D_Pattern.h"

using namespace std;

class Array_of_Three_D_Patterns
{

public: // ------------------- For now...

	unsigned int max_number_of_patterns;	
	unsigned int number_of_patterns;

	Three_D_Pattern ** pattern;

	Array_of_Three_D_Patterns(unsigned int _max_number_of_patterns=100000, ostream & err_msg_out = cout);	// 10,000
	~Array_of_Three_D_Patterns();

// NOTE: compatison is besd only on OTU_Index_1 and OTU_Index_1 (order specific!)
	bool is_Pattern_Present(Three_D_Pattern * TD_p, ostream & err_msg_out = cout);	


// NOTE: No Pattern copy will be created! The pointer to the pattern will be added to the array.
	bool add_Pattern(Three_D_Pattern * TD_p, ostream & err_msg_out = cout);

	void show_Statistics();
	void show_Statistics(ofstream & out_f);

	void show_All();
	void show_All(ofstream & out_f);


};
//-------------------------------------------------------------------------------------------------------------
//-------------------------------------------------------------------------------------------------------------
void Array_of_Three_D_Patterns::show_Statistics()
{
	cout << "\n============== Array_of_Three_D_Patterns::show_Statistics ========== (START)\n";

	cout<<"number_of_patterns="<<number_of_patterns<<endl;

	unsigned int i;
	for (i = 0; i < number_of_patterns; i++)
	{
		cout << "OTU_index1=" << pattern[i]->OTU_Index_1 << "\t";
		cout << "OTU_index2=" << pattern[i]->OTU_Index_2 << "\t";
		cout << "OTU_index3=" << pattern[i]->OTU_Index_3 << "\t";

		cout << "score=" << pattern[i]->score << "\t";
		cout << "pupulation threshold=" << pattern[i]->population_threshold << "\t";
		cout << "threshold_x1=" << pattern[i]->threshold_x1 << "\t";
		cout << "threshold_x2=" << pattern[i]->threshold_x2 << endl;
		cout << "threshold_x3=" << pattern[i]->threshold_x3 << endl;
	}

	cout << "\n============== Array_of_Three_D_Patterns::show_Statistics ========== (END)\n";

}
//-------------------------------------------------------------------------------------------------------------

void Array_of_Three_D_Patterns::show_Statistics(ofstream & out_f)
{
	out_f << "number_of_patterns=" << number_of_patterns << endl;

	out_f << "OTU 1\t taxonomy\tOTU 2\ttaxonomy\tOTU 3\ttaxonomy\tobserved_population_threshold\tscore\tthr_x1\tthr_x2\tthr_x3\tp_000\tp_001\tp_010\tp_011\tp_100\tp_101\tp_110\tp_111" << endl;

	unsigned int i;
	for (i = 0; i < number_of_patterns; i++)
	{
	
		out_f <<       pattern[i]->OTU_Index_1 << pattern[i]->OTU_p_1->taxonomy_text
			<< "\t" << pattern[i]->OTU_Index_2 << pattern[i]->OTU_p_2->taxonomy_text
			<< "\t" << pattern[i]->OTU_Index_3 << pattern[i]->OTU_p_3->taxonomy_text
			<< "\t" << pattern[i]->population_threshold << "\t" 
			<< pattern[i]->score << "\t" << pattern[i]->threshold_x1 << "\t" << pattern[i]->threshold_x2 << "\t" << pattern[i]->threshold_x3 << "\t"
			<< pattern[i]->p_000 << "\t" << pattern[i]->p_001 << "\t" << pattern[i]->p_010 << "\t" << pattern[i]->p_011 << "\t"
			<< pattern[i]->p_100 << "\t" << pattern[i]->p_101 << "\t" << pattern[i]->p_110 << "\t" << pattern[i]->p_111 << endl;
	}
}
//-------------------------------------------------------------------------------------------------------------

void Array_of_Three_D_Patterns::show_All(ofstream & out_f)
{
	out_f << "number_of_patterns=" << number_of_patterns << endl;

	unsigned int i;
	unsigned int ii;
	for (i = 0; i < number_of_patterns; i++)
	{
		out_f << "\n==============================================================\n";

		out_f << "OTU 1\t taxonomy\tOTU 2\ttaxonomy\tOTU 3\ttaxonomy\tobserved_population_threshold\tscore\tthr_x1\tthr_x2\tthr_x3\tp_000\tp_001\tp_010\tp_011\tp_100\tp_101\tp_110\tp_111" << endl;

		out_f << pattern[i]->OTU_Index_1 << pattern[i]->OTU_p_1->taxonomy_text
			<< "\t" << pattern[i]->OTU_Index_2 << pattern[i]->OTU_p_2->taxonomy_text
			<< "\t" << pattern[i]->OTU_Index_3 << pattern[i]->OTU_p_3->taxonomy_text
			<< "\t" << pattern[i]->population_threshold << "\t" << pattern[i]->score << "\t" 
			<< pattern[i]->threshold_x1 << "\t" << pattern[i]->threshold_x2 << "\t" << pattern[i]->threshold_x3 << "\t"
			<< pattern[i]->p_000 << "\t" << pattern[i]->p_001 << "\t" << pattern[i]->p_010 << "\t" << pattern[i]->p_011 << "\t"
			<< pattern[i]->p_100 << "\t" << pattern[i]->p_101 << "\t" << pattern[i]->p_110 << "\t" << pattern[i]->p_111 << endl;

		out_f << "OTU 1\t";
		for (ii = 0; ii < pattern[i]->OTU_p_1->number_of_samples; ii++) out_f << pattern[i]->OTU_p_1->abundance_value[ii] << "\t";
		out_f << endl;

		out_f << "OTU 2\t";
		for (ii = 0; ii < pattern[i]->OTU_p_2->number_of_samples; ii++) out_f << pattern[i]->OTU_p_2->abundance_value[ii] << "\t";
		out_f << endl;

		out_f << "OTU 3\t";
		for (ii = 0; ii < pattern[i]->OTU_p_3->number_of_samples; ii++) out_f << pattern[i]->OTU_p_3->abundance_value[ii] << "\t";
		out_f << endl << endl;
	}
}
//-------------------------------------------------------------------------------------------------------------


//-------------------------------------------------------------------------------------------------------------

Array_of_Three_D_Patterns::Array_of_Three_D_Patterns(unsigned int _max_number_of_patterns, ostream & err_msg_out)
{
	number_of_patterns=0;
	
	max_number_of_patterns=_max_number_of_patterns;

	pattern = new Three_D_Pattern *[max_number_of_patterns];

	unsigned int i;
	for (i = 0; i < max_number_of_patterns; i++) pattern[i] = NULL;
}
//-------------------------------------------------------------------------------------------------------------

Array_of_Three_D_Patterns::~Array_of_Three_D_Patterns()
{
	unsigned int i;
	for (i = 0; i < max_number_of_patterns; i++) if(pattern[i] != NULL) delete pattern[i];
	delete [] pattern;
}
//-------------------------------------------------------------------------------------------------------------

bool Array_of_Three_D_Patterns::is_Pattern_Present(Three_D_Pattern * TD_p, ostream & err_msg_out)
{
	unsigned int i;
	for (i = 0; i < number_of_patterns; i++)
	{
		if (pattern[i]->OTU_Index_1 != TD_p->OTU_Index_1) continue;
		if (pattern[i]->OTU_Index_2 != TD_p->OTU_Index_2) continue;
		if (pattern[i]->OTU_Index_3 != TD_p->OTU_Index_3) continue;

		if (pattern[i]->population_threshold != TD_p->population_threshold) continue;
		if (pattern[i]->score != TD_p->score) continue;
		return true;
	}
		
	return false;
}
//-------------------------------------------------------------------------------------------------------------

bool Array_of_Three_D_Patterns::add_Pattern(Three_D_Pattern * TD_p, ostream & err_msg_out)
{
	if (number_of_patterns >= max_number_of_patterns) return false;
	
	pattern[number_of_patterns] = TD_p;
	number_of_patterns++;

	return true;
}
//-------------------------------------------------------------------------------------------------------------
//-------------------------------------------------------------------------------------------------------------
//-------------------------------------------------------------------------------------------------------------
#endif //Array_of_Three_D_Patterns
```
