## Supplementary material for "Identification of Complex Multidimensional Patterns in Microbial Communities": OTU_Profile.h

```
/*
THE SOFTWARE IS PROVIDED "AS IS", WITHOUT WARRANTY OF ANY KIND, EXPRESS OR IMPLIED, INCLUDING BUT NOT LIMITED TO THE WARRANTIES OF
MERCHANTABILITY, FITNESS FOR A PARTICULAR PURPOSE AND NONINFRINGEMENT. IN NO EVENT SHALL THE AUTHORS OR COPYRIGHT HOLDERS BE LIABLE FOR
ANY CLAIM, DAMAGES OR OTHER LIABILITY, WHETHER IN AN ACTION OF CONTRACT, TORT OR OTHERWISE, ARISING FROM, OUT OF OR IN CONNECTION WITH
THE SOFTWARE OR THE USE OR OTHER DEALINGS IN THE SOFTWARE.
*/
#ifndef _OTU_PROFILE_H_
#define _OTU_PROFILE_H_


#include <iostream>
#include <fstream>
#include <stdlib.h>
#include <random>
#include <time.h>

using namespace std;

class OTU_Profile
{

public:

	unsigned int number_of_samples;


	unsigned int sample_name_length; // = 200
	unsigned int OTU_short_name_length; // = 100 
	unsigned int taxonomy_text_length; // = 2000

	double * abundance_value;	// first index ==> sample, second index ==> OTU

	char * OTU_short_name;	// first index ==> sample, second index ==> char array
	char * taxonomy_text;	// first index ==> sample, second index ==> char array

// ------- derivative values

	double OTUs_min;
	double OTUs_max; 
	double OTUs_average; 

	double percent_of_samples_OTU_present;


public:

	OTU_Profile(OTU_Profile * _p);
	OTU_Profile(unsigned int _number_of_samples);

	~OTU_Profile();

	bool calculate_derivative_values(double OTU_presence_threshold=0.0, ostream & err_msg_out = cout);
	void show_Statistics(ostream & err_msg_out = cout);
	void show_Statistics(ofstream & out_f, ostream & err_msg_out = cout);
	void show_All(ostream & err_msg_out = cout);


// NOTE: The OTU_profile array has to be allocated outside 
//	bool get_OTU_profile(double * OTU_profile, unsigned int OTU_index, ostream & err_msg_out = cout);

};
//-------------------------------------------------------------------------------------------------------------
//-------------------------------------------------------------------------------------------------------------
//-------------------------------------------------------------------------------------------------------------

void OTU_Profile::show_Statistics(ostream & err_msg_out)
{

	cout << "============== OTU_Profile::show_Statistics ========== (START)\n";

	cout << OTU_short_name << "\t" << taxonomy_text << endl;
	cout << "Present in " << percent_of_samples_OTU_present<< " % of samples "
		<< "min=" << OTUs_min<< "  max=" << OTUs_max<<endl;

	cout << "============== OTU_Profile::show_Statistics ========== (END)\n";
}
//-------------------------------------------------------------------------------------------------------------

void OTU_Profile::show_Statistics(ofstream & out_f, ostream & err_msg_out)
{
	out_f << "============== OTU_Profile::show_Statistics ========== (START)\n";

	out_f << OTU_short_name << "\t" << taxonomy_text << endl;
	out_f << "Present in " << percent_of_samples_OTU_present << " % of samples "
		<< "min=" << OTUs_min << "  max=" << OTUs_max << "  Average=" << OTUs_average << endl;

	out_f << "============== OTU_Profile::show_Statistics ========== (END)\n";
}
//-------------------------------------------------------------------------------------------------------------


void OTU_Profile::show_All(ostream & err_msg_out)
{
	cout << "============== OTU_Profile::All ========== (START)\n";
	show_Statistics();

	unsigned int i;
	for (i = 0; i < number_of_samples; i++) cout << "abundance_value["<<i<<"]="<< abundance_value<< endl;

	cout << "============== OTU_Table::show_Statistics ========== (END)\n";

}
//-------------------------------------------------------------------------------------------------------------

bool OTU_Profile::calculate_derivative_values(double OTU_presence_threshold, ostream & err_msg_out)
{
	OTUs_min = abundance_value[0];
	OTUs_max = abundance_value[0];
	OTUs_average = 0.;

	unsigned int i;
	unsigned int counter=0;

	for (i = 0; i < number_of_samples; i++)
	{
		OTUs_average += abundance_value[i];
		if (OTUs_min > abundance_value[i])OTUs_min = abundance_value[i];
		if (OTUs_max < abundance_value[i])OTUs_max = abundance_value[i];
		if (abundance_value[i] > OTU_presence_threshold) counter++;
	}
	percent_of_samples_OTU_present = (double)counter / (double)number_of_samples;
	OTUs_average = OTUs_average / (double)number_of_samples;

	return true;
}
//-------------------------------------------------------------------------------------------------------------

OTU_Profile::OTU_Profile(OTU_Profile * OTU_p)
{
	number_of_samples=OTU_p->number_of_samples;

// ------- default parameters
	sample_name_length = 200;
	OTU_short_name_length = 200;
	taxonomy_text_length = 2000;

	// --------- allocations
	OTU_short_name = new char[OTU_short_name_length];
	taxonomy_text = new char[taxonomy_text_length];
	abundance_value = new double[number_of_samples];

	unsigned int i;
	for (i = 0; i < OTU_short_name_length; i++) OTU_short_name[i] = OTU_p->OTU_short_name[i];
	for (i = 0; i < number_of_samples; i++) abundance_value[i]=OTU_p->abundance_value[i];
	for (i = 0; i < taxonomy_text_length; i++) taxonomy_text[i] = OTU_p->taxonomy_text[i];
		
	calculate_derivative_values();
}
//-------------------------------------------------------------------------------------------------------------

OTU_Profile::OTU_Profile(unsigned int _number_of_samples)
{
	number_of_samples = _number_of_samples;

	// ------- default parameters
	sample_name_length = 200;
	OTU_short_name_length = 200;
	taxonomy_text_length = 2000;

	// --------- allocations
	OTU_short_name = new char[OTU_short_name_length];
	taxonomy_text = new char[taxonomy_text_length];
	abundance_value = new double[number_of_samples];

	// --------------------------- Initial values
	OTUs_min = 0.;
	OTUs_max = 0.;
	OTUs_average = 0.;

	double percent_of_samples_OTU_present=0.;

}
//-------------------------------------------------------------------------------------------------------------

OTU_Profile::~OTU_Profile()
{
	delete [] abundance_value;
	delete [] OTU_short_name;
	delete [] taxonomy_text;
}


//-------------------------------------------------------------------------------------------------------------
//-------------------------------------------------------------------------------------------------------------
//-------------------------------------------------------------------------------------------------------------
//-------------------------------------------------------------------------------------------------------------
#endif //_OTU_PROFILE_H_
```
