## Supplementary material for "Identification of Complex Multidimensional Patterns in Microbial Communities": OTU_Table.h

```
/*
THE SOFTWARE IS PROVIDED "AS IS", WITHOUT WARRANTY OF ANY KIND, EXPRESS OR IMPLIED, INCLUDING BUT NOT LIMITED TO THE WARRANTIES OF
MERCHANTABILITY, FITNESS FOR A PARTICULAR PURPOSE AND NONINFRINGEMENT. IN NO EVENT SHALL THE AUTHORS OR COPYRIGHT HOLDERS BE LIABLE FOR
ANY CLAIM, DAMAGES OR OTHER LIABILITY, WHETHER IN AN ACTION OF CONTRACT, TORT OR OTHERWISE, ARISING FROM, OUT OF OR IN CONNECTION WITH
THE SOFTWARE OR THE USE OR OTHER DEALINGS IN THE SOFTWARE.
*/
#ifndef _OTU_TABLE_H_
#define _OTU_TABLE_H_


#include <iostream>
#include <fstream>
#include <stdlib.h>
#include <random>
#include <time.h>

#include "OTU_Profile.h"

using namespace std;

class OTU_Table
{

public: // ------------------- For now...

	unsigned int number_of_samples;
	unsigned int number_of_OTUs;

	unsigned int sample_name_length; // = 200
	unsigned int OTU_short_name_length; // = 200 
	unsigned int taxonomy_text_length; // = 2000

	double ** abundance_value;	// first index ==> sample, second index ==> OTU

	char ** sample_name;	// first index ==> sample, second index ==> char array
	char ** OTU_short_name;	// first index ==> sample, second index ==> char array
	char ** taxonomy_text;	// first index ==> sample, second index ==> char array

// ------- derivative values

	double * OTUs_min;
	double * OTUs_max;
	double * percent_of_samples_OTU_present;


public:

	OTU_Table(char * file_name);
	OTU_Table(OTU_Table * _OTU_t);

	~OTU_Table();

	bool calculate_derivative_values(double OTU_presence_threshold=0.0, ostream & err_msg_out = cout);
	void show_Statistics(ostream & err_msg_out = cout);
	void show_Statistics(ofstream & out_f, ostream & err_msg_out = cout);
	void show_All(ostream & err_msg_out = cout);


//	bool exclude_Infrequent_OTUs(double threshold = 0.05, ostream & err_msg_out = cout);
	
	bool shuffle_inside_OTUs(ostream & err_msg_out = cout);
	bool shuffle_inside_samples(ostream & err_msg_out = cout);

	bool shuffle_All(ostream & err_msg_out = cout);
	bool re_normalize(ostream & err_msg_out = cout);


// NOTE: The OTU_profile array have to be allocated outside 
	bool get_OTU_profile(double * OTU_profile, unsigned int OTU_index, ostream & err_msg_out = cout);
	bool get_OTU_profile(OTU_Profile *& OTU_p, unsigned int OTU_index, ostream & err_msg_out = cout);

// ------ Very special functions....

	bool is_it_make_sense_to_compare_two_OTUs(unsigned int OTU_index_1, unsigned int OTU_index_2, ostream & err_msg_out = cout);

};
//-------------------------------------------------------------------------------------------------------------
//-------------------------------------------------------------------------------------------------------------

bool OTU_Table::is_it_make_sense_to_compare_two_OTUs(unsigned int OTU_index_1, unsigned int OTU_index_2, ostream & err_msg_out)
{
	if (OTU_index_1 == OTU_index_2) return false;
	if (OTU_index_1>number_of_OTUs)return false;
	if (OTU_index_2>number_of_OTUs)return false;

	
	double min_OTU_Value_required = 0.01; // 1%
	double min_percent_of_samples_required = 0.15; // 10%

	if (OTUs_max[OTU_index_1] < min_OTU_Value_required) return false;
	if (OTUs_max[OTU_index_2] < min_OTU_Value_required) return false;

	if (percent_of_samples_OTU_present[OTU_index_1] < min_percent_of_samples_required) return false;
	if (percent_of_samples_OTU_present[OTU_index_2] < min_percent_of_samples_required) return false;

	return true;
}
//-------------------------------------------------------------------------------------------------------------

bool OTU_Table::re_normalize(ostream & err_msg_out)
{
	unsigned int i_samples;
	unsigned int i_OTUs;
	double sum;
	for (i_samples = 0; i_samples < number_of_samples; i_samples++)
	{
		sum = 0.;
		for (i_OTUs = 0; i_OTUs < number_of_OTUs; i_OTUs++)sum += abundance_value[i_samples][i_OTUs];

		if (sum == 0.)
		{
			err_msg_out << "ERROR: OTU_Table::re_normalize()==> sum == 0.\n"; return false;
		}

		for (i_OTUs = 0; i_OTUs < number_of_OTUs; i_OTUs++) abundance_value[i_samples][i_OTUs] /= sum;
	}
	calculate_derivative_values();
	return true;
}

//-------------------------------------------------------------------------------------------------------------

bool OTU_Table::shuffle_All(ostream & err_msg_out)
{
	Shuffle::shuffle_double(abundance_value, number_of_samples, number_of_OTUs, err_msg_out);
//	calculate_derivative_values();
	return true;
}
//-------------------------------------------------------------------------------------------------------------

bool OTU_Table::shuffle_inside_OTUs(ostream & err_msg_out)
{
	double * OTU_profile = new double[number_of_samples];

	unsigned int i_samples;
	unsigned int i_OTUs;

	for (i_OTUs = 0; i_OTUs < number_of_OTUs; i_OTUs++)
	{
		for (i_samples = 0; i_samples < number_of_samples; i_samples++ )
			OTU_profile[i_samples] = abundance_value[i_samples][i_OTUs];
		
		Shuffle::shuffle_double(OTU_profile, number_of_samples, err_msg_out);

		for (i_samples = 0; i_samples < number_of_samples; i_samples++)
			abundance_value[i_samples][i_OTUs]=OTU_profile[i_samples];


	}
	delete[] OTU_profile;
	calculate_derivative_values();
	return true;
}
//-------------------------------------------------------------------------------------------------------------

bool OTU_Table::shuffle_inside_samples(ostream & err_msg_out)
{

	unsigned int i_samples;

	for (i_samples = 0; i_samples < number_of_samples; i_samples++)
		Shuffle::shuffle_double(abundance_value[i_samples], number_of_OTUs, err_msg_out);

	calculate_derivative_values();
	return true;
}
//-------------------------------------------------------------------------------------------------------------

void OTU_Table::show_All(ostream & err_msg_out)
{
	show_Statistics(err_msg_out);

	cout << "============== OTU_Table::show_All ========== (START)\n";


	unsigned int i;	
	for (i = 0; i < number_of_OTUs; i++)
	{
		cout << OTU_short_name[i] << "\t" << taxonomy_text[i] << endl;
		cout << "Present in " << percent_of_samples_OTU_present [i]<< " % of samples "
			<< "min=" << OTUs_min[i]<< "  max=" << OTUs_max[i]<<endl;
	}
	cout << "============== OTU_Table::show_All ========== (END)\n";
}
//-------------------------------------------------------------------------------------------------------------

void OTU_Table::show_Statistics(ostream & err_msg_out)
{
	cout << "============== OTU_Table::show_Statistics ========== (START)\n";
	cout << "\tnumber_of_samples=" << number_of_samples << "\tnumber_of_OTUs=" << number_of_OTUs << endl;

	unsigned int i;
	unsigned int counter = 0;
	for (i = 0; i < number_of_OTUs; i++)
		if (percent_of_samples_OTU_present[i] < 0.001) counter++;

	if (counter > 0)
		cout << "NOTE: " << counter << " OTUs are absent in each sample. " << endl;

	cout << "============== OTU_Table::show_Statistics ========== (END)\n";

}
//-------------------------------------------------------------------------------------------------------------

void OTU_Table::show_Statistics(ofstream & out_f, ostream & err_msg_out)
{
	out_f << "============== OTU_Table::show_Statistics ========== (START)\n";
	out_f << "\tnumber_of_samples=" << number_of_samples << "\nnumber_of_OTUs=" << number_of_OTUs << endl;

	unsigned int i;
	unsigned int counter = 0;
	for (i = 0; i < number_of_OTUs; i++)
		if (percent_of_samples_OTU_present[i] <= 0.) counter++;

	if (counter > 0)
		out_f << "NOTE: " << counter << " OTUs are absent in each sample. " << endl;

	out_f << "============== OTU_Table::show_Statistics ========== (END)\n";

}
//-------------------------------------------------------------------------------------------------------------

bool OTU_Table::calculate_derivative_values(double OTU_presence_threshold, ostream & err_msg_out)
{
	OTUs_min = new double[number_of_OTUs];
	OTUs_max = new double[number_of_OTUs];
	percent_of_samples_OTU_present = new double[number_of_OTUs];
	//cout<<"OTU_Table::calculate_derivative_values Allocation Success"<<endl;

	unsigned int i;
	unsigned int ii;
	for (i = 0; i < number_of_OTUs; i++)
	{
		OTUs_min[i] = 0.;
		OTUs_max[i] = 0.;
		percent_of_samples_OTU_present[i] = 0.;
		

		for (ii = 0; ii < number_of_samples; ii++)
		{

										// first index ==> sample, second index ==> OTU
			if (OTUs_min[i] > abundance_value[ii][i])OTUs_min[i] = abundance_value[ii][i];
			if (OTUs_max[i] < abundance_value[ii][i])OTUs_max[i] = abundance_value[ii][i];
			if (abundance_value[ii][i] > OTU_presence_threshold) percent_of_samples_OTU_present[i]++;
		}

		percent_of_samples_OTU_present[i] = percent_of_samples_OTU_present[i]/(double)number_of_samples;
		
	}
	return true;
}
//-------------------------------------------------------------------------------------------------------------

OTU_Table::OTU_Table(OTU_Table * _OTU_t)
{
//cout << "-------------OTU_Table==> copy constructor--------(START)\n";
	number_of_samples=_OTU_t->number_of_samples;
//cout << "number_of_samples=" << number_of_samples << "\t";

	number_of_OTUs=_OTU_t->number_of_OTUs;
//cout << "number_of_OTUs=" << number_of_OTUs << endl;

	// ------- default parameters
	sample_name_length = 200;
	OTU_short_name_length = 200;
	taxonomy_text_length = 2000;

	// --------- allocations

	unsigned int i;
	sample_name = new char *[number_of_samples];
	for (i = 0; i < number_of_samples; i++) sample_name[i] = new char[sample_name_length];

	OTU_short_name = new char *[number_of_OTUs];
	for (i = 0; i < number_of_OTUs; i++) OTU_short_name[i] = new char[OTU_short_name_length];

	taxonomy_text = new char *[number_of_OTUs];
	for (i = 0; i < number_of_OTUs; i++) taxonomy_text[i] = new char[taxonomy_text_length];

	abundance_value = new double *[number_of_samples];	// first index ==> sample, second index ==> OTU	
	for (i = 0; i < number_of_samples; i++)abundance_value[i] = new double[number_of_OTUs];


	unsigned int i_sample;
	for (i_sample = 0; i_sample < number_of_samples; i_sample++)
	{
		for (i = 0; i < sample_name_length; i++) 
			sample_name[i_sample][i] = _OTU_t->sample_name[i_sample][i];

//cout << "t=" << sample_name[i_sample] << endl;
	}

	unsigned int i_otu;
	for (i_otu = 0; i_otu < number_of_OTUs; i_otu++)
	{
		for (i = 0; i < OTU_short_name_length; i++) 
			OTU_short_name[i_otu][i] = _OTU_t->OTU_short_name[i_otu][i];
//		cout << endl << OTU_short_name[i_otu] << "\t";

		for (i_sample = 0; i_sample < number_of_samples; i_sample++)
		{
			abundance_value[i_sample][i_otu]=_OTU_t->abundance_value[i_sample][i_otu];
//cout << abundance_value[i_sample][i_otu] << "\t";
		}

		for (i = 0; i < taxonomy_text_length; i++)
		taxonomy_text[i_otu][i]= _OTU_t->taxonomy_text[i_otu][i];
//cout << taxonomy_text[i_otu] << endl;
	}
	calculate_derivative_values();

	cout << "-------------OTU_Table==> copy constructor--------(END)\n";
}
//-------------------------------------------------------------------------------------------------------------

bool OTU_Table::get_OTU_profile(double * OTU_profile, unsigned int OTU_index, ostream & err_msg_out)
{
	if (OTU_index >= number_of_OTUs) return false;
	
	unsigned int i;
	for(i=0;i<number_of_samples; i++) OTU_profile[i] = abundance_value[i][OTU_index];
	
	return true;
}
//-------------------------------------------------------------------------------------------------------------

bool OTU_Table::get_OTU_profile(OTU_Profile *& p, unsigned int OTU_index, ostream & err_msg_out)
{
	if (OTU_index >= number_of_OTUs) return false;
	p= new OTU_Profile(number_of_samples);

	unsigned int i;
	for (i = 0; i < OTU_short_name_length; i++) p->OTU_short_name[i] = OTU_short_name[OTU_index][i];
	for (i = 0; i < taxonomy_text_length; i++) p->taxonomy_text[i] = taxonomy_text[OTU_index][i];

	for (i = 0; i < number_of_samples; i++) p->abundance_value[i] = abundance_value[i][OTU_index];

	p->calculate_derivative_values();
	return true;
}
//-------------------------------------------------------------------------------------------------------------

OTU_Table::OTU_Table(char * file_name)
{
    ifstream in_f;
    //Count number of samples and OTU's before processing the file
    in_f.open(file_name);
    number_of_OTUs=0;
    number_of_samples=0;
    bool samples_counted=false;

    if (in_f.is_open())
    {
        string line="";
        while (getline(in_f, line))
        {
            number_of_OTUs++;
            if(!samples_counted)
            for(unsigned int l=0;l<line.length();l++)
            {
            	if(line[l]=='\t') number_of_samples++;
            	samples_counted=true;
            }

        }
        in_f.close();
    }

    
    in_f.clear();                 // clear fail and eof bits
    in_f.seekg(0, std::ios::beg);

    number_of_samples=number_of_samples-1; //discount taxonomy tab to the right
    number_of_OTUs=number_of_OTUs-1;

	//cout << "-------------Reading OTU file --------------\n" << file_name << endl;

	in_f.open(file_name);

    
	//cout << "number_of_samples=" << number_of_samples << "\t";

    
	//cout << "number_of_OTUs=" << number_of_OTUs << endl;

// ------- default parameters
	sample_name_length = 200;
	OTU_short_name_length = 200;
	taxonomy_text_length = 2000;

// --------- allocations

	unsigned int i;
	sample_name = new char *[number_of_samples];
	for (i = 0; i < number_of_samples; i++) sample_name[i] = new char[sample_name_length];

	OTU_short_name = new char *[number_of_OTUs];	
	for (i = 0; i < number_of_OTUs; i++) OTU_short_name[i] = new char[OTU_short_name_length];

	taxonomy_text = new char *[number_of_OTUs];	
	for (i = 0; i < number_of_OTUs; i++) taxonomy_text[i] = new char[taxonomy_text_length];

	abundance_value=new double * [number_of_samples];	// first index ==> sample, second index ==> OTU	
	for (i = 0; i < number_of_samples; i++)abundance_value[i] = new double[number_of_OTUs];
	
	char * temp_string = new char[500];
	
// ------------------------- reading first row

	in_f.getline(temp_string, 500, '\t');	// read word "otu_id" 
	//cout<<"reading first row: "<<temp_string<< endl;

	unsigned int i_sample;
	for (i_sample = 0; i_sample < number_of_samples; i_sample++)
	{
		in_f.getline(sample_name[i_sample], sample_name_length, '\t');	
	}

	in_f.getline(temp_string, 500, '\n');	// read word "taxa" 


// ------------ read OTU row
	unsigned int i_otu;
	for (i_otu = 0; i_otu < number_of_OTUs; i_otu++)
	{
		in_f.getline(OTU_short_name[i_otu], OTU_short_name_length, '\t');


		for (i_sample = 0; i_sample < number_of_samples; i_sample++)
		{
			in_f >> abundance_value[i_sample][i_otu];

		}

		in_f.getline(taxonomy_text[i_otu], taxonomy_text_length, '\n');
		//cout<<"OTU#"<<i_otu<<taxonomy_text[i_otu]<<endl;
	}
	

	calculate_derivative_values();

//cout<<"----Reading complete-------\n"; 
delete[] temp_string;
}
//-------------------------------------------------------------------------------------------------------------

OTU_Table::~OTU_Table()
{
	unsigned int i;

	for (i = 0; i < number_of_samples; i++) delete [] abundance_value[i];
	delete [] abundance_value;

	for (i = 0; i < number_of_samples; i++) delete [] sample_name[i];
	delete [] sample_name;

	for (i = 0; i < number_of_OTUs; i++) delete [] OTU_short_name[i];
	delete [] OTU_short_name;

	for (i = 0; i < number_of_OTUs; i++) delete[] taxonomy_text[i];
	delete [] taxonomy_text;
}


//-------------------------------------------------------------------------------------------------------------
//-------------------------------------------------------------------------------------------------------------
//-------------------------------------------------------------------------------------------------------------
//-------------------------------------------------------------------------------------------------------------
#endif //_OTU_TABLE_H_
```
