## Supplementary material for "Identification of Complex Multidimensional Patterns in Microbial Communities": Shuffle.h

```
/*
THE SOFTWARE IS PROVIDED "AS IS", WITHOUT WARRANTY OF ANY KIND, EXPRESS OR IMPLIED, INCLUDING BUT NOT LIMITED TO THE WARRANTIES OF
MERCHANTABILITY, FITNESS FOR A PARTICULAR PURPOSE AND NONINFRINGEMENT. IN NO EVENT SHALL THE AUTHORS OR COPYRIGHT HOLDERS BE LIABLE FOR
ANY CLAIM, DAMAGES OR OTHER LIABILITY, WHETHER IN AN ACTION OF CONTRACT, TORT OR OTHERWISE, ARISING FROM, OUT OF OR IN CONNECTION WITH
THE SOFTWARE OR THE USE OR OTHER DEALINGS IN THE SOFTWARE.
*/
#ifndef _SHUFFLE_H_
#define _SHUFFLE_H_


#include <iostream>
#include <fstream>
#include <stdlib.h>
#include <random>
#include <time.h>

using namespace std;

class Shuffle
{
	static mt19937 mt_rand;
public:

//Must be called once per program before using the shuffle method.
	static void initialize() { mt_rand.seed(time(0)); };

	// 1D ------------------
	static bool shuffle_bool(bool * value, unsigned int number_of_values, ostream & err_msg_out = cout);
	static bool shuffle_double(double * value, unsigned int number_of_values, ostream & err_msg_out = cout);
	static bool shuffle_unsigned_int(unsigned int * value, unsigned int number_of_values, ostream & err_msg_out = cout);
	static bool shuffle_int(int * value, unsigned int number_of_values, ostream & err_msg_out = cout);

	// 2D ----------------
	//n1 => rows, n2 => columns
	static bool shuffle_bool(bool ** value, unsigned int n1, unsigned int n2, ostream & err_msg_out = cout);	
	static bool shuffle_double(double ** value, unsigned int n1, unsigned int n2, ostream & err_msg_out = cout);
	static bool shuffle_unsigned_int(unsigned int ** value, unsigned int n1, unsigned int n2, ostream & err_msg_out = cout);
	static bool shuffle_int(int ** value, unsigned int n1, unsigned int n2, ostream & err_msg_out = cout);

//			 Service functions -----------------

};
//-------------------------------------------------------------------------------------------------------------
//-------------------------------------------------------------------------------------------------------------
//-------------------------------------------------------------------------------------------------------------
//-------------------------------------------------------------------------------------------------------------
//-------------------------------------------------------------------------------------------------------------

mt19937 Shuffle::mt_rand;

bool Shuffle::shuffle_bool(bool * value, unsigned int number_of_values, ostream & err_msg_out)
{
	//	if(value==NULL)
	if (number_of_values == 0) return false;
	if (number_of_values == 1) return true;

	unsigned int i;
	unsigned int k;
	for (i = 0; i <= number_of_values - 2; i++)
	{
		k = i + mt_rand() % (number_of_values - i);
		swap(value[k], value[i]);
	}
	return true;
}

//-------------------------------------------------------------------------------------------------------------
bool Shuffle::shuffle_double(double * value, unsigned int number_of_values, ostream & err_msg_out)
{
	if (number_of_values == 0) return false;
	if (number_of_values == 1) return true;

	unsigned int i;
	unsigned int k;
	for (i = 0; i <= number_of_values - 2; i++)
	{
		k = i + mt_rand() % (number_of_values - i);
		swap(value[k], value[i]);
	}
	return true;
}

//-------------------------------------------------------------------------------------------------------------
bool Shuffle::shuffle_unsigned_int(unsigned int * value, unsigned int number_of_values, ostream & err_msg_out)
{
	if (number_of_values == 0) return false;
	if (number_of_values == 1) return true;

	unsigned int i;
	unsigned int k;
	for (i = 0; i <= number_of_values - 2; i++)
	{
		k = i + mt_rand() % (number_of_values - i);
		swap(value[k], value[i]);
	}
	return true;
}

//-------------------------------------------------------------------------------------------------------------
bool Shuffle::shuffle_int(int * value, unsigned int number_of_values, ostream & err_msg_out)
{
	if (number_of_values == 0) return false;
	if (number_of_values == 1) return true;

	unsigned int i;
	unsigned int k;
	for (i = 0; i <= number_of_values - 2; i++)
	{
		k = i + mt_rand() % (number_of_values - i);
		swap(value[k], value[i]);
	}
	return true;
}
//-------------------------------------------------------------------------------------------------------------

bool Shuffle::shuffle_bool(bool ** value, unsigned int n1, unsigned int n2, ostream & err_msg_out)
{
	if (n1 == 0) return false;
	if (n2 == 0) return false;
	if (n1 == 1 && n2 == 1) return true;

	unsigned int number_of_values = n1*n2;
	unsigned int i;
	unsigned int k;
	for (i = 0; i <= number_of_values - 2; i++)
	{
		k = i + mt_rand() % (number_of_values - i);
		swap(value[k / n2][k % n2], value[i / n2][i % n2]);
	}
	return true;
}
//-------------------------------------------------------------------------------------------------------------

bool Shuffle::shuffle_double(double ** value, unsigned int n1, unsigned int n2, ostream & err_msg_out)
{
	if (n1 == 0) return false;
	if (n2 == 0) return false;
	if (n1 == 1 && n2 == 1) return true;

	unsigned int number_of_values = n1*n2;
	unsigned int i;
	unsigned int k;
	for (i = 0; i <= number_of_values - 2; i++)
	{
		k = i + mt_rand() % (number_of_values - i);
		swap(value[k / n2][k % n2], value[i / n2][i % n2]);
	}
	return true;
}
//-------------------------------------------------------------------------------------------------------------

bool Shuffle::shuffle_unsigned_int(unsigned int ** value, unsigned int n1, unsigned int n2, ostream & err_msg_out)
{
	if (n1 == 0) return false;
	if (n2 == 0) return false;
	if (n1 == 1 && n2 == 1) return true;

	unsigned int number_of_values = n1*n2;
	unsigned int i;
	unsigned int k;
	for (i = 0; i <= number_of_values - 2; i++)
	{
		k = i + mt_rand() % (number_of_values - i);
		swap(value[k / n2][k % n2], value[i / n2][i % n2]);
	}
	return true;
}
//-------------------------------------------------------------------------------------------------------------

bool Shuffle::shuffle_int(int ** value, unsigned int n1, unsigned int n2, ostream & err_msg_out)
{
	if (n1 == 0) return false;
	if (n2 == 0) return false;
	if (n1 == 1 && n2 == 1) return true;

	unsigned int number_of_values = n1*n2;
	unsigned int i;
	unsigned int k;
	for (i = 0; i <= number_of_values - 2; i++)
	{
		k = i + mt_rand() % (number_of_values - i);
		swap(value[k / n2][k % n2], value[i / n2][i % n2]);
	}
	return true;
}

//-------------------------------------------------------------------------------------------------------------
//-------------------------------------------------------------------------------------------------------------
#endif //_SHUFFLE_H_
```
