## Supplementary material for "Identification of Complex Multidimensional Patterns in Microbial Communities": Stat_and_Sort.h

```
/*
THE SOFTWARE IS PROVIDED "AS IS", WITHOUT WARRANTY OF ANY KIND, EXPRESS OR IMPLIED, INCLUDING BUT NOT LIMITED TO THE WARRANTIES OF
MERCHANTABILITY, FITNESS FOR A PARTICULAR PURPOSE AND NONINFRINGEMENT. IN NO EVENT SHALL THE AUTHORS OR COPYRIGHT HOLDERS BE LIABLE FOR
ANY CLAIM, DAMAGES OR OTHER LIABILITY, WHETHER IN AN ACTION OF CONTRACT, TORT OR OTHERWISE, ARISING FROM, OUT OF OR IN CONNECTION WITH
THE SOFTWARE OR THE USE OR OTHER DEALINGS IN THE SOFTWARE.
*/
#ifndef _STAT_AND_SORT_H_
#define _STAT_AND_SORT_H_


#include <iostream>
#include <fstream>
#include <stdlib.h>
#include <random>
#include <time.h>

using namespace std;

class Stat_and_Sort
{
	
public:	
	static bool sort_Int(int * value, unsigned int number_of_values, ostream & err_msg_out=cout);
	static bool sort_Double(double * value, unsigned int number_of_values, ostream & err_msg_out=cout);

	static bool calculate_Basic_Statistics_Int(int * value, unsigned int number_of_values, 
												double & average, double & stDev,
												double & min_v, double & max_v,
												ostream & err_msg_out=cout);	
	
	static bool calculate_Basic_Statistics_Double(double * value, unsigned int number_of_values, 
												double & average, double & stDev,
												double & min_v, double & max_v,
												ostream & err_msg_out=cout);

// ============== Probabilities ================================

	static bool One_D_Probabilities(bool * x, unsigned int _array_size, double & p_0, double & p_1, ostream & err_msg_out = cout);
	static bool Two_D_Probabilities(bool * x_1, bool * x_2, unsigned int _array_size,
									double & p_00, double & p_01, double & p_10, double & p_11,
									ostream & err_msg_out = cout);

	static bool Three_D_Probabilities(bool * x_1, bool * x_2, bool * x_3, unsigned int _array_size,
										double & p_000, double & p_001, double & p_010, double & p_011,
										double & p_100, double & p_101, double & p_110, double & p_111,
										ostream & err_msg_out = cout);

	// ------- Same but double...

	static bool One_D_Probabilities(double * x, unsigned int _array_size, double threshold, double & p_0, double & p_1, ostream & err_msg_out = cout);

	static bool Two_D_Probabilities(double * x_1, double * x_2, unsigned int _array_size, double threshold_x1, double threshold_x2,
									double & p_00, double & p_01, double & p_10, double & p_11,
									ostream & err_msg_out = cout);

	static bool Three_D_Probabilities(double * x_1, double * x_2, double * x_3, unsigned int _array_size,
										double threshold_x1, double threshold_x2, double threshold_x3,
										double & p_000, double & p_001, double & p_010, double & p_011,
										double & p_100, double & p_101, double & p_110, double & p_111,
										ostream & err_msg_out = cout);

};
//-------------------------------------------------------------------------------------------------------------
//-------------------------------------------------------------------------------------------------------------
//-------------------------------------------------------------------------------------------------------------
//-------------------------------------------------------------------------------------------------------------
//-------------------------------------------------------------------------------------------------------------

bool Stat_and_Sort::One_D_Probabilities(bool * x, unsigned int _array_size, double & p_0, double & p_1, ostream & err_msg_out)
{
	p_0 = 0.;
	p_1 = 0.;
	unsigned int i;
	for (i = 0; i < _array_size; i++)
		if (x[i]) p_1 += 1.;
		else p_0 += 1.;

		p_0 = p_0 / (double)_array_size;
		p_1 = p_1 / (double)_array_size;

		return true;
}
//-------------------------------------------------------------------------------------------------------------

bool Stat_and_Sort::One_D_Probabilities(double * x, unsigned int _array_size, double threshold, double & p_0, double & p_1, ostream & err_msg_out)
{
	p_0 = 0.;
	p_1 = 0.;
	unsigned int i;
	for (i = 0; i < _array_size; i++)
		if (x[i] >= threshold) p_1 += 1.;
		else p_0 += 1.;

		p_0 = p_0 / (double)_array_size;
		p_1 = p_1 / (double)_array_size;

		return true;
}
//-------------------------------------------------------------------------------------------------------------

bool Stat_and_Sort::Two_D_Probabilities(bool * x_1, bool * x_2, unsigned int _array_size,
	double & p_00, double & p_01, double & p_10, double & p_11, ostream & err_msg_out)

{
	p_00 = 0.;
	p_01 = 0.;
	p_10 = 0.;
	p_11 = 0.;

	unsigned int i;

	for (i = 0; i < _array_size; i++)
	{
		if (!x_1[i] && !x_2[i]) { p_00 += 1.; continue; }
		if (x_1[i] && x_2[i]) { p_11 += 1.; continue; }
		if (x_1[i] && !x_2[i]) { p_10 += 1.; continue; }
		if (!x_1[i] && x_2[i]) { p_01 += 1.; continue; }
	}


	p_00 = p_00 / (double)_array_size;
	p_01 = p_01 / (double)_array_size;
	p_10 = p_10 / (double)_array_size;
	p_11 = p_11 / (double)_array_size;

	return true;
}
//-------------------------------------------------------------------------------------------------------------

bool Stat_and_Sort::Two_D_Probabilities(double * x_1, double * x_2, unsigned int _array_size, double threshold_x1, double threshold_x2,
	double & p_00, double & p_01, double & p_10, double & p_11, ostream & err_msg_out)
{
	p_00 = 0.;
	p_01 = 0.;
	p_10 = 0.;
	p_11 = 0.;

	unsigned int i;

	for (i = 0; i < _array_size; i++)
	{
		if (x_1[i] <  threshold_x1  &&  x_2[i] <  threshold_x2) { p_00 += 1.; continue; }
		if (x_1[i] >= threshold_x1 && x_2[i] >= threshold_x2) { p_11 += 1.; continue; }
		if (x_1[i] >= threshold_x1 && x_2[i] <  threshold_x2) { p_10 += 1.; continue; }
		if (x_1[i] <  threshold_x1  &&  x_2[i] >= threshold_x2) { p_01 += 1.; continue; }
	}


	p_00 = p_00 / (double)_array_size;
	p_01 = p_01 / (double)_array_size;
	p_10 = p_10 / (double)_array_size;
	p_11 = p_11 / (double)_array_size;

	return true;
}
//-------------------------------------------------------------------------------------------------------------

bool Stat_and_Sort::Three_D_Probabilities(bool * x_1, bool * x_2, bool * x_3, unsigned int _array_size,
	double & p_000, double & p_001, double & p_010, double & p_011,
	double & p_100, double & p_101, double & p_110, double & p_111,
	ostream & err_msg_out)
{
	p_000 = 0.;
	p_001 = 0.;
	p_010 = 0.;
	p_011 = 0.;
	p_100 = 0.;
	p_101 = 0.;
	p_110 = 0.;
	p_111 = 0.;

	unsigned int i;

	for (i = 0; i < _array_size; i++)
	{
		if (!x_1[i] && !x_2[i] && !x_3[i]) { p_000 += 1.; continue; }
		if (!x_1[i] && !x_2[i] && x_3[i]) { p_001 += 1.; continue; }
		if (!x_1[i] && x_2[i] && !x_3[i]) { p_010 += 1.; continue; }
		if (!x_1[i] && x_2[i] && x_3[i]) { p_011 += 1.; continue; }

		if (x_1[i] && !x_2[i] && !x_3[i]) { p_100 += 1.; continue; }
		if (x_1[i] && !x_2[i] && x_3[i]) { p_101 += 1.; continue; }
		if (x_1[i] && x_2[i] && !x_3[i]) { p_110 += 1.; continue; }
		if (x_1[i] && x_2[i] && x_3[i]) { p_111 += 1.; continue; }
	}


	p_000 = p_000 / (double)_array_size;
	p_001 = p_001 / (double)_array_size;
	p_010 = p_010 / (double)_array_size;
	p_011 = p_011 / (double)_array_size;
	p_100 = p_100 / (double)_array_size;
	p_101 = p_101 / (double)_array_size;
	p_110 = p_110 / (double)_array_size;
	p_111 = p_111 / (double)_array_size;

	return true;
}
//-------------------------------------------------------------------------------------------------------------

bool Stat_and_Sort::Three_D_Probabilities(double * x_1, double * x_2, double * x_3, unsigned int _array_size,
	double threshold_x1, double threshold_x2, double threshold_x3,
	double & p_000, double & p_001, double & p_010, double & p_011,
	double & p_100, double & p_101, double & p_110, double & p_111,
	ostream & err_msg_out)
{
	p_000 = 0.;
	p_001 = 0.;
	p_010 = 0.;
	p_011 = 0.;
	p_100 = 0.;
	p_101 = 0.;
	p_110 = 0.;
	p_111 = 0.;

	unsigned int i;

	for (i = 0; i < _array_size; i++)
	{
		if (x_1[i] < threshold_x1 && x_2[i] < threshold_x2  && x_3[i] < threshold_x3) { p_000 += 1.; continue; }
		if (x_1[i] < threshold_x1 && x_2[i] < threshold_x2  && x_3[i] >= threshold_x3) { p_001 += 1.; continue; }
		if (x_1[i] < threshold_x1 && x_2[i] >= threshold_x2 && x_3[i] < threshold_x3) { p_010 += 1.; continue; }
		if (x_1[i] < threshold_x1 && x_2[i] >= threshold_x2 && x_3[i] >= threshold_x3) { p_011 += 1.; continue; }

		if (x_1[i] >= threshold_x1 && x_2[i] < threshold_x2  && x_3[i] < threshold_x3) { p_100 += 1.; continue; }
		if (x_1[i] >= threshold_x1 && x_2[i] < threshold_x2  && x_3[i] >= threshold_x3) { p_101 += 1.; continue; }
		if (x_1[i] >= threshold_x1 && x_2[i] >= threshold_x2 && x_3[i] < threshold_x3) { p_110 += 1.; continue; }
		if (x_1[i] >= threshold_x1 && x_2[i] >= threshold_x2 && x_3[i] >= threshold_x3) { p_111 += 1.; continue; }
	}


	p_000 = p_000 / (double)_array_size;
	p_001 = p_001 / (double)_array_size;
	p_010 = p_010 / (double)_array_size;
	p_011 = p_011 / (double)_array_size;
	p_100 = p_100 / (double)_array_size;
	p_101 = p_101 / (double)_array_size;
	p_110 = p_110 / (double)_array_size;
	p_111 = p_111 / (double)_array_size;

	return true;
}
//-------------------------------------------------------------------------------------------------------------
//-------------------------------------------------------------------------------------------------------------

bool Stat_and_Sort::sort_Int(int * value, unsigned int number_of_values, ostream & err_msg_out)
{
//	if(value == NULL)
//	if (number_of_values == 0) return false;
//	if (number_of_values == 1) return true;
	
	
	
	bool swapped=true;
	unsigned int i;

	while(swapped)
	{
		swapped = false;
		for (i=1; i<number_of_values; i++)
		{
			if(value[i-1] > value[i])
			{
				swap(value[i], value[i-1]);
				swapped = true;
			}
		}
	}
	return true;
}
//-------------------------------------------------------------------------------------------------------------

bool Stat_and_Sort::sort_Double(double * value, unsigned int number_of_values, ostream & err_msg_out)
{

	//	if(value == NULL)
	//	if (number_of_values == 0) return false;
	//	if (number_of_values == 1) return true;


	bool swapped=true;
	unsigned int i;

	while(swapped)
	{
		swapped = false;
		for (i=1; i<number_of_values; i++)
		{
			if(value[i-1] > value[i])
			{
				swap(value[i], value[i-1]);
				swapped = true;
			}
		}
	}
	return true;
}
//-------------------------------------------------------------------------------------------------------------

bool Stat_and_Sort::calculate_Basic_Statistics_Int(int * value, unsigned int number_of_values, 
												double & average, double & stDev,
												double & min_v, double & max_v,
												ostream & err_msg_out)
{
	//	if(value == NULL)
	//	if (number_of_values == 0) return false;
	//	if (number_of_values == 1) return true;

	average=0.;
	stDev=0.;
	min_v=value[0];
	max_v=value[0];

	unsigned int i;
	for(i=0;i<number_of_values;i++)
	{
		average+=value[i];
		if(value[i]>max_v)max_v=value[i];
		if(value[i]<min_v)min_v=value[i];
	}
	average=average/number_of_values;

	for(i=0;i<number_of_values;i++)stDev+=(value[i]-average)*(value[i]-average);

	stDev=sqrt(stDev/(number_of_values-1));

	return true;
}
//-------------------------------------------------------------------------------------------------------------

bool Stat_and_Sort::calculate_Basic_Statistics_Double(double * value, unsigned int number_of_values, 
													double & average, double & stDev,
													double & min_v, double & max_v,
													ostream & err_msg_out)
{
	//	if(value == NULL)
	//	if (number_of_values == 0) return false;
	//	if (number_of_values == 1) return true;

	average=0.;
	stDev=0.;
	min_v=value[0];
	max_v=value[0];

	unsigned int i;
	for(i=0;i<number_of_values;i++)
	{
		average+=value[i];
		if(value[i]>max_v)max_v=value[i];
		if(value[i]<min_v)min_v=value[i];
	}
	average=average/number_of_values;

	for(i=0;i<number_of_values;i++)stDev+=(value[i]-average)*(value[i]-average);

	stDev=sqrt(stDev/(number_of_values-1));

	return true;
}


//-------------------------------------------------------------------------------------------------------------
#endif //_STAT_AND_SORT_H_
```
