## Supplementary material for "Identification of Complex Multidimensional Patterns in Microbial Communities": Three_D_Patterns_Pipelines.h

```
/*
THE SOFTWARE IS PROVIDED "AS IS", WITHOUT WARRANTY OF ANY KIND, EXPRESS OR IMPLIED, INCLUDING BUT NOT LIMITED TO THE WARRANTIES OF
MERCHANTABILITY, FITNESS FOR A PARTICULAR PURPOSE AND NONINFRINGEMENT. IN NO EVENT SHALL THE AUTHORS OR COPYRIGHT HOLDERS BE LIABLE FOR
ANY CLAIM, DAMAGES OR OTHER LIABILITY, WHETHER IN AN ACTION OF CONTRACT, TORT OR OTHERWISE, ARISING FROM, OUT OF OR IN CONNECTION WITH
THE SOFTWARE OR THE USE OR OTHER DEALINGS IN THE SOFTWARE.
*/
#ifndef _THREE_D_PATTERNS_PIPELINES_H_
#define _THREE_D_PATTERNS_PIPELINES_H_

#include <iostream>
#include <fstream>

#include "Shuffle.h"
#include "Stat_and_Sort.h"

#include "OTU_Table.h"
#include "OTU_Profile.h"

#include "Two_D_Pattern.h"
#include "Three_D_Pattern.h"

#include "Array_of_Two_D_Patterns.h"
#include "Array_of_Three_D_Patterns.h"

#include "Two_D_Patterns_Toolbox.h"
#include "Three_D_Patterns_Toolbox.h"

using namespace std;

class Three_D_Patterns_Pipelines
{
	
public:	

	static void find_3D_Type_2_co_exclusion_Patterns_for_each_Score_and_Presence_Threshold(char * OTU_file_name, char * out_file_name, char * out_file_name_long,
																				double min_threshold = 0., double max_threshold = 0.005, double threshold_step = 0.0005,
																				double min_population_threshold = 0.1, double max_population_threshold = 0.2, double population_threshold_step = 0.01,
																				double min_score = 0.95, double max_score = 1., double score_step = 0.01);


	static void find_3D_Pattern_X23_copresent_if_X1_present_and_co_exclude_if_X1_absent_for_each_Score_and_Presence_Threshold(char * OTU_file_name, char * out_file_name, char * out_file_name_long,
																				double min_threshold = 0., double max_threshold = 0.005, double threshold_step = 0.0005,
																				double min_population_threshold = 0.1, double max_population_threshold = 0.2, double population_threshold_step = 0.01,
																				double min_score = 0.95, double max_score = 1., double score_step = 0.01);


	static void find_3D_Pattern_Present_all_together_or_Alone_for_each_Score_and_Presence_Threshold(char * OTU_file_name, char * out_file_name, char * out_file_name_long,
																				double min_threshold = 0., double max_threshold = 0.005, double threshold_step = 0.0005,
																				double min_population_threshold = 0.1, double max_population_threshold = 0.2, double population_threshold_step = 0.01,
																				double min_score = 0.95, double max_score = 1., double score_step = 0.01);
};

//-------------------------------------------------------------------------------------------------------------
//-------------------------------------------------------------------------------------------------------------
//-------------------------------------------------------------------------------------------------------------
void Three_D_Patterns_Pipelines::find_3D_Pattern_Present_all_together_or_Alone_for_each_Score_and_Presence_Threshold(char * OTU_file_name, char * out_file_name, char * out_file_name_long,
																	double min_threshold, double max_threshold, double threshold_step,
																	double min_population_threshold, double max_population_threshold, double population_threshold_step,
																	double min_score, double max_score, double score_step)
{
	Shuffle::initialize();

	//cout << "\n=============== Real ========================\n";
	OTU_Table * t = new OTU_Table(OTU_file_name);
	t->re_normalize();
	//t->show_Statistics();
	//t->show_All();


	//cout << "\n=============== Random ========================\n";
	OTU_Table * t_random = new OTU_Table(OTU_file_name);

//	t_random->shuffle_All();/////////////////////////////////////////////////////////////////////////////
	t_random->shuffle_inside_OTUs();

	t_random->re_normalize();
	//t_random->show_Statistics();

	//cout << "++++++++++++++++++++++++++++++++++++++++++++++++++++++++\n";

	ofstream out_f;
	out_f.open(out_file_name);
	ofstream out_f_long;
	out_f_long.open(out_file_name_long);

	double * OTU_1 = new double[t->number_of_samples];
	double * OTU_2 = new double[t->number_of_samples];
	double * OTU_3 = new double[t->number_of_samples];

	double population_threshold;
	double real_data_score_threshold;

// -------- variables


	unsigned int number_of_scores = (unsigned int)((max_score - min_score) / score_step);

	unsigned int * number_of_patterns_for_given_score = new unsigned int[number_of_scores];

	Array_of_Three_D_Patterns * patterns_array = new Array_of_Three_D_Patterns();

	bool flag;
	unsigned int patterns_counter;
	unsigned int smallest_score_threshold_index;
	Three_D_Pattern * pattern = NULL;
	unsigned int i;;
	unsigned int ii;
	unsigned int iii;
	unsigned int j;
	double score;

	for (population_threshold = min_population_threshold; population_threshold < max_population_threshold; population_threshold += population_threshold_step)
	{
		for (j = 0; j < number_of_scores; j++) number_of_patterns_for_given_score[j] = 0;
		flag = false;	

		//cout << "population_threshold=" << population_threshold << endl;

		score = 0.;

		for (i = 0; i < t_random->number_of_OTUs - 2; i++)
		{
			//cout << "\nFirst OTU index = " << i << "\n Second index \t";

			for (ii = i + 1; ii < t_random->number_of_OTUs - 1; ii++)
			{
				//cout << ii << "\t";

				for (iii = ii + 1; iii < t_random->number_of_OTUs; iii++)
				{

					t_random->get_OTU_profile(OTU_1, i);
					t_random->get_OTU_profile(OTU_2, ii);
					t_random->get_OTU_profile(OTU_3, iii);


					Three_D_Patterns_Toolbox::find_3D_Pattern_All_Together_or_Alone_ONLY_using_GRID(OTU_1, OTU_2, OTU_3, t_random->number_of_samples,
						score, min_population_threshold, min_threshold, max_threshold, threshold_step);


					for (j = 0; j < number_of_scores; j++)
						if (score >= min_score + score_step * j) number_of_patterns_for_given_score[j]++;


					if (number_of_patterns_for_given_score[number_of_scores - 1] > 0) { flag = true; break; }
				}
				if (flag) break;
			}
			if (flag) break;
		}

		if (flag) continue;


// are there any zeros on the number_of_patterns_for_given_score?

		smallest_score_threshold_index = 0;
		for (j = 0; j < number_of_scores; j++)
		{
			if (number_of_patterns_for_given_score[j] == 0)
			{
				smallest_score_threshold_index = j;
				break;
			}
		}

		if (smallest_score_threshold_index == number_of_scores) continue; // There are no zeros. No reason to try to run real data


 // run real data...

		real_data_score_threshold = min_score + score_step * smallest_score_threshold_index;

		//cout << " real_data_score_threshold=" << real_data_score_threshold << endl;

		patterns_counter = 0;

		for (i = 0; i < t->number_of_OTUs - 2; i++)
		{
			//cout << "\nREAL: First OTU Index = " << i << "\nSecond:\t";
			for (ii = i + 1; ii < t->number_of_OTUs - 1; ii++)
			{
				//cout << ii << "\t";
				for (iii = ii + 1; iii < t->number_of_OTUs; iii++)
				{
					t->get_OTU_profile(OTU_1, i);
					t->get_OTU_profile(OTU_2, ii);
					t->get_OTU_profile(OTU_3, iii);

					Three_D_Patterns_Toolbox::find_3D_Pattern_All_Together_or_Alone_using_GRID(OTU_1, OTU_2, OTU_3, t->number_of_samples, pattern, population_threshold,
																								min_threshold, max_threshold, threshold_step);


					if (pattern->score >= real_data_score_threshold)
					{
						pattern->OTU_Index_1 = i;
						pattern->OTU_Index_2 = ii;
						pattern->OTU_Index_3 = iii;

						if (patterns_array->is_Pattern_Present(pattern))
						{
							//cout << "duplicated pattern!\n";
							delete pattern;
							pattern = NULL;
						}
						else
						{
							patterns_array->add_Pattern(pattern);
							patterns_counter++;
							//cout << "patterns_counter=" << patterns_counter++ << endl;
						}
					}
					else
					{
						delete pattern;
						pattern = NULL;
					}
				}
			}
		}

		if (smallest_score_threshold_index == 0) break;	// no patterns in random data;

	}

// final output....

	OTU_Profile * p;
	for (i = 0; i < patterns_array->number_of_patterns; i++)
	{
		t->get_OTU_profile(p, patterns_array->pattern[i]->OTU_Index_1);
		patterns_array->pattern[i]->OTU_p_1 = p;

		t->get_OTU_profile(p, patterns_array->pattern[i]->OTU_Index_2);
		patterns_array->pattern[i]->OTU_p_2 = p;

		t->get_OTU_profile(p, patterns_array->pattern[i]->OTU_Index_3);
		patterns_array->pattern[i]->OTU_p_3 = p;
	}


	//patterns_array->show_Statistics();
	patterns_array->show_Statistics(out_f);


	patterns_array->show_All(out_f_long);

	out_f.close();
	out_f_long.close();

	delete t;
	delete t_random;

	delete[] number_of_patterns_for_given_score;
	delete[] OTU_1;
	delete[] OTU_2;
	delete[] OTU_3;

	delete patterns_array;

}
//-------------------------------------------------------------------------------------------------------------

void Three_D_Patterns_Pipelines::find_3D_Pattern_X23_copresent_if_X1_present_and_co_exclude_if_X1_absent_for_each_Score_and_Presence_Threshold(char * OTU_file_name, char * out_file_name, char * out_file_name_long,
																				double min_threshold, double max_threshold, double threshold_step,
																				double min_population_threshold, double max_population_threshold, double population_threshold_step,
																				double min_score, double max_score, double score_step)
{
	Shuffle::initialize();

	//cout << "\n=============== Real ========================\n";
	OTU_Table * t = new OTU_Table(OTU_file_name);
	t->re_normalize();
	//t->show_Statistics();
	//t->show_All();


	//cout << "\n=============== Random ========================\n";
	OTU_Table * t_random = new OTU_Table(OTU_file_name);

//	t_random->shuffle_All();/////////////////////////////////////////////////////////////////////////////
	t_random->shuffle_inside_OTUs();

	t_random->re_normalize();
	//t_random->show_Statistics();

	//cout << "++++++++++++++++++++++++++++++++++++++++++++++++++++++++\n";

	ofstream out_f;
	out_f.open(out_file_name);
	ofstream out_f_long;
	out_f_long.open(out_file_name_long);

	double * OTU_1 = new double[t->number_of_samples];
	double * OTU_2 = new double[t->number_of_samples];
	double * OTU_3 = new double[t->number_of_samples];

	double population_threshold;
	double real_data_score_threshold;

	unsigned int number_of_scores = (unsigned int)((max_score - min_score) / score_step);

	unsigned int * number_of_patterns_for_given_score = new unsigned int[number_of_scores];

	Array_of_Three_D_Patterns * patterns_array = new Array_of_Three_D_Patterns();

	bool flag;
	unsigned int patterns_counter;
	unsigned int smallest_score_threshold_index;
	Three_D_Pattern * pattern = NULL;
	unsigned int i;;
	unsigned int ii;
	unsigned int iii;
	unsigned int j;
	double score;

	for (population_threshold = min_population_threshold; population_threshold < max_population_threshold; population_threshold += population_threshold_step)
	{
		for (j = 0; j < number_of_scores; j++) number_of_patterns_for_given_score[j] = 0;
		flag = false;

		//cout << "population_threshold=" << population_threshold << endl;

		score = 0.;

		for (i = 0; i < t_random->number_of_OTUs; i++)
		{
			//cout << "\nFirst OTU index = " << i << "\n Second index \t";

			for (ii = 0; ii < t_random->number_of_OTUs-1; ii++)
			{
				if (ii == i) continue;
				//cout << ii << "\t";

				for (iii = ii+1; iii < t_random->number_of_OTUs; iii++)
				{
					if (iii == i) continue;

					t_random->get_OTU_profile(OTU_1, i);
					t_random->get_OTU_profile(OTU_2, ii);
					t_random->get_OTU_profile(OTU_3, iii);


					Three_D_Patterns_Toolbox::find_3D_Pattern_X23_co_present_if_X1_present_othervise_co_excluded_ONLY_using_GRID(OTU_1, OTU_2, OTU_3, t_random->number_of_samples,
																				score, min_population_threshold, min_threshold, max_threshold, threshold_step);

					for (j = 0; j < number_of_scores; j++)
						if (score >= min_score + score_step * j) number_of_patterns_for_given_score[j]++;

					if (number_of_patterns_for_given_score[number_of_scores - 1] > 0) { flag = true;  break; }
				}
				if (flag) break;
			}
			if (flag) break;
		}

		if (flag) continue;


		smallest_score_threshold_index = 0;
		for (j = 0; j < number_of_scores; j++)
		{
			if (number_of_patterns_for_given_score[j] == 0)
			{
				smallest_score_threshold_index = j;
				break;
			}
		}

		if (smallest_score_threshold_index == number_of_scores) continue; // There are no zeros (places without random patterns). No reason to try to run real data

// run real data...

		real_data_score_threshold = min_score + score_step * smallest_score_threshold_index;

		//cout << " real_data_score_threshold=" << real_data_score_threshold << endl;

		patterns_counter = 0;

		for (i = 0; i < t_random->number_of_OTUs; i++)
		{
			//cout << "\nREAL: First OTU index = " << i << "\n Second index \t";

			for (ii = 0; ii < t_random->number_of_OTUs - 1; ii++)
			{
				if (ii == i) continue;
				//cout << ii << "\t";

				for (iii = ii + 1; iii < t_random->number_of_OTUs; iii++)
				{
					if (iii == i) continue;
					t->get_OTU_profile(OTU_1, i);
					t->get_OTU_profile(OTU_2, ii);
					t->get_OTU_profile(OTU_3, iii);

					Three_D_Patterns_Toolbox::find_3D_Pattern_X23_co_present_if_X1_present_othervise_co_excluded_using_GRID(OTU_1, OTU_2, OTU_3, t->number_of_samples, pattern, population_threshold,
						min_threshold, max_threshold, threshold_step);


					if (pattern->score >= real_data_score_threshold)
					{
						pattern->OTU_Index_1 = i;
						pattern->OTU_Index_2 = ii;
						pattern->OTU_Index_3 = iii;

						if (patterns_array->is_Pattern_Present(pattern))
						{
							//cout << "duplicated pattern!\n";
							delete pattern;
							pattern = NULL;
						}
						else
						{
							patterns_array->add_Pattern(pattern);
							patterns_counter++;
							//cout << "patterns_counter=" << patterns_counter++ << endl;
						}
					}
					else
					{
						delete pattern;
						pattern = NULL;
					}
				}
			}
		}

		if (smallest_score_threshold_index == 0) break;	// no patterns in random data;

	}

	// final output....

	OTU_Profile * p;
	for (i = 0; i < patterns_array->number_of_patterns; i++)
	{
		t->get_OTU_profile(p, patterns_array->pattern[i]->OTU_Index_1);
		patterns_array->pattern[i]->OTU_p_1 = p;

		t->get_OTU_profile(p, patterns_array->pattern[i]->OTU_Index_2);
		patterns_array->pattern[i]->OTU_p_2 = p;

		t->get_OTU_profile(p, patterns_array->pattern[i]->OTU_Index_3);
		patterns_array->pattern[i]->OTU_p_3 = p;
	}

	//patterns_array->show_Statistics();
	patterns_array->show_Statistics(out_f);


	patterns_array->show_All(out_f_long);

	out_f.close();
	out_f_long.close();

	delete t;
	delete t_random;

	delete[] number_of_patterns_for_given_score;
	delete[] OTU_1;
	delete[] OTU_2;
	delete[] OTU_3;

	delete patterns_array;
}
//-------------------------------------------------------------------------------------------------------------
//-------------------------------------------------------------------------------------------------------------


void Three_D_Patterns_Pipelines::find_3D_Type_2_co_exclusion_Patterns_for_each_Score_and_Presence_Threshold(char * OTU_file_name, char * out_file_name, char * out_file_name_long,
																			double min_threshold, double max_threshold, double threshold_step,
																			double min_population_threshold, double max_population_threshold, double population_threshold_step,
																			double min_score, double max_score, double score_step)
{
	Shuffle::initialize();

	//cout << "\n=============== Real ========================\n";
	OTU_Table * t = new OTU_Table(OTU_file_name);
	t->re_normalize();
	//t->show_Statistics();
	//t->show_All();


	//cout << "\n=============== Random ========================\n";
	OTU_Table * t_random = new OTU_Table(OTU_file_name);

//	t_random->shuffle_All();/////////////////////////////////////////////////////////////////////////////
	t_random->shuffle_inside_OTUs();

	t_random->re_normalize();
	//t_random->show_Statistics();

	//cout << "++++++++++++++++++++++++++++++++++++++++++++++++++++++++\n";

	ofstream out_f;
	out_f.open(out_file_name);
	ofstream out_f_long;
	out_f_long.open(out_file_name_long);


	double * OTU_1 = new double[t->number_of_samples];
	double * OTU_2 = new double[t->number_of_samples];
	double * OTU_3 = new double[t->number_of_samples];

	double population_threshold;
	double real_data_score_threshold;


	unsigned int number_of_scores = (unsigned int)((max_score - min_score) / score_step);

	unsigned int * number_of_patterns_for_given_score = new unsigned int[number_of_scores];

	Array_of_Three_D_Patterns * patterns_array = new Array_of_Three_D_Patterns();

	bool flag;
	unsigned int patterns_counter;
	unsigned int smallest_score_threshold_index;
	Three_D_Pattern * pattern = NULL;
	unsigned int i;;
	unsigned int ii;
	unsigned int iii;
	unsigned int j;
	double score;

	for (population_threshold = min_population_threshold; population_threshold < max_population_threshold; population_threshold += population_threshold_step)
	{
		for (j = 0; j < number_of_scores; j++) number_of_patterns_for_given_score[j] = 0;
		flag = false;

		//cout << "population_threshold=" << population_threshold << endl;

		score = 0.;

		for (i = 0; i < t_random->number_of_OTUs - 2; i++)
		{
			//cout << "\nFirst OTU index = " << i << "\n Second index \t";

			for (ii = i + 1; ii < t_random->number_of_OTUs - 1; ii++)
			{
				//cout << ii << "\t";

				for (iii = ii + 1; iii < t_random->number_of_OTUs; iii++)
				{

					t_random->get_OTU_profile(OTU_1, i);
					t_random->get_OTU_profile(OTU_2, ii);
					t_random->get_OTU_profile(OTU_3, iii);


					Three_D_Patterns_Toolbox::find_3D_Type_2_Co_Exclusion_Score_ONLY_using_GRID(OTU_1, OTU_2, OTU_3, t_random->number_of_samples,
						score, min_population_threshold, min_threshold, max_threshold, threshold_step);

					for (j = 0; j < number_of_scores; j++)
						if (score >= min_score + score_step * j) number_of_patterns_for_given_score[j]++;

					if (number_of_patterns_for_given_score[number_of_scores - 1] > 0) { flag = true; break; }
				}
				if (flag) break;
			}
			if (flag) break;
		}

		if (flag) continue;


// are there any zeros on the number_of_patterns_for_given_score?

		smallest_score_threshold_index = 0;
		for (j = 0; j < number_of_scores; j++)
		{
			if (number_of_patterns_for_given_score[j] == 0)
			{
				smallest_score_threshold_index = j;
				break;
			}
		}


		if (smallest_score_threshold_index == number_of_scores) continue; // There are no zeros (places without random patterns). No reason to try to run real data

// run real data...

		real_data_score_threshold = min_score + score_step * smallest_score_threshold_index;

		//cout << " real_data_score_threshold=" << real_data_score_threshold << endl;

		patterns_counter = 0;

		for (i = 0; i < t->number_of_OTUs - 2; i++)
		{
			//cout << "\nREAL: First OTU Index = " << i << "\nSecond:\t";
			for (ii = i + 1; ii < t->number_of_OTUs - 1; ii++)
			{
				//cout << ii << "\t";
				for (iii = ii + 1; iii < t->number_of_OTUs; iii++)
				{
					t->get_OTU_profile(OTU_1, i);
					t->get_OTU_profile(OTU_2, ii);
					t->get_OTU_profile(OTU_3, iii);

					Three_D_Patterns_Toolbox::find_3D_Type_2_Co_Exclusion_Score_using_GRID(OTU_1, OTU_2, OTU_3, t->number_of_samples, pattern, population_threshold,
						min_threshold, max_threshold, threshold_step);


					if (pattern->score >= real_data_score_threshold)
					{
						pattern->OTU_Index_1 = i;
						pattern->OTU_Index_2 = ii;
						pattern->OTU_Index_3 = iii;

						if (patterns_array->is_Pattern_Present(pattern))
						{
							//cout << "duplicated pattern!\n";
							delete pattern;
							pattern = NULL;
						}
						else
						{
							patterns_array->add_Pattern(pattern);
							patterns_counter++;
							//cout << "patterns_counter=" << patterns_counter++ << endl;
						}
					}
					else
					{
						delete pattern;
						pattern = NULL;
					}
				}
			}
		}

		if (smallest_score_threshold_index == 0) break;	// no patterns in random data;

	}

// final output....

	OTU_Profile * p;
	for (i = 0; i < patterns_array->number_of_patterns; i++)
	{
		t->get_OTU_profile(p, patterns_array->pattern[i]->OTU_Index_1);
		patterns_array->pattern[i]->OTU_p_1 = p;

		t->get_OTU_profile(p, patterns_array->pattern[i]->OTU_Index_2);
		patterns_array->pattern[i]->OTU_p_2 = p;

		t->get_OTU_profile(p, patterns_array->pattern[i]->OTU_Index_3);
		patterns_array->pattern[i]->OTU_p_3 = p;
	}


	patterns_array->show_Statistics();
	patterns_array->show_Statistics(out_f);


	patterns_array->show_All(out_f_long);

	out_f.close();
	out_f_long.close();

	delete t;
	delete t_random;

	delete[] number_of_patterns_for_given_score;
	delete[] OTU_1;
	delete[] OTU_2;
	delete[] OTU_3;

	delete patterns_array;
}
//-------------------------------------------------------------------------------------------------------------


#endif //_THREE_D_PATTERNS_PIPELINES_H_
```
