## Supplementary material for "Identification of Complex Multidimensional Patterns in Microbial Communities": Three_D_Patterns_Toolbox.h

```
/*
THE SOFTWARE IS PROVIDED "AS IS", WITHOUT WARRANTY OF ANY KIND, EXPRESS OR IMPLIED, INCLUDING BUT NOT LIMITED TO THE WARRANTIES OF
MERCHANTABILITY, FITNESS FOR A PARTICULAR PURPOSE AND NONINFRINGEMENT. IN NO EVENT SHALL THE AUTHORS OR COPYRIGHT HOLDERS BE LIABLE FOR
ANY CLAIM, DAMAGES OR OTHER LIABILITY, WHETHER IN AN ACTION OF CONTRACT, TORT OR OTHERWISE, ARISING FROM, OUT OF OR IN CONNECTION WITH
THE SOFTWARE OR THE USE OR OTHER DEALINGS IN THE SOFTWARE.
*/
#ifndef _THREE_D_PATTERNS_TOOLBOX_H_
#define _THREE_D_PATTERNS_TOOLBOX_H_


#include <iostream>
#include "Shuffle.h"
#include "Stat_and_Sort.h"
#include "Two_D_Pattern.h"
#include "Three_D_Pattern.h"

using namespace std;

//-------------------------------------------------------------------------------------------------------------
//This class is designed to calculate various entropy and mutual information scores.
// NOTE: All calculations are performed using base 2 log: ===> log2(2.0)==1.0;
// NOTE: All calculations performed for binay (true(1)/false(0)) values;
//-------------------------------------------------------------------------------------------------------------


class Three_D_Patterns_Toolbox
{
public:

// ============================ Patterns Specific... 3D only
//-------------------------------------------------------------------------------------------------------------
//--------------------------    PATTERN SPECIFIC SCORES    ----------------------------------------------------
//-------------------------------------------------------------------------------------------------------------
// --- Search for patterns  --------------------------------------------------------------------
// ------------------------------------------------------------------------------------------------------------
// For each pattern the "score" is calculated by maximizing the total percent of points associated (belonging) 
// to the pattern under condition that persentage of points in each sector (000,001,...,111) is above the "minimum_population_threshold"
// 
// -------------------------------------------------------------------------------------------------------
//-------------------------------------------------------------------------------------------------------------
//--------------------------    PATTERN SPECIFIC SCORES    ----------------------------------------------------
//-------------------------------------------------------------------------------------------------------------

	static bool find_3D_Type_2_Co_Exclusion_Score_using_GRID(double * x_1, double * x_2, double * x_3, unsigned int _array_size,
																	Three_D_Pattern *& TD_Pattern,
																	double minimum_population_threshold,
																	double min_threshold_x = 0.,
																	double max_threshold_x = 0.3,
																	double threshold_step = 0.0001,
																	ostream & err_msg_out = cout);

	static bool find_3D_Type_2_Co_Exclusion_Score_ONLY_using_GRID(double * x_1, double * x_2, double * x_3, unsigned int _array_size,
																	double & score,
																	double minimum_population_threshold,
																	double min_threshold_x = 0.,
																	double max_threshold_x = 0.3,
																	double threshold_step = 0.0001,
																	ostream & err_msg_out = cout);

//----------------------  another pattern: X2 and X3 co-present in X1 present and co-excluded if X1 absent 
	static bool find_3D_Pattern_X23_co_present_if_X1_present_othervise_co_excluded_using_GRID(double * x_1, double * x_2, double * x_3, unsigned int _array_size,// NEW
																								Three_D_Pattern *& TD_Pattern,
																								double minimum_population_threshold,
																								double min_threshold_x = 0.,
																								double max_threshold_x = 0.3,
																								double threshold_step = 0.0001,
																								ostream & err_msg_out = cout);

	static bool find_3D_Pattern_X23_co_present_if_X1_present_othervise_co_excluded_ONLY_using_GRID(double * x_1, double * x_2, double * x_3, unsigned int _array_size,// NEW
																								double & score,
																								double minimum_population_threshold,
																								double min_threshold_x = 0.,
																								double max_threshold_x = 0.3,
																								double threshold_step = 0.0001,
																								ostream & err_msg_out = cout);

//----------------------  another pattern: All Together or Alone 
	static bool find_3D_Pattern_All_Together_or_Alone_using_GRID(double * x_1, double * x_2, double * x_3, unsigned int _array_size,// NEW
																								Three_D_Pattern *& TD_Pattern,
																								double minimum_population_threshold,
																								double min_threshold_x = 0.,
																								double max_threshold_x = 0.3,
																								double threshold_step = 0.0001,
																								ostream & err_msg_out = cout);

	static bool find_3D_Pattern_All_Together_or_Alone_ONLY_using_GRID(double * x_1, double * x_2, double * x_3, unsigned int _array_size,// NEW
																								double & score,
																								double minimum_population_threshold,
																								double min_threshold_x = 0.,
																								double max_threshold_x = 0.3,
																								double threshold_step = 0.0001,
																								ostream & err_msg_out = cout);


};


//-------------------------------------------------------------------------------------------------------------
//-------------------------------------------------------------------------------------------------------------
//-------------------------------------------------------------------------------------------------------------

bool Three_D_Patterns_Toolbox::find_3D_Pattern_All_Together_or_Alone_ONLY_using_GRID(double * x_1, double * x_2, double * x_3, unsigned int _array_size,
																							double & score,
																							double minimum_population_threshold,
																							double min_threshold_x,
																							double max_threshold_x,
																							double threshold_step,
																							ostream & err_msg_out)
{
	// ---- Initial values----
	score = 0.;

	double p_000 = 0.;
	double p_001 = 0.;
	double p_010 = 0.;
	double p_011 = 0.;
	double p_100 = 0.;
	double p_101 = 0.;
	double p_110 = 0.;
	double p_111 = 0.;

	//----

	double th_1 = 0.;	// threasholds
	double th_2 = 0.;
	double th_3 = 0.;

	// ---- calculating max so no calculations can be performed for threasholds above max values

	double max_x_1 = x_1[0];
	double max_x_2 = x_2[0];
	double max_x_3 = x_3[0];

	unsigned int i;
	for (i = 0; i<_array_size; i++)
	{
		if (max_x_1 < x_1[i]) max_x_1 = x_1[i];
		if (max_x_2 < x_2[i]) max_x_2 = x_2[i];
		if (max_x_3 < x_3[i]) max_x_3 = x_3[i];
	}


	//--------------------- Main loop

	for (th_1 = min_threshold_x; th_1 <= max_threshold_x; th_1 += threshold_step)
	{
		if (th_1 > max_x_1) break;

		for (th_2 = min_threshold_x; th_2 <= max_threshold_x; th_2 += threshold_step)
		{
			if (th_2 > max_x_2) break;

			for (th_3 = min_threshold_x; th_3 <= max_threshold_x; th_3 += threshold_step)
			{
				if (th_3 > max_x_3) break;

				Stat_and_Sort::Three_D_Probabilities(x_1, x_2, x_3, _array_size, th_1, th_2, th_3, p_000, p_001, p_010, p_011, p_100, p_101, p_110, p_111, err_msg_out);

				if (p_111 < minimum_population_threshold || p_001 < minimum_population_threshold || p_010 < minimum_population_threshold || p_001 < minimum_population_threshold) continue;

				if (score < p_111 + p_100 + p_010 + p_001 + p_000)  
					score = p_111 + p_100 + p_010 + p_001 + p_000;
			}
		}
	}
	return true;
}


//-------------------------------------------------------------------------------------------------------------

bool Three_D_Patterns_Toolbox::find_3D_Pattern_All_Together_or_Alone_using_GRID(double * x_1, double * x_2, double * x_3, unsigned int _array_size,
																							Three_D_Pattern *& TD_Pattern,
																							double minimum_population_threshold,
																							double min_threshold_x,
																							double max_threshold_x,
																							double threshold_step,
																							ostream & err_msg_out)
{
	// ---- Initial values----
	double score = 0.;
	double observed_population_threshold = 0.;

	double threshold_x1 = 0.;
	double threshold_x2 = 0.;
	double threshold_x3 = 0.;

	double p_000 = 0.;
	double p_001 = 0.;
	double p_010 = 0.;
	double p_011 = 0.;
	double p_100 = 0.;
	double p_101 = 0.;
	double p_110 = 0.;
	double p_111 = 0.;

	//----

	double th_1 = 0.;	// threshold
	double th_2 = 0.;
	double th_3 = 0.;

	// ---- calculating max so no calculations can be performed for thresholds above max values

	double max_x_1 = x_1[0];
	double max_x_2 = x_2[0];
	double max_x_3 = x_3[0];

	unsigned int i;
	for (i = 0; i<_array_size; i++)
	{
		if (max_x_1 < x_1[i]) max_x_1 = x_1[i];
		if (max_x_2 < x_2[i]) max_x_2 = x_2[i];
		if (max_x_3 < x_3[i]) max_x_3 = x_3[i];
	}


	//--------------------- Main loop

	for (th_1 = min_threshold_x; th_1 <= max_threshold_x; th_1 += threshold_step)
	{
		if (th_1 > max_x_1) break;

		for (th_2 = min_threshold_x; th_2 <= max_threshold_x; th_2 += threshold_step)
		{
			if (th_2 > max_x_2) break;

			for (th_3 = min_threshold_x; th_3 <= max_threshold_x; th_3 += threshold_step)
			{
				if (th_3 > max_x_3) break;

				Stat_and_Sort::Three_D_Probabilities(x_1, x_2, x_3, _array_size, th_1, th_2, th_3, p_000, p_001, p_010, p_011, p_100, p_101, p_110, p_111, err_msg_out);

				if (p_111 < minimum_population_threshold || p_100 < minimum_population_threshold || p_010 < minimum_population_threshold || p_001 < minimum_population_threshold) continue;

				if (score < p_111 + p_100 + p_010 + p_001 + p_000)  
				{
					score = p_111 + p_100 + p_010 + p_001 + p_000;
					threshold_x1 = th_1;
					threshold_x2 = th_2;
					threshold_x3 = th_3;
					observed_population_threshold = min(min(p_111,p_100), min(p_010, p_001));
				}
			}
		}
	}

	//  ---------------- final output -------------

	Stat_and_Sort::Three_D_Probabilities(x_1, x_2, x_3, _array_size, threshold_x1, threshold_x2, threshold_x3, p_000, p_001, p_010, p_011, p_100, p_101, p_110, p_111, err_msg_out);

	TD_Pattern = new Three_D_Pattern();	// Pattern is created, but not filed with all the data.

	TD_Pattern->score = score;
	TD_Pattern->population_threshold = observed_population_threshold;
	TD_Pattern->threshold_x1 = threshold_x1;
	TD_Pattern->threshold_x2 = threshold_x2;
	TD_Pattern->threshold_x3 = threshold_x3;

	TD_Pattern->p_000 = p_000;
	TD_Pattern->p_001 = p_001;
	TD_Pattern->p_010 = p_010;
	TD_Pattern->p_011 = p_011;
	TD_Pattern->p_100 = p_100;
	TD_Pattern->p_101 = p_101;
	TD_Pattern->p_110 = p_110;
	TD_Pattern->p_111 = p_111;

	return true;
}

//-------------------------------------------------------------------------------------------------------------
//-------------------------------------------------------------------------------------------------------------

bool Three_D_Patterns_Toolbox::find_3D_Pattern_X23_co_present_if_X1_present_othervise_co_excluded_ONLY_using_GRID(double * x_1, double * x_2, double * x_3, unsigned int _array_size,
																																		double & score,
																																		double minimum_population_threshold,
																																		double min_threshold_x,
																																		double max_threshold_x,
																																		double threshold_step,
																																		ostream & err_msg_out)
{
	// ---- Initial values----
	score = 0.;
	double observed_population_threshold = 0.;

	double threshold_x1 = 0.;
	double threshold_x2 = 0.;
	double threshold_x3 = 0.;

	double p_000 = 0.;
	double p_001 = 0.;
	double p_010 = 0.;
	double p_011 = 0.;
	double p_100 = 0.;
	double p_101 = 0.;
	double p_110 = 0.;
	double p_111 = 0.;

	//----

	double th_1 = 0.;	// thresholds
	double th_2 = 0.;
	double th_3 = 0.;

	// ---- calculating max so no calculations can be performed for thresholds above max values

	double max_x_1 = x_1[0];
	double max_x_2 = x_2[0];
	double max_x_3 = x_3[0];

	unsigned int i;
	for (i = 0; i<_array_size; i++)
	{
		if (max_x_1 < x_1[i]) max_x_1 = x_1[i];
		if (max_x_2 < x_2[i]) max_x_2 = x_2[i];
		if (max_x_3 < x_3[i]) max_x_3 = x_3[i];
	}


	//--------------------- Main loop

	for (th_1 = min_threshold_x; th_1 <= max_threshold_x; th_1 += threshold_step)
	{
		if (th_1 > max_x_1) break;

		for (th_2 = min_threshold_x; th_2 <= max_threshold_x; th_2 += threshold_step)
		{
			if (th_2 > max_x_2) break;

			for (th_3 = min_threshold_x; th_3 <= max_threshold_x; th_3 += threshold_step)
			{
				if (th_3 > max_x_3) break;

				Stat_and_Sort::Three_D_Probabilities(x_1, x_2, x_3, _array_size, th_1, th_2, th_3, p_000, p_001, p_010, p_011, p_100, p_101, p_110, p_111, err_msg_out);

				if (p_111 < minimum_population_threshold || p_001 < minimum_population_threshold || p_010 < minimum_population_threshold) continue;

				if (score < p_111 + p_001 + p_010 + p_000) 
					score = p_111 + p_001 + p_010 + p_000;
			}
		}
	}
	return true;
}
//-------------------------------------------------------------------------------------------------------------

bool Three_D_Patterns_Toolbox::find_3D_Pattern_X23_co_present_if_X1_present_othervise_co_excluded_using_GRID(double * x_1, double * x_2, double * x_3, unsigned int _array_size,
																																		Three_D_Pattern *& TD_Pattern,
																																		double minimum_population_threshold,
																																		double min_threshold_x,
																																		double max_threshold_x,
																																		double threshold_step,
																																		ostream & err_msg_out)
{
	// ---- Initial values----
	double score = 0.;
	double observed_population_threshold = 0.;

	double threshold_x1 = 0.;
	double threshold_x2 = 0.;
	double threshold_x3 = 0.;

	double p_000 = 0.;
	double p_001 = 0.;
	double p_010 = 0.;
	double p_011 = 0.;
	double p_100 = 0.;
	double p_101 = 0.;
	double p_110 = 0.;
	double p_111 = 0.;

	//----

	double th_1 = 0.;	// thresholds
	double th_2 = 0.;
	double th_3 = 0.;

	// ---- calculating max so no calculations can be performed for thresholds above max values

	double max_x_1 = x_1[0];
	double max_x_2 = x_2[0];
	double max_x_3 = x_3[0];

	unsigned int i;
	for (i = 0; i<_array_size; i++)
	{
		if (max_x_1 < x_1[i]) max_x_1 = x_1[i];
		if (max_x_2 < x_2[i]) max_x_2 = x_2[i];
		if (max_x_3 < x_3[i]) max_x_3 = x_3[i];
	}


	//--------------------- Main loop

	for (th_1 = min_threshold_x; th_1 <= max_threshold_x; th_1 += threshold_step)
	{
		if (th_1 > max_x_1) break;

		for (th_2 = min_threshold_x; th_2 <= max_threshold_x; th_2 += threshold_step)
		{
			if (th_2 > max_x_2) break;

			for (th_3 = min_threshold_x; th_3 <= max_threshold_x; th_3 += threshold_step)
			{
				if (th_3 > max_x_3) break;

				Stat_and_Sort::Three_D_Probabilities(x_1, x_2, x_3, _array_size, th_1, th_2, th_3, p_000, p_001, p_010, p_011, p_100, p_101, p_110, p_111, err_msg_out);

				if (p_111 < minimum_population_threshold || p_001 < minimum_population_threshold || p_010 < minimum_population_threshold) continue;

				if (score < p_111 + p_001 + p_010 + p_000)  // !!!!!!!!!!!!!!!!!!!!!!!!!!!
				{
					score = p_111 + p_001 + p_010 + p_000;
					threshold_x1 = th_1;
					threshold_x2 = th_2;
					threshold_x3 = th_3;
					observed_population_threshold = min(p_111, min(p_001, p_010));
				}
			}
		}
	}

	//  ---------------- final output -------------

	Stat_and_Sort::Three_D_Probabilities(x_1, x_2, x_3, _array_size, threshold_x1, threshold_x2, threshold_x3, p_000, p_001, p_010, p_011, p_100, p_101, p_110, p_111, err_msg_out);

	TD_Pattern = new Three_D_Pattern();	// Pattern is created, but not filed with all the data

	TD_Pattern->score = score;
	TD_Pattern->population_threshold = observed_population_threshold;
	TD_Pattern->threshold_x1 = threshold_x1;
	TD_Pattern->threshold_x2 = threshold_x2;
	TD_Pattern->threshold_x3 = threshold_x3;

	TD_Pattern->p_000 = p_000;
	TD_Pattern->p_001 = p_001;
	TD_Pattern->p_010 = p_010;
	TD_Pattern->p_011 = p_011;
	TD_Pattern->p_100 = p_100;
	TD_Pattern->p_101 = p_101;
	TD_Pattern->p_110 = p_110;
	TD_Pattern->p_111 = p_111;

	return true;
}
//-------------------------------------------------------------------------------------------------------------
//-------------------------------------------------------------------------------------------------------------
//-------------------------------------------------------------------------------------------------------------
//-------------------------------------------------------------------------------------------------------------

bool Three_D_Patterns_Toolbox::find_3D_Type_2_Co_Exclusion_Score_ONLY_using_GRID(double * x_1, double * x_2, double * x_3, unsigned int _array_size,
																										double & score,
																										double minimum_population_threshold,
																										double min_threshold_x,
																										double max_threshold_x,
																										double threshold_step,
																										ostream & err_msg_out)
{
	// ---- Initial values----
	score = 0.;

	double p_000 = 0.;
	double p_001 = 0.;
	double p_010 = 0.;
	double p_011 = 0.;
	double p_100 = 0.;
	double p_101 = 0.;
	double p_110 = 0.;
	double p_111 = 0.;

	//----

	double th_1 = 0.;	// thresholds
	double th_2 = 0.;
	double th_3 = 0.;

	// ---- calculating max so no calculations can be performed for thresholds above max values

	double max_x_1 = x_1[0];
	double max_x_2 = x_2[0];
	double max_x_3 = x_3[0];

	unsigned int i;
	for (i = 0; i<_array_size; i++)
	{
		if (max_x_1 < x_1[i]) max_x_1 = x_1[i];
		if (max_x_2 < x_2[i]) max_x_2 = x_2[i];
		if (max_x_3 < x_3[i]) max_x_3 = x_3[i];
	}


	//--------------------- Main loop

	for (th_1 = min_threshold_x; th_1 <= max_threshold_x; th_1 += threshold_step)
	{
		if (th_1 > max_x_1) break;

		for (th_2 = min_threshold_x; th_2 <= max_threshold_x; th_2 += threshold_step)
		{
			if (th_2 > max_x_2) break;

			for (th_3 = min_threshold_x; th_3 <= max_threshold_x; th_3 += threshold_step)
			{
				if (th_3 > max_x_3) break;

				Stat_and_Sort::Three_D_Probabilities(x_1, x_2, x_3, _array_size, th_1, th_2, th_3, p_000, p_001, p_010, p_011, p_100, p_101, p_110, p_111, err_msg_out);

				if (p_011 < minimum_population_threshold || p_101 < minimum_population_threshold || p_110 < minimum_population_threshold) continue;

				if (score < p_011 + p_101 + p_110 + p_000 + p_001 + p_010 + p_100)
					score = p_011 + p_101 + p_110 + p_000 + p_001 + p_010 + p_100;
			}
		}
	}
	return true;
}


//-------------------------------------------------------------------------------------------------------------

bool Three_D_Patterns_Toolbox::find_3D_Type_2_Co_Exclusion_Score_using_GRID(double * x_1, double * x_2, double * x_3, unsigned int _array_size,
																						Three_D_Pattern *& TD_Pattern,
																						double minimum_population_threshold,
																						double min_threshold_x,
																						double max_threshold_x,
																						double threshold_step,
																						ostream & err_msg_out)
{
	// ---- Initial values----
	double score = 0.;
	double observed_population_threshold = 0.;

	double threshold_x1 = 0.;
	double threshold_x2 = 0.;
	double threshold_x3 = 0.;

	double p_000 = 0.;
	double p_001 = 0.;
	double p_010 = 0.;
	double p_011 = 0.;
	double p_100 = 0.;
	double p_101 = 0.;
	double p_110 = 0.;
	double p_111 = 0.;

	//----

	double th_1 = 0.;	// thresholds
	double th_2 = 0.;
	double th_3 = 0.;

	// ---- calculating max so no calculations can be performed for thresholds above max values

	double max_x_1 = x_1[0];
	double max_x_2 = x_2[0];
	double max_x_3 = x_3[0];

	unsigned int i;
	for (i = 0; i<_array_size; i++)
	{
		if (max_x_1 < x_1[i]) max_x_1 = x_1[i];
		if (max_x_2 < x_2[i]) max_x_2 = x_2[i];
		if (max_x_3 < x_3[i]) max_x_3 = x_3[i];
	}


	//--------------------- Main loop

	for (th_1 = min_threshold_x; th_1 <= max_threshold_x; th_1 += threshold_step)
	{
		if (th_1 > max_x_1) break;

		for (th_2 = min_threshold_x; th_2 <= max_threshold_x; th_2 += threshold_step)
		{
			if (th_2 > max_x_2) break;

			for (th_3 = min_threshold_x; th_3 <= max_threshold_x; th_3 += threshold_step)
			{
				if (th_3 > max_x_3) break;

				Stat_and_Sort::Three_D_Probabilities(x_1, x_2, x_3, _array_size, th_1, th_2, th_3, p_000, p_001, p_010, p_011, p_100, p_101, p_110, p_111, err_msg_out);

				if (p_011 < minimum_population_threshold || p_101 < minimum_population_threshold || p_110 < minimum_population_threshold) continue;

				if (score < p_011 + p_101 + p_110 + p_000 + p_001 + p_010 + p_100) 
				{
					score = p_011 + p_101 + p_110 + p_000 + p_001 + p_010 + p_100;
					threshold_x1 = th_1;
					threshold_x2 = th_2;
					threshold_x3 = th_3;
					observed_population_threshold = min(p_011, min(p_101, p_110));
				}
			}
		}
	}

	//  ---------------- final output -------------

	Stat_and_Sort::Three_D_Probabilities(x_1, x_2, x_3, _array_size, threshold_x1, threshold_x2, threshold_x3, p_000, p_001, p_010, p_011, p_100, p_101, p_110, p_111, err_msg_out);

	TD_Pattern = new Three_D_Pattern();	// Pattern is created, but not filed with all the data

	TD_Pattern->score = score;
	TD_Pattern->population_threshold = observed_population_threshold;
	TD_Pattern->threshold_x1 = threshold_x1;
	TD_Pattern->threshold_x2 = threshold_x2;
	TD_Pattern->threshold_x3 = threshold_x3;

	TD_Pattern->p_000 = p_000;
	TD_Pattern->p_001 = p_001;
	TD_Pattern->p_010 = p_010;
	TD_Pattern->p_011 = p_011;
	TD_Pattern->p_100 = p_100;
	TD_Pattern->p_101 = p_101;
	TD_Pattern->p_110 = p_110;
	TD_Pattern->p_111 = p_111;

	return true;
}
//-------------------------------------------------------------------------------------------------------------

//-------------------------------------------------------------------------------------------------------------
//-------------------------------------------------------------------------------------------------------------
//-------------------------------------------------------------------------------------------------------------
//-------------------------------------------------------------------------------------------------------------
//-------------------------------------------------------------------------------------------------------------
#endif //_THREE_D_PATTERNS_TOOLBOX_H_
```
