## Supplementary material for "Identification of Complex Multidimensional Patterns in Microbial Communities": Two_D_Pattern.h

```
/*
THE SOFTWARE IS PROVIDED "AS IS", WITHOUT WARRANTY OF ANY KIND, EXPRESS OR IMPLIED, INCLUDING BUT NOT LIMITED TO THE WARRANTIES OF
MERCHANTABILITY, FITNESS FOR A PARTICULAR PURPOSE AND NONINFRINGEMENT. IN NO EVENT SHALL THE AUTHORS OR COPYRIGHT HOLDERS BE LIABLE FOR
ANY CLAIM, DAMAGES OR OTHER LIABILITY, WHETHER IN AN ACTION OF CONTRACT, TORT OR OTHERWISE, ARISING FROM, OUT OF OR IN CONNECTION WITH
THE SOFTWARE OR THE USE OR OTHER DEALINGS IN THE SOFTWARE.
*/
#ifndef _Two_D_PATTERN_H_
#define _Two_D_PATTERN_H_


#include <iostream>
#include <fstream>
#include <stdlib.h>
#include <random>
#include <time.h>

#include "OTU_Profile.h"

using namespace std;

class Two_D_Pattern
{

public:

	unsigned int pattern_type; // 1=> co-presence; 2=> co-exclusion; 3=>one way relation.  0=> nothing (default)

	unsigned int OTU_Index_1;	// OTU index in the OTU_Table object
	unsigned int OTU_Index_2;

	OTU_Profile * OTU_p_1;
	OTU_Profile * OTU_p_2;

	double score;					// real (observed) value
	double population_threshold;	// real (observed) value

	double threshold_x1;
	double threshold_x2;
	double p_00;
	double p_01;
	double p_10;
	double p_11;

	double x1_p_0;
	double x1_p_1;

	double x2_p_0;
	double x2_p_1;

// -------------------------------

	Two_D_Pattern();	// creates EMPTY pattern. All zeros
	~Two_D_Pattern();
};
//-------------------------------------------------------------------------------------------------------------
//-------------------------------------------------------------------------------------------------------------
//-------------------------------------------------------------------------------------------------------------

Two_D_Pattern::Two_D_Pattern()
{
	pattern_type = 0;
	OTU_Index_1 = 0;
	OTU_Index_2 = 0;
	
	OTU_p_1=NULL;
	OTU_p_2=NULL;

	score=0.;					
	population_threshold=0.;

	threshold_x1=0.;
	threshold_x2=0.;
	p_00=0.;
	p_01=0.;
	p_10=0.;
	p_11=0.;
	
	x1_p_0=0.;
	x1_p_1=0.;

	x2_p_0=0.;
	x2_p_1 = 0.;

}
//-------------------------------------------------------------------------------------------------------------

Two_D_Pattern::~Two_D_Pattern()
{
	if (OTU_p_1 != NULL) delete OTU_p_1;
	if (OTU_p_2 != NULL) delete OTU_p_2;
}
//-------------------------------------------------------------------------------------------------------------
//-------------------------------------------------------------------------------------------------------------
//-------------------------------------------------------------------------------------------------------------
//-------------------------------------------------------------------------------------------------------------
//-------------------------------------------------------------------------------------------------------------
#endif //Two_D_Pattern
```
