## Supplementary material for "Identification of Complex Multidimensional Patterns in Microbial Communities": Two_D_Patterns_Pipelines.h

```
/*
THE SOFTWARE IS PROVIDED "AS IS", WITHOUT WARRANTY OF ANY KIND, EXPRESS OR IMPLIED, INCLUDING BUT NOT LIMITED TO THE WARRANTIES OF
MERCHANTABILITY, FITNESS FOR A PARTICULAR PURPOSE AND NONINFRINGEMENT. IN NO EVENT SHALL THE AUTHORS OR COPYRIGHT HOLDERS BE LIABLE FOR
ANY CLAIM, DAMAGES OR OTHER LIABILITY, WHETHER IN AN ACTION OF CONTRACT, TORT OR OTHERWISE, ARISING FROM, OUT OF OR IN CONNECTION WITH
THE SOFTWARE OR THE USE OR OTHER DEALINGS IN THE SOFTWARE.
*/
#ifndef _TWO_D_PATTERNS_PIPELINES_H_
#define _TWO_D_PATTERNS_PIPELINES_H_

# include <iostream>
# include <fstream>

#include "Shuffle.h"
#include "Stat_and_Sort.h"

#include "OTU_Table.h"
#include "OTU_Profile.h"
#include "Two_D_Pattern.h"
#include "Array_of_Two_D_Patterns.h"

#include "Two_D_Patterns_Toolbox.h"

using namespace std;

class Two_D_Patterns_Pipelines
{
	
public:	
    static void Make_Tables_of_Number_of_Patterns_for_each_Score_and_Presence_Threshold(char * otu_file,
                                                                                        char * copresence_matrix_file,
                                                                                        char * coexclusion_matrix_file,
                                                                                        char * oneway_matrix_file);

	static void find_2D_Co_Presence_patterns(char * OTU_file_name, char * out_file_name, char * out_file_name_long, 
											double min_threshold = 0., double max_threshold = 0.005, double threshold_step = 0.0005,
											double min_population_threshold = 0.1, double max_population_threshold = 0.2, double population_threshold_step = 0.01,
											double min_score = 0.95, double max_score = 1.,	double score_step = 0.01);


	static void find_2D_Co_Exclusion_patterns(char * OTU_file_name, char * out_file_name, char * out_file_name_long,
											double min_threshold = 0., double max_threshold = 0.005, double threshold_step = 0.0005,
											double min_population_threshold = 0.1, double max_population_threshold = 0.2, double population_threshold_step = 0.01,
											double min_score = 0.95, double max_score = 1., double score_step = 0.01);


	static void find_2D_One_Way_Relation_patterns(char * OTU_file_name, char * out_file_name, char * out_file_name_long,
											double min_threshold = 0., double max_threshold = 0.005, double threshold_step = 0.0005,
											double min_population_threshold = 0.1, double max_population_threshold = 0.2, double population_threshold_step = 0.01,
											double min_score = 0.95, double max_score = 1., double score_step = 0.01);

};

//-------------------------------------------------------------------------------------------------------------
//-------------------------------------------------------------------------------------------------------------
//-------------------------------------------------------------------------------------------------------------

void Two_D_Patterns_Pipelines::find_2D_Co_Presence_patterns(char * OTU_file_name, char * out_file_name, char * out_file_name_long,
															double min_threshold, double max_threshold, double threshold_step,
															double min_population_threshold, double max_population_threshold, double population_threshold_step,
															double min_score, double max_score, double score_step)
{
	Shuffle::initialize();
	
	
	OTU_Table * t = new OTU_Table(OTU_file_name);
	t->re_normalize();
	


	
	OTU_Table * t_random = new OTU_Table(OTU_file_name);


	t_random->shuffle_inside_OTUs();

	t_random->re_normalize();
	

	ofstream out_f;
	out_f.open(out_file_name);
	ofstream out_f_long;
	out_f_long.open(out_file_name_long);


	double * OTU_1 = new double[t->number_of_samples];
	double * OTU_2 = new double[t->number_of_samples];


	double population_threshold;
	double real_data_score_threshold;

	double score;

	unsigned int number_of_scores = (unsigned int)((max_score - min_score) / score_step) + 1;

	unsigned int * number_of_patterns_for_given_score = new unsigned int[number_of_scores];

	Array_of_Two_D_Patterns * patterns_array = new Array_of_Two_D_Patterns();

	bool flag;
	unsigned int patterns_counter;
	unsigned int smallest_score_threshold_index;
	Two_D_Pattern * pattern = NULL;
	unsigned int i;;
	unsigned int ii;
	unsigned int iii;
	for (population_threshold = min_population_threshold; population_threshold < max_population_threshold; population_threshold += population_threshold_step)
	{
		for (iii = 0; iii < number_of_scores; iii++) number_of_patterns_for_given_score[iii] = 0;
		flag = false;

		


		for (i = 0; i < t_random->number_of_OTUs - 1; i++)
		{
			

			for (ii = i + 1; ii < t_random->number_of_OTUs; ii++)
			{
				t_random->get_OTU_profile(OTU_1, i);
				t_random->get_OTU_profile(OTU_2, ii);
				
				Two_D_Patterns_Toolbox::find_Co_Pesence_Score_ONLY_using_GRID(OTU_1, OTU_2, t_random->number_of_samples, score,	population_threshold,min_threshold, max_threshold, threshold_step);

				if (score >= min_score)
				{
					for (iii = 0; iii < number_of_scores; iii++)
						if (score >= min_score + score_step * iii) number_of_patterns_for_given_score[iii]++;
				}
				if (number_of_patterns_for_given_score[number_of_scores - 1] > 0) { flag = true;  break; }
			}
			if (flag) break; 
		}
		if (flag) continue; 

		smallest_score_threshold_index = number_of_scores;
		for (iii = 0; iii < number_of_scores; iii++)
		{
			if (number_of_patterns_for_given_score[iii] == 0)
			{
				smallest_score_threshold_index = iii;
				break;
			}
		}


		if (smallest_score_threshold_index == number_of_scores) continue; // There are no zeros. No reason to try to run real data

// run real data...

		real_data_score_threshold = min_score + score_step * smallest_score_threshold_index;

		

		patterns_counter = 0;

		for (i = 0; i < t->number_of_OTUs - 1; i++)
		{
//			

			for (ii = i + 1; ii < t->number_of_OTUs; ii++)
			{
				t->get_OTU_profile(OTU_1, i);
				t->get_OTU_profile(OTU_2, ii);

				Two_D_Patterns_Toolbox::find_Co_Pesence_Score_using_GRID(OTU_1, OTU_2, t->number_of_samples, pattern, population_threshold,min_threshold, max_threshold, threshold_step);

				if (pattern->score >= real_data_score_threshold)
				{
					pattern->OTU_Index_1 = i;
					pattern->OTU_Index_2 = ii;

					if (patterns_array->is_Pattern_Present(pattern))
					{
						
						delete pattern;
						pattern = NULL;
					}
					else
					{
						patterns_array->add_Pattern(pattern);
						
					}
				}
				else
				{
					delete pattern;
					pattern = NULL;
				}

			}
		}

		if (smallest_score_threshold_index == 0) break;	// no patterns in random data;

	}

	// final output....

	OTU_Profile * p;
	for (i = 0; i < patterns_array->number_of_patterns; i++)
	{
		t->get_OTU_profile(p, patterns_array->pattern[i]->OTU_Index_1);
		patterns_array->pattern[i]->OTU_p_1 = p;
		t->get_OTU_profile(p, patterns_array->pattern[i]->OTU_Index_2);
		patterns_array->pattern[i]->OTU_p_2 = p;
		patterns_array->pattern[i]->pattern_type = 1;
	}

	//patterns_array->show_Statistics();
	patterns_array->show_Statistics(out_f);


	patterns_array->show_All(out_f_long);

	out_f.close();
	out_f_long.close();

	delete t;
	delete t_random;

	delete[] number_of_patterns_for_given_score;
	delete[] OTU_1;
	delete[] OTU_2;

	delete patterns_array;
}
//-------------------------------------------------------------------------------------------------------------


void Two_D_Patterns_Pipelines::find_2D_Co_Exclusion_patterns(char * OTU_file_name, char * out_file_name, char * out_file_name_long,
															double min_threshold, double max_threshold, double threshold_step,
															double min_population_threshold, double max_population_threshold, double population_threshold_step,
															double min_score, double max_score, double score_step)
{
	Shuffle::initialize();

	//cout << "\n=============== Real ========================\n";
	OTU_Table * t = new OTU_Table(OTU_file_name);
	t->re_normalize();
	//t->show_Statistics();


	//cout << "\n=============== Random ========================\n";
	OTU_Table * t_random = new OTU_Table(OTU_file_name);


	t_random->shuffle_inside_OTUs();

	t_random->re_normalize();
	//t_random->show_Statistics();
	//cout << "++++++++++++++++++++++++++++++++++++++++++++++++++++++++\n";


	ofstream out_f;
	out_f.open(out_file_name);
	ofstream out_f_long;
	out_f_long.open(out_file_name_long);


	double * OTU_1 = new double[t->number_of_samples];
	double * OTU_2 = new double[t->number_of_samples];


	// ----- Parameters

	double population_threshold;
	double real_data_score_threshold;

	double score;

	unsigned int number_of_scores = (unsigned int)((max_score - min_score) / score_step) + 1;

	unsigned int * number_of_patterns_for_given_score = new unsigned int[number_of_scores];

	Array_of_Two_D_Patterns * patterns_array = new Array_of_Two_D_Patterns();

	bool flag;
	unsigned int patterns_counter;
	unsigned int smallest_score_threshold_index;
	Two_D_Pattern * pattern = NULL;
	unsigned int i;;
	unsigned int ii;
	unsigned int iii;
	for (population_threshold = min_population_threshold; population_threshold < max_population_threshold; population_threshold += population_threshold_step)
	{
		for (iii = 0; iii < number_of_scores; iii++) number_of_patterns_for_given_score[iii] = 0;
		flag = false;	// do we ned to exit this population threshold iteration?

		//cout << "population_threshold=" << population_threshold << endl;


		for (i = 0; i < t_random->number_of_OTUs - 1; i++)
		{
			//cout << i << "\t";

			for (ii = i + 1; ii < t_random->number_of_OTUs; ii++)
			{
				t_random->get_OTU_profile(OTU_1, i);
				t_random->get_OTU_profile(OTU_2, ii);

				Two_D_Patterns_Toolbox::find_Co_Exclusion_Score_ONLY_using_GRID(OTU_1, OTU_2, t_random->number_of_samples, score, population_threshold,min_threshold, max_threshold, threshold_step);

				if (score >= min_score)
				{
					for (iii = 0; iii < number_of_scores; iii++)
						if (score >= min_score + score_step * iii) number_of_patterns_for_given_score[iii]++;
				}
				if (number_of_patterns_for_given_score[number_of_scores - 1] > 0) { flag = true; break; }
			}
			if (flag) break;
		}

		if (flag) continue;

// are there any zeros on the number_of_patterns_for_given_score?

		smallest_score_threshold_index = number_of_scores;
		for (iii = 0; iii < number_of_scores; iii++)
		{
			if (number_of_patterns_for_given_score[iii] == 0)
			{
				smallest_score_threshold_index = iii;
				break;
			}
		}

//		cout << "\nsmallest_score_threshold_index=" << smallest_score_threshold_index << endl;


		if (smallest_score_threshold_index == number_of_scores) continue; // There are no zeros. No reason to try to run real data

// run real data...

		real_data_score_threshold = min_score + score_step * smallest_score_threshold_index;

		//cout << "\n real_data_score_threshold=" << real_data_score_threshold << endl;

		patterns_counter = 0;

		for (i = 0; i < t->number_of_OTUs - 1; i++)
		{
//			cout << i << "\treal\t";

			for (ii = i + 1; ii < t->number_of_OTUs; ii++)
			{
				t->get_OTU_profile(OTU_1, i);
				t->get_OTU_profile(OTU_2, ii);

				Two_D_Patterns_Toolbox::find_Co_Exclusion_Score_using_GRID(OTU_1, OTU_2, t->number_of_samples, pattern, population_threshold, min_threshold, max_threshold, threshold_step);

				if (pattern->score >= real_data_score_threshold)
				{
					pattern->OTU_Index_1 = i;
					pattern->OTU_Index_2 = ii;

					if (patterns_array->is_Pattern_Present(pattern))
					{
						//cout << "duplicated pattern!\n";
						delete pattern;
						pattern = NULL;
					}
					else
					{
						patterns_array->add_Pattern(pattern);
						//cout << "patterns_counter=" << patterns_counter++ << endl;
					}
				}
				else
				{
					delete pattern;
					pattern = NULL;
				}

			}
		}

		if (smallest_score_threshold_index == 0) break;	// no patterns in random data;

	}

// final output....

	OTU_Profile * p;
	for (i = 0; i < patterns_array->number_of_patterns; i++)
	{
		t->get_OTU_profile(p, patterns_array->pattern[i]->OTU_Index_1);
		patterns_array->pattern[i]->OTU_p_1 = p;
		t->get_OTU_profile(p, patterns_array->pattern[i]->OTU_Index_2);
		patterns_array->pattern[i]->OTU_p_2 = p;
		patterns_array->pattern[i]->pattern_type = 2;
	}

	//patterns_array->show_Statistics();
	patterns_array->show_Statistics(out_f);


	patterns_array->show_All(out_f_long);

	out_f.close();
	out_f_long.close();

	delete t;
	delete t_random;

	delete[] number_of_patterns_for_given_score;
	delete[] OTU_1;
	delete[] OTU_2;

	delete patterns_array;
}
//-------------------------------------------------------------------------------------------------------------


void Two_D_Patterns_Pipelines::find_2D_One_Way_Relation_patterns(char * OTU_file_name, char * out_file_name, char * out_file_name_long,
															double min_threshold, double max_threshold, double threshold_step,
															double min_population_threshold, double max_population_threshold, double population_threshold_step,
															double min_score, double max_score, double score_step)
{
	Shuffle::initialize();

	//cout << "\n=============== Real ========================\n";
	OTU_Table * t = new OTU_Table(OTU_file_name);
	t->re_normalize();
//	t->show_Statistics();


	//cout << "\n=============== Random ========================\n";
	OTU_Table * t_random = new OTU_Table(OTU_file_name);

//	t_random->shuffle_All();/////////////////////////////////////////////////////////////////////////////
	t_random->shuffle_inside_OTUs();

	t_random->re_normalize();
//	t_random->show_Statistics();
	//cout << "++++++++++++++++++++++++++++++++++++++++++++++++++++++++\n";


	ofstream out_f;
	out_f.open(out_file_name);
	ofstream out_f_long;
	out_f_long.open(out_file_name_long);


	double * OTU_1 = new double[t->number_of_samples];
	double * OTU_2 = new double[t->number_of_samples];

	double population_threshold;
	double real_data_score_threshold;

	double score;

	unsigned int number_of_scores = (unsigned int)((max_score - min_score) / score_step) + 1;

	unsigned int * number_of_patterns_for_given_score = new unsigned int[number_of_scores];

	Array_of_Two_D_Patterns * patterns_array = new Array_of_Two_D_Patterns();

	bool flag;
	unsigned int patterns_counter;
	unsigned int smallest_score_threshold_index;
	Two_D_Pattern * pattern = NULL;
	unsigned int i;;
	unsigned int ii;
	unsigned int iii;
	for (population_threshold = min_population_threshold; population_threshold < max_population_threshold; population_threshold += population_threshold_step)
	{
		for (iii = 0; iii < number_of_scores; iii++) number_of_patterns_for_given_score[iii] = 0;
		flag = false;	// do we ned to exit this population threshold iteration?

		//cout << "population_threshold=" << population_threshold << endl;


		for (i = 0; i < t_random->number_of_OTUs; i++)
		{
			//cout << i << "\t";

			for (ii = 0; ii < t_random->number_of_OTUs; ii++)
			{
				if (i == ii) continue;

				t_random->get_OTU_profile(OTU_1, i);
				t_random->get_OTU_profile(OTU_2, ii);

				Two_D_Patterns_Toolbox::find_One_Way_Relation_X1_need_X2_Score_ONLY_using_GRID(OTU_1, OTU_2, t_random->number_of_samples, score, population_threshold,
																								min_threshold, max_threshold, threshold_step);

				if (score >= min_score)
				{
					for (iii = 0; iii < number_of_scores; iii++)
						if (score >= min_score + score_step * iii) number_of_patterns_for_given_score[iii]++;
				}
//				cout << number_of_patterns_for_given_score[number_of_scores - 1] << endl;
				if (number_of_patterns_for_given_score[number_of_scores - 1] > 0) { flag = true; break; }
			}
			if (flag) break;
		}

		if (flag) continue;


// are there any zeros on the number_of_patterns_for_given_score?

		smallest_score_threshold_index = number_of_scores;
		for (iii = 0; iii < number_of_scores; iii++)
		{
			if (number_of_patterns_for_given_score[iii] == 0)
			{
				smallest_score_threshold_index = iii;
				break;
			}
		}

//		cout << "\nsmallest_score_threshold_index=" << smallest_score_threshold_index << endl;


		if (smallest_score_threshold_index == number_of_scores) continue; // There are no zeros. No reason to try to run real data

// run real data...

		real_data_score_threshold = min_score + score_step * smallest_score_threshold_index;

		//cout << "\n real_data_score_threshold=" << real_data_score_threshold << endl;

		patterns_counter = 0;

		for (i = 0; i < t->number_of_OTUs - 1; i++)
		{
//			cout << i << "\treal\t";

			for (ii = i + 1; ii < t->number_of_OTUs; ii++)
			{
				t->get_OTU_profile(OTU_1, i);
				t->get_OTU_profile(OTU_2, ii);

				Two_D_Patterns_Toolbox::find_One_Way_Relation_X1_need_X2_Score_using_GRID(OTU_1, OTU_2, t->number_of_samples, pattern, population_threshold,
																							min_threshold, max_threshold, threshold_step);

				if (pattern->score >= real_data_score_threshold)
				{
					pattern->OTU_Index_1 = i;
					pattern->OTU_Index_2 = ii;

					if (patterns_array->is_Pattern_Present(pattern))
					{
						//cout << "duplicated pattern!\n";
						delete pattern;
						pattern = NULL;
					}
					else
					{
						patterns_array->add_Pattern(pattern);
						//cout << "patterns_counter=" << patterns_counter++ << endl;
					}
				}
				else
				{
					delete pattern;
					pattern = NULL;
				}
			}
		}
		if (smallest_score_threshold_index == 0) break;	// no patterns in random data;
	}

	// final output....

	OTU_Profile * p;
	for (i = 0; i < patterns_array->number_of_patterns; i++)
	{
		t->get_OTU_profile(p, patterns_array->pattern[i]->OTU_Index_1);
		patterns_array->pattern[i]->OTU_p_1 = p;
		t->get_OTU_profile(p, patterns_array->pattern[i]->OTU_Index_2);
		patterns_array->pattern[i]->OTU_p_2 = p;
		patterns_array->pattern[i]->pattern_type = 3;
	}

	//patterns_array->show_Statistics();
	patterns_array->show_Statistics(out_f);


	patterns_array->show_All(out_f_long);

	out_f.close();
	out_f_long.close();

	delete t;
	delete t_random;

	delete[] number_of_patterns_for_given_score;
	delete[] OTU_1;
	delete[] OTU_2;

	delete patterns_array;

}
//-------------------------------------------------------------------------------------------------------------

// Dependance of patterns scores from min_population_threshold (min_population_threshold X score matrix)
void Two_D_Patterns_Pipelines::Make_Tables_of_Number_of_Patterns_for_each_Score_and_Presence_Threshold(char * otu_file,
                                                                                                       char * copresence_matrix_file,
                                                                                                       char * coexclusion_matrix_file,
                                                                                                       char * oneway_matrix_file
                                                                                                       )

{

	Shuffle::initialize();


    char * OTU_file_name = otu_file;


    char * out_file_name_co_presence = copresence_matrix_file;
    char * out_file_name_co_exclusion = coexclusion_matrix_file;
    char * out_file_name_One_Way_Relation_X1_need_X2 = oneway_matrix_file;


	//cout << "\n=============== Real ========================\n";
	OTU_Table * t = new OTU_Table(OTU_file_name);
	t->re_normalize();
	//t->show_Statistics();


//	cout << "\n=============== Random ========================\n";
	OTU_Table * t_random = new OTU_Table(OTU_file_name);

//	t_random->shuffle_All();  ////////////////////////////////////////////////////////////
	t_random->shuffle_inside_OTUs();
	
	
	t_random->re_normalize();
	//t_random->show_Statistics();


	//cout << "++++++++++++++++++++++++++++++++++++++++++++++++++++++++\n";

	double * OTU_1 = new double[t->number_of_samples];
	double * OTU_2 = new double[t->number_of_samples];


	ofstream out_f_co_presence;
	out_f_co_presence.open(out_file_name_co_presence);

	ofstream out_f_co_exclusion;
	out_f_co_exclusion.open(out_file_name_co_exclusion);

	ofstream out_f_One_Way_Relation_X1_need_X2;
	out_f_One_Way_Relation_X1_need_X2.open(out_file_name_One_Way_Relation_X1_need_X2);

	// ----- Parameters

	double min_threshold = 0.000;
	double max_threshold = 0.4;
	double threshold_step = 0.0005;

	double min_population_threshold = 0.05;
	double max_population_threshold = 0.4;
	double population_threshold_step = 0.025;// 0.01;
	double population_threshold;

	double min_score = 0.80;
	double max_score = 1.01;
	double score_step = 0.01;//0.01;

	// -------- variables


	double score;

	unsigned int number_of_scores = (unsigned int)((max_score - min_score) / score_step) + 1;

	unsigned int * co_presence_scores = new unsigned int[number_of_scores];
	unsigned int * co_exclusion_scores = new unsigned int[number_of_scores];
	unsigned int * One_Way_Relation_X1_need_X2_scores = new unsigned int[number_of_scores];

	// ----- Files headerts 

	out_f_co_presence << "REAL\nscore";
	out_f_co_exclusion << "REAL\nscore";
	out_f_One_Way_Relation_X1_need_X2 << "REAL\nscore";

	unsigned int iii;
	for (iii = 0; iii < number_of_scores; iii++)
	{
		out_f_co_presence << "\t" << min_score + score_step * iii;
		out_f_co_exclusion << "\t" << min_score + score_step * iii;
		out_f_One_Way_Relation_X1_need_X2 << "\t" << min_score + score_step * iii;
	}

	out_f_co_presence << "\npopulation_threshold\n";
	out_f_co_exclusion << "\npopulation_threshold\n";
	out_f_One_Way_Relation_X1_need_X2 << "\npopulation_threshold\n";

	// ============ Main loop REAL =========================

	bool calculate_co_presence = true;
	bool calculate_co_exclusion = true;
	bool calculate_One_Way_Relation = true;

	unsigned int i;
	unsigned int ii;
	for (population_threshold = min_population_threshold; population_threshold < max_population_threshold; population_threshold += population_threshold_step)
	{
		for (iii = 0; iii < number_of_scores; iii++)
		{
			co_presence_scores[iii] = 0;
			co_exclusion_scores[iii] = 0;
			One_Way_Relation_X1_need_X2_scores[iii] = 0;
		}

		//cout << "population_threshold=" << population_threshold << endl;

		out_f_co_presence << population_threshold;
		out_f_co_exclusion << population_threshold;
		out_f_One_Way_Relation_X1_need_X2 << population_threshold;

		for (i = 0; i < t->number_of_OTUs; i++)
		{
			//		if (t->OTUs_max[i] < 0.001) continue;
			//cout << i << "\t";

			for (ii = i + 1; ii < t->number_of_OTUs; ii++)
			{
				t->get_OTU_profile(OTU_1, i);
				t->get_OTU_profile(OTU_2, ii);

//----------- Co-presence
				if (calculate_co_presence)
				{
					Two_D_Patterns_Toolbox::find_Co_Pesence_Score_ONLY_using_GRID(OTU_1, OTU_2, t->number_of_samples, score, population_threshold,
						min_threshold, max_threshold, threshold_step);

					if (score >= min_score)
					{
						for (iii = 0; iii < number_of_scores; iii++)
							if (score >= min_score + score_step * iii) co_presence_scores[iii]++;
					}
				}

//---------- Co-exclusion
				if (calculate_co_exclusion)
				{
					Two_D_Patterns_Toolbox::find_Co_Exclusion_Score_ONLY_using_GRID(OTU_1, OTU_2, t->number_of_samples, score, population_threshold,
						min_threshold, max_threshold, threshold_step);

					if (score >= min_score)
					{
						for (iii = 0; iii < number_of_scores; iii++)
							if (score >= min_score + score_step * iii) co_exclusion_scores[iii]++;
					}
				}
			}

		}
//-----------One_Way_Relation_X1_need_X2


		for (i = 0; i < t->number_of_OTUs; i++)
		{
			//cout << i << "\t";

			for (ii = 0; ii < t->number_of_OTUs; ii++)
			{
				if (i == ii) continue;
				t->get_OTU_profile(OTU_1, i);
				t->get_OTU_profile(OTU_2, ii);

//-----------One_Way_Relation_X1_need_X2

				if (calculate_One_Way_Relation)
				{
					Two_D_Patterns_Toolbox::find_One_Way_Relation_X1_need_X2_Score_ONLY_using_GRID(OTU_1, OTU_2, t->number_of_samples, score, population_threshold,
						min_threshold, max_threshold, threshold_step);

					if (score >= min_score)
					{
						for (iii = 0; iii < number_of_scores; iii++)
							if (score >= min_score + score_step * iii) One_Way_Relation_X1_need_X2_scores[iii]++;
					}
				}
			}
		}


		for (iii = 0; iii < number_of_scores; iii++)
		{
			out_f_co_presence << "\t" << co_presence_scores[iii];
			out_f_co_exclusion << "\t" << co_exclusion_scores[iii];
			out_f_One_Way_Relation_X1_need_X2 << "\t" << One_Way_Relation_X1_need_X2_scores[iii];
		}


		out_f_co_presence << endl;
		out_f_co_exclusion << endl;
		out_f_One_Way_Relation_X1_need_X2 << endl;

		if (co_presence_scores[0] == 0) calculate_co_presence = false;
		if (co_exclusion_scores[0] == 0) calculate_co_exclusion = false;
		if (One_Way_Relation_X1_need_X2_scores[0] == 0) calculate_One_Way_Relation = false;
		cout << endl;
	}


	// ============ Main loop RANDOM =========================

	// ----- Files headerts 

	out_f_co_presence << "\n\n\n\n\nRANDOM\nscore";
	out_f_co_exclusion << "\n\n\n\n\nRANDOM\nscore";
	out_f_One_Way_Relation_X1_need_X2 << "\n\n\n\n\nRANDOM\nscore";


	for (iii = 0; iii < number_of_scores; iii++)
	{
		out_f_co_presence << "\t" << min_score + score_step * iii;
		out_f_co_exclusion << "\t" << min_score + score_step * iii;
		out_f_One_Way_Relation_X1_need_X2 << "\t" << min_score + score_step * iii;
	}

	out_f_co_presence << "\npopulation_threshold\n";
	out_f_co_exclusion << "\npopulation_threshold\n";
	out_f_One_Way_Relation_X1_need_X2 << "\npopulation_threshold\n";

	t_random->show_Statistics();

	calculate_co_presence = true;
	calculate_co_exclusion = true;
	calculate_One_Way_Relation = true;


	for (population_threshold = min_population_threshold; population_threshold < max_population_threshold; population_threshold += population_threshold_step)
	{
		for (iii = 0; iii < number_of_scores; iii++)
		{
			co_presence_scores[iii] = 0;
			co_exclusion_scores[iii] = 0;
			One_Way_Relation_X1_need_X2_scores[iii] = 0;
		}

		//cout << "population_threshold=" << population_threshold << endl;

		out_f_co_presence << population_threshold;
		out_f_co_exclusion << population_threshold;
		out_f_One_Way_Relation_X1_need_X2 << population_threshold;

		for (i = 0; i < t_random->number_of_OTUs; i++)
		{
			//cout << i << "\t";
			for (ii = i + 1; ii < t_random->number_of_OTUs; ii++)
			{
				t_random->get_OTU_profile(OTU_1, i);
				t_random->get_OTU_profile(OTU_2, ii);

//----------- Co-presence
				if (calculate_co_presence)
				{
					Two_D_Patterns_Toolbox::find_Co_Pesence_Score_ONLY_using_GRID(OTU_1, OTU_2, t_random->number_of_samples, score, population_threshold,
						min_threshold, max_threshold, threshold_step);

					if (score >= min_score)
					{
						for (iii = 0; iii < number_of_scores; iii++)
							if (score >= min_score + score_step * iii) co_presence_scores[iii]++;
					}
				}

//----------- Co-exclusion

				if (calculate_co_exclusion)
				{
					Two_D_Patterns_Toolbox::find_Co_Exclusion_Score_ONLY_using_GRID(OTU_1, OTU_2, t_random->number_of_samples, score, population_threshold,
						min_threshold, max_threshold, threshold_step);

					if (score >= min_score)
					{
						for (iii = 0; iii < number_of_scores; iii++)
							if (score >= min_score + score_step * iii) co_exclusion_scores[iii]++;
					}
				}
			}
		}

//-----------One_Way_Relation_X1_need_X2
		for (i = 0; i < t_random->number_of_OTUs; i++)
		{
			//cout << i << "\t";

			for (ii = 0; ii < t_random->number_of_OTUs; ii++)
			{
				if (i == ii) continue;
				t_random->get_OTU_profile(OTU_1, i);
				t_random->get_OTU_profile(OTU_2, ii);

				if (calculate_One_Way_Relation)
				{
					Two_D_Patterns_Toolbox::find_One_Way_Relation_X1_need_X2_Score_ONLY_using_GRID(OTU_1, OTU_2, t_random->number_of_samples, score, population_threshold,
						min_threshold, max_threshold, threshold_step);

					if (score >= min_score)
					{
						for (iii = 0; iii < number_of_scores; iii++)
							if (score >= min_score + score_step * iii) One_Way_Relation_X1_need_X2_scores[iii]++;
					}
				}
			}
		}

		for (iii = 0; iii < number_of_scores; iii++)
		{
			out_f_co_presence << "\t" << co_presence_scores[iii];
			out_f_co_exclusion << "\t" << co_exclusion_scores[iii];
			out_f_One_Way_Relation_X1_need_X2 << "\t" << One_Way_Relation_X1_need_X2_scores[iii];
		}

		out_f_co_presence << endl;
		out_f_co_exclusion << endl;
		out_f_One_Way_Relation_X1_need_X2 << endl;

		if (co_presence_scores[0] == 0) calculate_co_presence = false;
		if (co_exclusion_scores[0] == 0) calculate_co_exclusion = false;
		if (One_Way_Relation_X1_need_X2_scores[0] == 0) calculate_One_Way_Relation = false;


		//cout << endl;
	}

	out_f_co_presence.close();
	out_f_co_exclusion.close();
	out_f_One_Way_Relation_X1_need_X2.close();

	delete t;
	delete t_random;

	delete[] OTU_1;
	delete[] OTU_2;

	delete[] co_presence_scores;
	delete[] co_exclusion_scores;
	delete[] One_Way_Relation_X1_need_X2_scores;

}
//-------------------------------------------------------------------------------------------------------------


#endif //_TWO_D_PATTERNS_PIPELINES_H_
```
