## Supplementary material for "Identification of Complex Multidimensional Patterns in Microbial Communities": Two_D_Patterns_Toolbox.h

```
/*
THE SOFTWARE IS PROVIDED "AS IS", WITHOUT WARRANTY OF ANY KIND, EXPRESS OR IMPLIED, INCLUDING BUT NOT LIMITED TO THE WARRANTIES OF
MERCHANTABILITY, FITNESS FOR A PARTICULAR PURPOSE AND NONINFRINGEMENT. IN NO EVENT SHALL THE AUTHORS OR COPYRIGHT HOLDERS BE LIABLE FOR
ANY CLAIM, DAMAGES OR OTHER LIABILITY, WHETHER IN AN ACTION OF CONTRACT, TORT OR OTHERWISE, ARISING FROM, OUT OF OR IN CONNECTION WITH
THE SOFTWARE OR THE USE OR OTHER DEALINGS IN THE SOFTWARE.
*/
#ifndef _TWO_D_PATTERNS_TOOLBOX_H_
#define _TWO_D_PATTERNS_TOOLBOX_H_


#include <iostream>
#include "Shuffle.h"
#include "Stat_and_Sort.h"
#include "Two_D_Pattern.h"
#include "Three_D_Pattern.h"

using namespace std;

//-------------------------------------------------------------------------------------------------------------
//	This class is designed to calculate various entropy and mutual information scores.
// NOTE: All calculations performed using base 2 log: ===> log2(2.0)==1.0;
// NOTE: All calculations perforned for binay (true(1)/false(0)) values;
//-------------------------------------------------------------------------------------------------------------


class Two_D_Patterns_Toolbox
{
public:

// ============================ Patterns Specific... 2D only
//-------------------------------------------------------------------------------------------------------------
//--------------------------    PATTERN SPECIFIC SCORES    ----------------------------------------------------
//-------------------------------------------------------------------------------------------------------------
// --- Search for patterns specific score.  --------------------------------------------------------------------
// ------------------------------------------------------------------------------------------------------------
// For each pattern the "score" is calculated by maximizing the total percent of points associated (belonging) 
// to the pattern under condition that the percentage of points in each sector (00,01,10,11) is above the "minimum_population_threshold"
// 
// -------------------------------------------------------------------------------------------------------

// NOTE: score is calculated as p_00+p_11
	static bool find_Co_Pesence_Score_ONLY_using_GRID(double * x_1, double * x_2, unsigned int _array_size, 
													double & score,
													double minimum_population_threshold,
													double min_threshold_x = 0.,
													double max_threshold_x = 0.005,
													double threshold_step =  0.0001,
													ostream & err_msg_out = cout);

	static bool find_Co_Pesence_Score_using_GRID(double * x_1, double * x_2, unsigned int _array_size,
												Two_D_Pattern *& TD_Pattern,	
												double minimum_population_threshold,
												double min_threshold_x = 0.,
												double max_threshold_x = 0.005,
												double threshold_step =	 0.0001,
												ostream & err_msg_out = cout);


//-----------------------------------------------------------------------------------------------------
// NOTE: score is calculated as p_01+p_10
	static bool find_Co_Exclusion_Score_ONLY_using_GRID(double * x_1, double * x_2, unsigned int _array_size,
													double & score,
													double minimum_population_threshold,
													double min_threshold_x = 0.,
													double max_threshold_x = 0.005,
													double threshold_step =  0.0001,
													ostream & err_msg_out = cout);


	static bool find_Co_Exclusion_Score_using_GRID(double * x_1, double * x_2, unsigned int _array_size,
													Two_D_Pattern *& TD_Pattern,
													double minimum_population_threshold,
													double min_threshold_x = 0.,
													double max_threshold_x = 0.3,
													double threshold_step = 0.0001,
													ostream & err_msg_out = cout);


//-----------------------------------------------------------------------------------------------------
// NOTE: This relation means that X1 is present only if X2 is present. meaning p_10=0;
//	score is calculated as p_00+p11+p_01
	static bool find_One_Way_Relation_X1_need_X2_Score_ONLY_using_GRID(double * x_1, double * x_2, unsigned int _array_size,
													double & score,
													double minimum_population_threshold,
													double min_threshold_x = 0.,
													double max_threshold_x = 0.3,
													double threshold_step = 0.0001,
													ostream & err_msg_out = cout);

	static bool find_One_Way_Relation_X1_need_X2_Score_using_GRID(double * x_1, double * x_2, unsigned int _array_size,
																	Two_D_Pattern *& TD_Pattern,
																	double minimum_population_threshold,
																	double min_threshold_x = 0.,
																	double max_threshold_x = 0.3,
																	double threshold_step = 0.0001,
																	ostream & err_msg_out = cout);

};
//-------------------------------------------------------------------------------------------------------------
//-------------------------------------------------------------------------------------------------------------
//-------------------------------------------------------------------------------------------------------------


// NOTE: This relation means that X1 is present only if X2 is present. meaning p_10=0;
//	score is calculated as p_00+p11+p_01
bool Two_D_Patterns_Toolbox::find_One_Way_Relation_X1_need_X2_Score_ONLY_using_GRID(double * x_1, double * x_2, unsigned int _array_size,
																								double & score,
																								double minimum_population_threshold,
																								double min_threshold_x,
																								double max_threshold_x,
																								double threshold_step,
																								ostream & err_msg_out)
{
	// ---- Initial values----

	score = 0.;

	double p_00 = 0.;
	double p_01 = 0.;
	double p_10 = 0.;
	double p_11 = 0.;
	//----

	double th_1=0.;	// thresholds
	double th_2=0.;

	// ---- calculating max so no calculations can be performed for thresholds above max values

	double max_x_1 = x_1[0];
	double max_x_2 = x_2[0];

	unsigned int i;
	for (i = 0; i<_array_size; i++)
	{
		if (max_x_1 < x_1[i]) max_x_1 = x_1[i];
		if (max_x_2 < x_2[i]) max_x_2 = x_2[i];
	}


	//--------------------- Main loop

	for (th_1 = min_threshold_x; th_1 <= max_threshold_x; th_1 += threshold_step)
	{
		if (th_1 > max_x_1) break;

		for (th_2 = min_threshold_x; th_2 <= max_threshold_x; th_2 += threshold_step)
		{
			if (th_2 > max_x_2) break;

			Stat_and_Sort::Two_D_Probabilities(x_1, x_2, _array_size, th_1, th_2, p_00, p_01, p_10, p_11, err_msg_out);

			if (p_01 < minimum_population_threshold || p_11 < minimum_population_threshold || p_00 < minimum_population_threshold) continue;

			if (score < p_01 + p_11 + p_00)
				score = p_01 + p_11 + p_00;
		}
	}
	return true;
}
//-------------------------------------------------------------------------------------------------------------

bool Two_D_Patterns_Toolbox::find_One_Way_Relation_X1_need_X2_Score_using_GRID(double * x_1, double * x_2, unsigned int _array_size,
																							Two_D_Pattern *& TD_Pattern,
																							double minimum_population_threshold,
																							double min_threshold_x,
																							double max_threshold_x,
																							double threshold_step,
																							ostream & err_msg_out)
{
// ---- Initial values----
	double score = 0.;
	double observed_population_threshold = 0.;

	double threshold_x1 = 0.;
	double threshold_x2 = 0.;
	double p_00 = 0.;
	double p_01 = 0.;
	double p_10 = 0.;
	double p_11 = 0.;

	double x1_p_0 = 0.;
	double x1_p_1 = 0.;

	double x2_p_0 = 0.;
	double x2_p_1 = 0.;

//----

	double th_1 = 0.;	// thresholds
	double th_2 = 0.;


// ---- calculating max so no calculations can be performed for thresholds above max values

	double max_x_1 = x_1[0];
	double max_x_2 = x_2[0];

	unsigned int i;
	for (i = 0; i<_array_size; i++)
	{
		if (max_x_1 < x_1[i]) max_x_1 = x_1[i];
		if (max_x_2 < x_2[i]) max_x_2 = x_2[i];
	}


	//--------------------- Main loop

	for (th_1 = min_threshold_x; th_1 <= max_threshold_x; th_1 += threshold_step)
	{
		if (th_1 > max_x_1) break;

		for (th_2 = min_threshold_x; th_2 <= max_threshold_x; th_2 += threshold_step)
		{
			if (th_2 > max_x_2) break;

			Stat_and_Sort::Two_D_Probabilities(x_1, x_2, _array_size, th_1, th_2, p_00, p_01, p_10, p_11, err_msg_out);

			if (p_01 < minimum_population_threshold || p_11 < minimum_population_threshold || p_00 < minimum_population_threshold) continue;

			if (score < p_01 + p_11 + p_00)
			{
				score = p_01 + p_11 + p_00;
				threshold_x1 = th_1;
				threshold_x2 = th_2;
				observed_population_threshold = min(p_11, min(p_01, p_00));
			}
		}
	}

	//  ---------------- final output -------------

	Stat_and_Sort::Two_D_Probabilities(x_1, x_2, _array_size, threshold_x1, threshold_x2, p_00, p_01, p_10, p_11, err_msg_out);
	Stat_and_Sort::One_D_Probabilities(x_1, _array_size, threshold_x1, x1_p_0, x1_p_1);
	Stat_and_Sort::One_D_Probabilities(x_2, _array_size, threshold_x2, x2_p_0, x2_p_1);

	TD_Pattern = new Two_D_Pattern();	// Pattern is created, but not filed with all the data...

	TD_Pattern->score = score;
	TD_Pattern->population_threshold = observed_population_threshold;
	TD_Pattern->threshold_x1 = threshold_x1;
	TD_Pattern->threshold_x2 = threshold_x2;
	TD_Pattern->p_00 = p_00;
	TD_Pattern->p_01 = p_01;
	TD_Pattern->p_10 = p_10;
	TD_Pattern->p_11 = p_11;
	TD_Pattern->x1_p_0 = x1_p_0;
	TD_Pattern->x1_p_1 = x1_p_1;
	TD_Pattern->x2_p_0 = x2_p_0;
	TD_Pattern->x2_p_1 = x2_p_1;

	return true;
}
//-------------------------------------------------------------------------------------------------------------

// NOTE: score is calculate as p_01+p_10
bool Two_D_Patterns_Toolbox::find_Co_Exclusion_Score_ONLY_using_GRID(double * x_1, double * x_2, unsigned int _array_size,
																		double & score,
																		double minimum_population_threshold,
																		double min_threshold_x,
																		double max_threshold_x,
																		double threshold_step,
																		ostream & err_msg_out)
{
	// ---- Initial values----

	score = 0.;

	double p_00 = 0.;
	double p_01 = 0.;
	double p_10 = 0.;
	double p_11 = 0.;

	//----

	double th_1;	// thresholds
	double th_2;

	// ---- calculating max so no calculations can be performed for thresholds above max values

	double max_x_1 = x_1[0];
	double max_x_2 = x_2[0];

	unsigned int i;
	for (i = 0; i<_array_size; i++)
	{
		if (max_x_1 < x_1[i]) max_x_1 = x_1[i];
		if (max_x_2 < x_2[i]) max_x_2 = x_2[i];
	}


	//--------------------- Main loop

	for (th_1 = min_threshold_x; th_1 <= max_threshold_x; th_1 += threshold_step)
	{
		if (th_1 > max_x_1) break;

		for (th_2 = min_threshold_x; th_2 <= max_threshold_x; th_2 += threshold_step)
		{
			if (th_2 > max_x_2) break;

			Stat_and_Sort::Two_D_Probabilities(x_1, x_2, _array_size, th_1, th_2, p_00, p_01, p_10, p_11, err_msg_out);

			if (p_01 < minimum_population_threshold || p_10 < minimum_population_threshold) continue;

			if (score < p_01 + p_10 + p_00)
				score = p_01 + p_10 + p_00;
		}
	}
	return true;
}
//-------------------------------------------------------------------------------------------------------------

// NOTE: score is calculated as p_01+p_10
bool Two_D_Patterns_Toolbox::find_Co_Exclusion_Score_using_GRID(double * x_1, double * x_2, unsigned int _array_size,
																			Two_D_Pattern *& TD_Pattern,
																			double minimum_population_threshold,
																			double min_threshold_x,
																			double max_threshold_x,
																			double threshold_step,
																			ostream & err_msg_out)
{
	// ---- Initial values----
	double score = 0.;
	double observed_population_threshold = 0.;

	double threshold_x1 = 0.;
	double threshold_x2 = 0.;
	double p_00 = 0.;
	double p_01 = 0.;
	double p_10 = 0.;
	double p_11 = 0.;

	double x1_p_0 = 0.;
	double x1_p_1 = 0.;

	double x2_p_0 = 0.;
	double x2_p_1 = 0.;

//----

	double th_1 = 0.;	// thresholds
	double th_2 = 0.;

// ---- calculating max so no calculations can be performed for thresholds above max values

	double max_x_1 = x_1[0];
	double max_x_2 = x_2[0];

	unsigned int i;
	for (i = 0; i<_array_size; i++)
	{
		if (max_x_1 < x_1[i]) max_x_1 = x_1[i];
		if (max_x_2 < x_2[i]) max_x_2 = x_2[i];
	}


	//--------------------- Main loop

	for (th_1 = min_threshold_x; th_1 <= max_threshold_x; th_1 += threshold_step)
	{
		if (th_1 > max_x_1) break;

		for (th_2 = min_threshold_x; th_2 <= max_threshold_x; th_2 += threshold_step)
		{
			if (th_2 > max_x_2) break;

			Stat_and_Sort::Two_D_Probabilities(x_1, x_2, _array_size, th_1, th_2, p_00, p_01, p_10, p_11, err_msg_out);

			if (p_01 < minimum_population_threshold || p_10 < minimum_population_threshold) continue;

			if (score < p_01 + p_10 + p_00)  // !!!!!!!!!!!!!!!!!!!!!!!!!!!
			{
				score = p_01 + p_10 + p_00;
				threshold_x1 = th_1;
				threshold_x2 = th_2;
				observed_population_threshold = min(p_10, p_01);
			}
		}
	}

	//  ---------------- final output -------------

	Stat_and_Sort::Two_D_Probabilities(x_1, x_2, _array_size, threshold_x1, threshold_x2, p_00, p_01, p_10, p_11, err_msg_out);
	Stat_and_Sort::One_D_Probabilities(x_1, _array_size, threshold_x1, x1_p_0, x1_p_1);
	Stat_and_Sort::One_D_Probabilities(x_2, _array_size, threshold_x2, x2_p_0, x2_p_1);

	TD_Pattern = new Two_D_Pattern();	// Pattern is created, but not filed with all the data

	TD_Pattern->score = score;
	TD_Pattern->population_threshold = observed_population_threshold;
	TD_Pattern->threshold_x1 = threshold_x1;
	TD_Pattern->threshold_x2 = threshold_x2;
	TD_Pattern->p_00 = p_00;
	TD_Pattern->p_01 = p_01;
	TD_Pattern->p_10 = p_10;
	TD_Pattern->p_11 = p_11;
	TD_Pattern->x1_p_0 = x1_p_0;
	TD_Pattern->x1_p_1 = x1_p_1;
	TD_Pattern->x2_p_0 = x2_p_0;
	TD_Pattern->x2_p_1 = x2_p_1;

	return true;
}
//-------------------------------------------------------------------------------------------------------------

// NOTE: score is calculated as p_00+p_11
bool Two_D_Patterns_Toolbox::find_Co_Pesence_Score_ONLY_using_GRID(double * x_1, double * x_2, unsigned int _array_size,
												double & score,
												double minimum_population_threshold,
												double min_threshold_x,
												double max_threshold_x,
												double threshold_step,
												ostream & err_msg_out)
{
// ---- Initial values----

	score = 0.;

	double p_00 = 0.;
	double p_01 = 0.;
	double p_10 = 0.;
	double p_11 = 0.;
//----

	double th_1;	// thresholds
	double th_2;

// ---- calculating max so no calculations can be performed for thresholds above max values

	double max_x_1 = x_1[0];
	double max_x_2 = x_2[0];

	unsigned int i;
	for (i = 0; i<_array_size; i++)
	{
		if (max_x_1 < x_1[i]) max_x_1 = x_1[i];
		if (max_x_2 < x_2[i]) max_x_2 = x_2[i];
	}


//--------------------- Main loop

	for (th_1 = min_threshold_x; th_1 <= max_threshold_x; th_1 += threshold_step)
	{
		if (th_1 > max_x_1) break;

		for (th_2 = min_threshold_x; th_2 <= max_threshold_x; th_2 += threshold_step)
		{
			if (th_2 > max_x_2) break;

			Stat_and_Sort::Two_D_Probabilities(x_1, x_2, _array_size, th_1, th_2, p_00, p_01, p_10, p_11, err_msg_out);
			
			if (p_00 < minimum_population_threshold || p_11 < minimum_population_threshold) continue;

			if (score < p_00+p_11) score = p_00 + p_11;
		}
	}

	return true;
}
//-------------------------------------------------------------------------------------------------------------

// NOTE: score is calculated as p_00+p_11
bool Two_D_Patterns_Toolbox::find_Co_Pesence_Score_using_GRID(double * x_1, double * x_2, unsigned int _array_size,
																		Two_D_Pattern *& TD_Pattern,
																		double minimum_population_threshold,
																		double min_threshold_x,
																		double max_threshold_x,
																		double threshold_step,
																		ostream & err_msg_out)
{
// ---- Initial values----
	double score = 0.;
	double observed_population_threshold = 0.;

	double threshold_x1 = 0.;
	double threshold_x2 = 0.;
	double p_00 = 0.;
	double p_01 = 0.;
	double p_10 = 0.;
	double p_11 = 0.;

	double x1_p_0 = 0.;
	double x1_p_1 = 0.;

	double x2_p_0 = 0.;
	double x2_p_1 = 0.;

	//----

	double th_1 = 0.;	// thresholds
	double th_2 = 0.;

	// ---- calculating max so no calculations can be performed for thresholds above max values

	double max_x_1 = x_1[0];
	double max_x_2 = x_2[0];

	unsigned int i;
	for (i = 0; i<_array_size; i++)
	{
		if (max_x_1 < x_1[i]) max_x_1 = x_1[i];
		if (max_x_2 < x_2[i]) max_x_2 = x_2[i];
	}


//--------------------- Main loop

	for (th_1 = min_threshold_x; th_1 <= max_threshold_x; th_1 += threshold_step)
	{
		if (th_1 > max_x_1) break;

		for (th_2 = min_threshold_x; th_2 <= max_threshold_x; th_2 += threshold_step)
		{
			if (th_2 > max_x_2) break;

			Stat_and_Sort::Two_D_Probabilities(x_1, x_2, _array_size, th_1, th_2, p_00, p_01, p_10, p_11, err_msg_out);

			if (p_00 < minimum_population_threshold || p_11 < minimum_population_threshold) continue;

			if (score < p_00 + p_11)
			{
				score = p_00 + p_11;
				threshold_x1 = th_1;
				threshold_x2 = th_2;
				observed_population_threshold = min(p_11, p_00);
			}
		}
	}

	//  ---------------- final output -------------

	Stat_and_Sort::Two_D_Probabilities(x_1, x_2, _array_size, threshold_x1, threshold_x2, p_00, p_01, p_10, p_11, err_msg_out);
	Stat_and_Sort::One_D_Probabilities(x_1, _array_size, threshold_x1, x1_p_0, x1_p_1);
	Stat_and_Sort::One_D_Probabilities(x_2, _array_size, threshold_x2, x2_p_0, x2_p_1);

	TD_Pattern = new Two_D_Pattern();	// Pattern is created, but not filed with all the data

	TD_Pattern->score = score;

//	cout <<" TD_Pattern->score = " << TD_Pattern->score << endl;

	TD_Pattern->population_threshold = observed_population_threshold;
	TD_Pattern->threshold_x1 = threshold_x1;
	TD_Pattern->threshold_x2 = threshold_x2;
	TD_Pattern->p_00 = p_00;
	TD_Pattern->p_01 = p_01;
	TD_Pattern->p_10 = p_10;
	TD_Pattern->p_11 = p_11;
	TD_Pattern->x1_p_0 = x1_p_0;
	TD_Pattern->x1_p_1 = x1_p_1;
	TD_Pattern->x2_p_0 = x2_p_0;
	TD_Pattern->x2_p_1 = x2_p_1;

	return true;
}
//-------------------------------------------------------------------------------------------------------------
//-------------------------------------------------------------------------------------------------------------
//-------------------------------------------------------------------------------------------------------------
//-------------------------------------------------------------------------------------------------------------
//-------------------------------------------------------------------------------------------------------------
//-------------------------------------------------------------------------------------------------------------
//-------------------------------------------------------------------------------------------------------------
//-------------------------------------------------------------------------------------------------------------
//-------------------------------------------------------------------------------------------------------------
#endif //_TWO_D_PATTERNS_TOOLBOX_H_
```
