## Supplementary material for "Identification of Complex Multidimensional Patterns in Microbial Communities": Supplementary Materials.pdf


| | |
|---|---|
| Golovko George*+ | 1. Department of Pharmacology and Toxicology, University of Texas Medical Branch–Galveston, Galveston, TX 77555-0144, USA,<br>2. Sealy Center for Structural Biology and Molecular Biophysics, University of Texas Medical Branch–Galveston, Galveston, TX 77555-0144, USA |
| Khanipov Kamil* | 1. Department of Pharmacology and Toxicology, University of Texas Medical Branch–Galveston, Galveston, TX 77555-0144, USA,<br>2. Sealy Center for Structural Biology and Molecular Biophysics, University of Texas Medical Branch–Galveston, Galveston, TX 77555-0144, USA |
| Levent Albayrak | 1. Department of Pharmacology and Toxicology, University of Texas Medical Branch–Galveston, Galveston, TX 77555-0144, USA,<br>2. Sealy Center for Structural Biology and Molecular Biophysics, University of Texas Medical Branch–Galveston, Galveston, TX 77555-0144, USA |
| Fofanov Yuriy | 1. Department of Pharmacology and Toxicology, University of Texas Medical Branch–Galveston, Galveston, TX 77555-0144, USA,<br>2. Sealy Center for Structural Biology and Molecular Biophysics, University of Texas Medical Branch–Galveston, Galveston, TX 77555-0144, USA |

*Authors contributed equally to this manuscript

+To whom correspondence should be addressed

**Abstract**

**Motivation:** Identification of complex relationships within members of microbial communities is key to understand and guide microbial transplantation and provide personalized anti-microbial and probiotic treatments. Since members of a given microbial community can be simultaneously involved in multiple relations that altogether will determine their abundance, not all significant relations between organisms are expected to be manifested as visually uninterrupted patterns and be detected using traditional correlation nor mutual information coefficient based approaches.

**Results:** This manuscript proposes a pattern specific way to quantify the strength and estimate the statistical significance of two-dimensional *co-presence*, *co-exclusion,* and *one-way relations* patterns between abundance profiles of two organisms which can be extended to three or more dimensional patterns. Presented approach can also be extended by including a variety of physical (pH, temperature, oxygen concentration) and biochemical (antimicrobial susceptibility, nutrient and metabolite concentration) variables into the search for multidimensional patterns. The presented approach has been tested using 2,380 microbiome samples from the Human Microbiome Project resulting in body-site specific networks of statistically significant 2D patterns. We also were able to demonstrate the presence of several 3D patterns in the Human Microbiome Project data.

**Availability:** C++ source code for two and three-dimensional patterns, as well as executable files for the Pickle pipeline, are in the attached supplementary materials.

**Contact:**

**Supplementary information:** Supplementary data are available at Bioinformatics online.

**2D Patterns**

**1. Mid Vagina**

**Data summary:** 85 OTUs present across 86 samples were used in the analysis. 15 OTUs involved in 18 statistically significant relationships were observed in the network.

**Figure 1:** 2D patterns identified with minimum pattern score 0.95. Node colors reflect different taxonomy assignments at Phylum level and node sizes are proportional to the average relative abundance of the microorganism across samples. Capital letters inside square brackets represent the lowest taxonomy level identified for each OTU: G (Genus), F (Family), O (Order), C (Class) and P (Phyla). Color of edges indicates relationship type: blue with black arrow (one-way relations), red (co-exclusion), and green (co-presence). See Table S1.xslx for pattern types, scores, OTU taxonomic labels, etc.

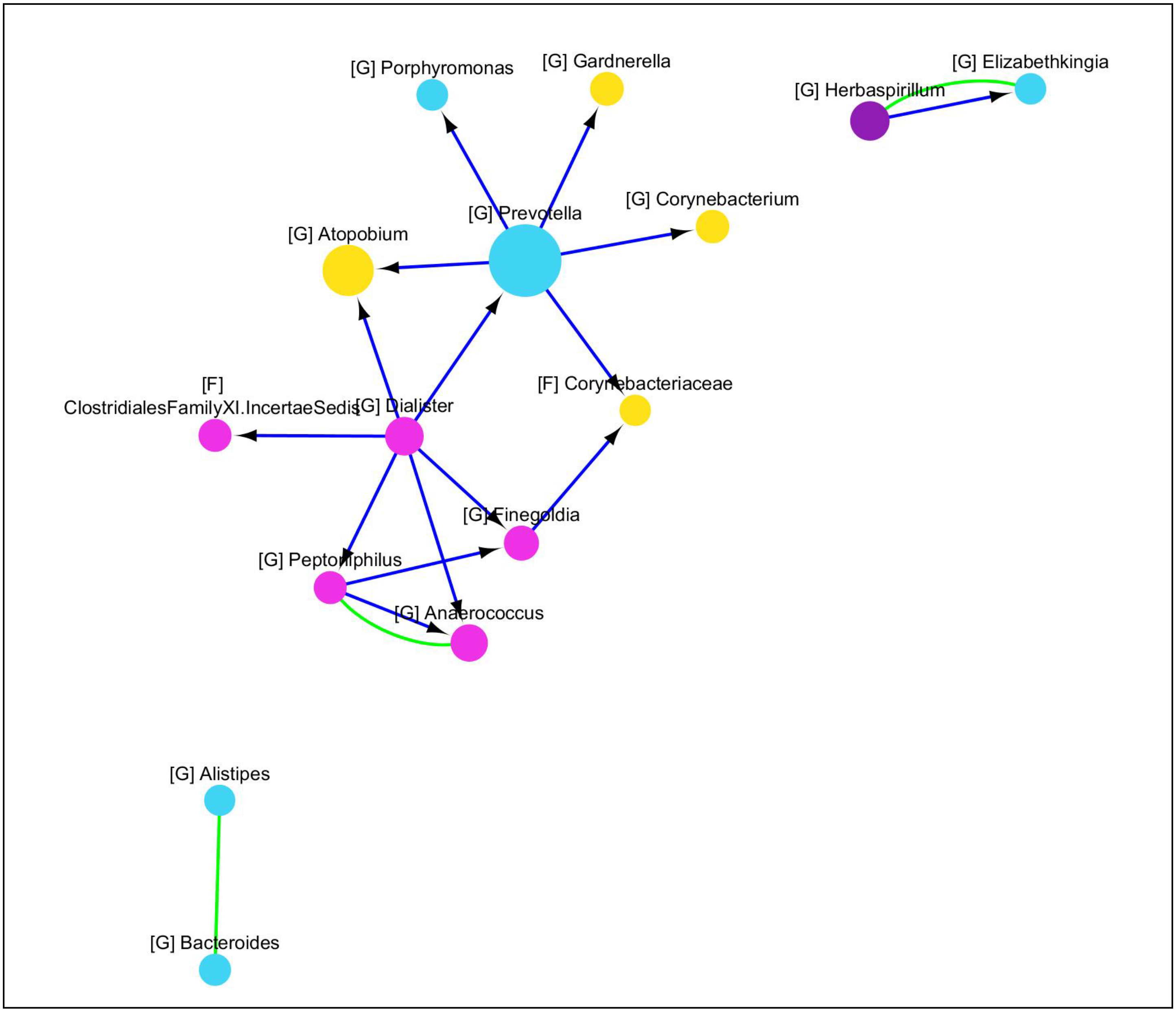

**2. Posterior Fornix**

**Data summary:** 67 OTUs present across 85 samples were used in the analysis. 19 OTUs involved in 26 statistically significant relationships were observed in the network.

**Figure 2:** 2D patterns identified with minimum pattern score 0.95. Node colors reflect different taxonomy assignments at Phylum level and node sizes are proportional to the average relative abundance of the microorganism across samples. Capital letters inside square brackets represent the lowest taxonomy level identified for each OTU: G (Genus), F (Family), O (Order), C (Class) and P (Phyla). Color of edges indicates relationship type: blue with black arrow (one-way relations), red (co-exclusion), and green (co-presence). See Table S2.xslx for pattern types, scores, OTU taxonomic labels, etc.

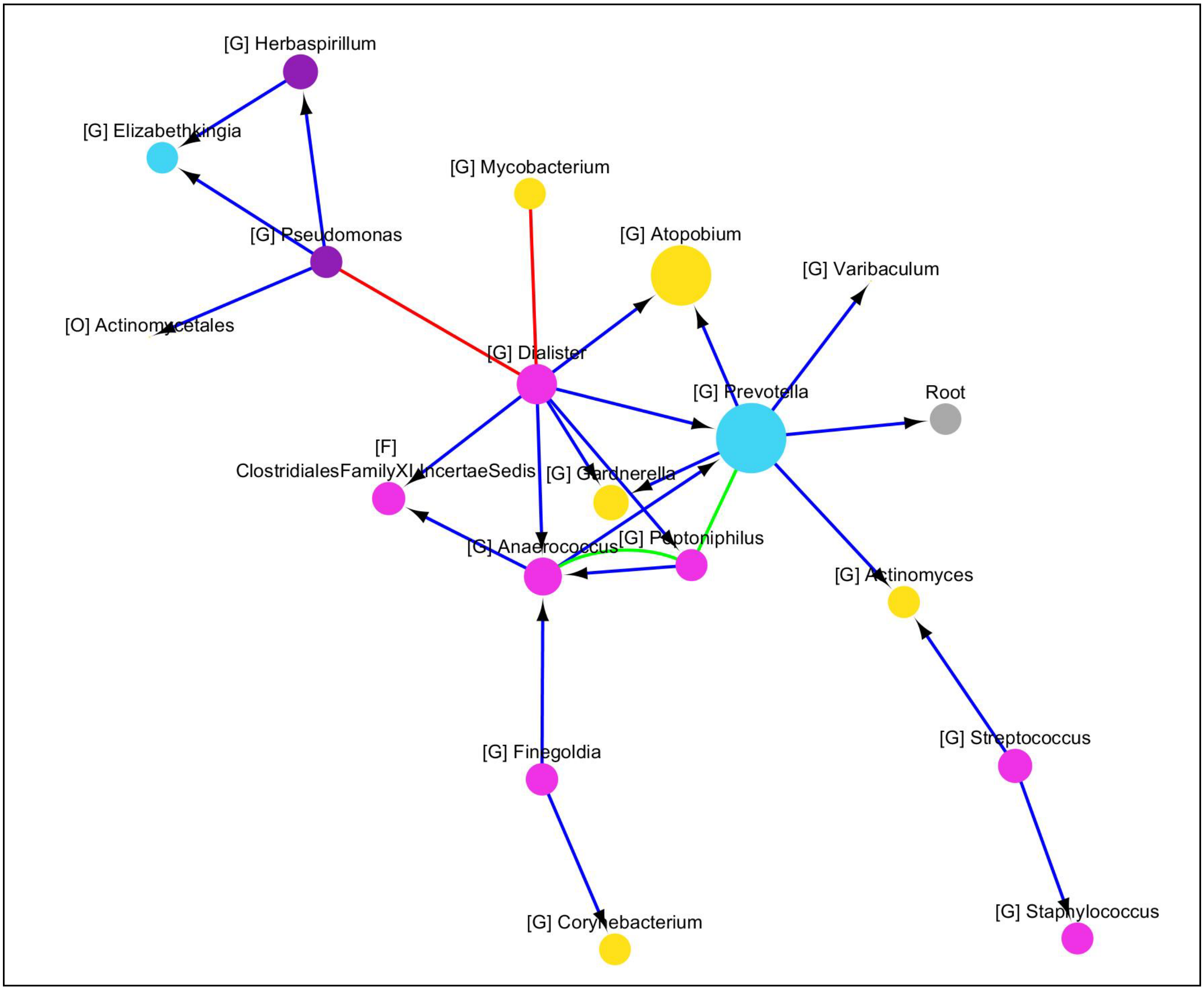

#### 3. Vaginal Introitus

**Data summary:** 96 OTUs present across 85 samples were used in the analysis. 11 OTUs involved in 11 statistically significant relationships were observed in the network.

**Figure 3:** 2D patterns identified with minimum pattern score 0.95. Node colors reflect different taxonomy assignments at Phylum level and node sizes are proportional to the average relative abundance of the microorganism across samples. Capital letters inside square brackets represent the lowest taxonomy level identified for each OTU: G (Genus), F (Family), O (Order), C (Class) and P (Phyla). Color of edges indicates relationship type: blue with black arrow (one-way relations), red (co-exclusion), and green (co-presence). See Table S3.xslx for pattern types, scores, OTU taxonomic labels, etc.

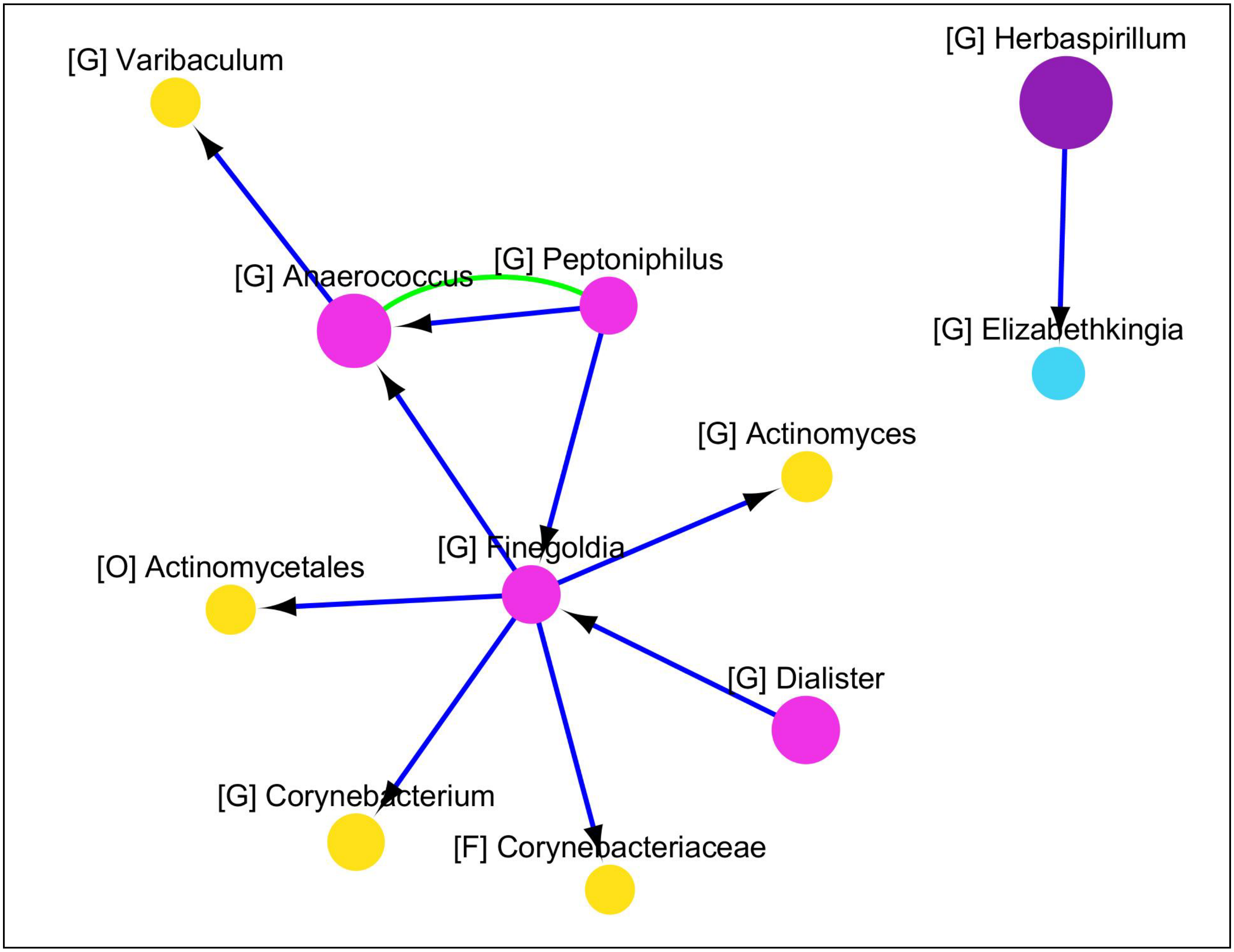

### 4. Stool

**Data summary:** 99 OTUs present across 160 samples were used in the analysis. 26 OTUs involved in 32 statistically significant relationships were observed in the network.

**Figure 4:** 2D patterns identified with minimum pattern score 0.95. Node colors reflect different taxonomy assignments at Phylum level and node sizes are proportional to the average relative abundance of the microorganism across samples. Capital letters inside square brackets represent the lowest taxonomy level identified for each OTU: G (Genus), F (Family), O (Order), C (Class) and P (Phyla). Color of edges indicates relationship type: blue with black arrow (one-way relations), red (co-exclusion), and green (co-presence). See Table S4.xslx for pattern types, scores, OTU taxonomic labels, etc.

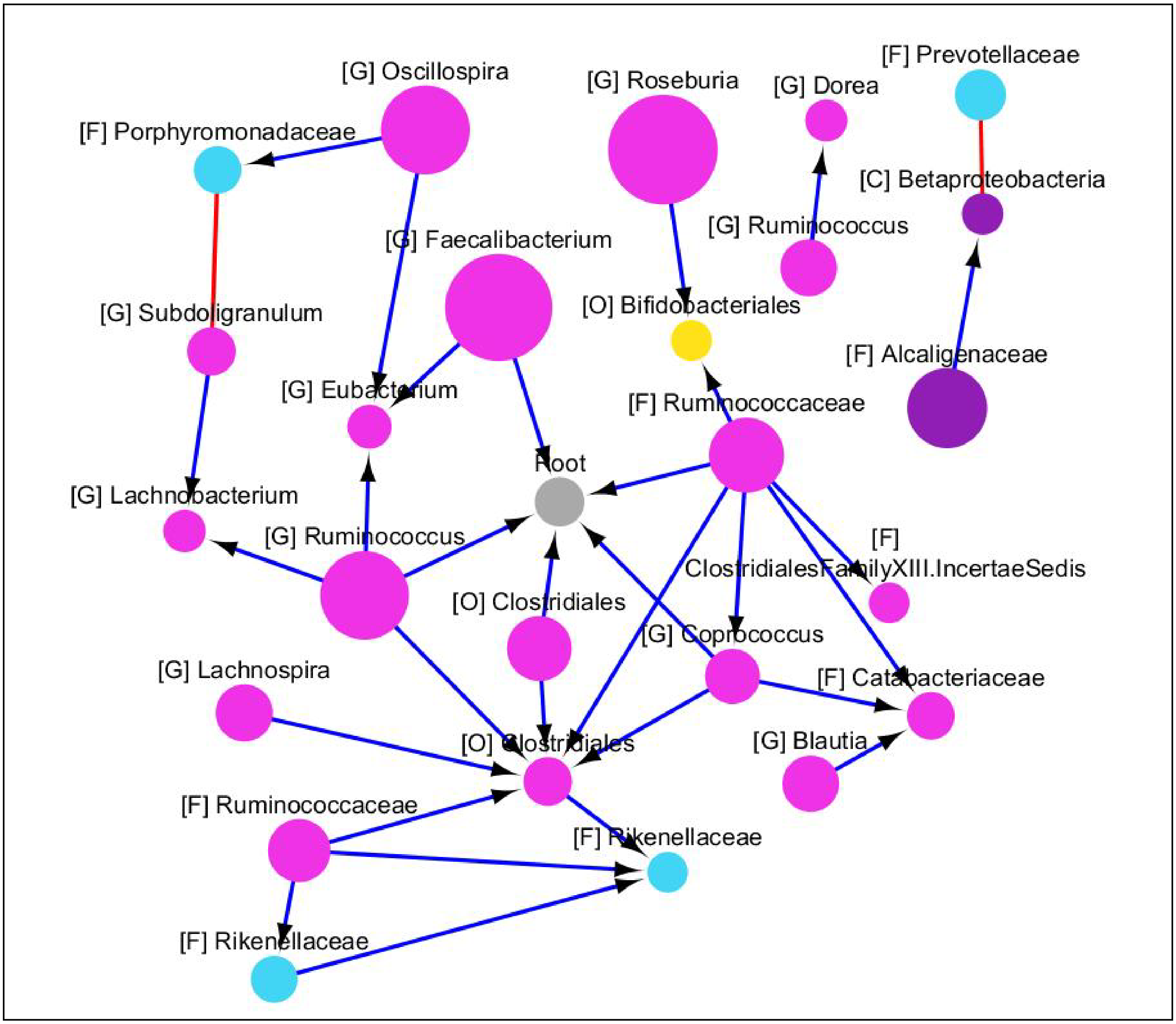

**5. Saliva**

**Data summary:** 113 OTUs present across 130 samples were used in the analysis. 6 OTUs involved in 4 statistically significant relationships were observed in the network.

**Figure 5:** 2D patterns identified with minimum pattern score 0.95. Node colors reflect different taxonomy assignments at Phylum level and node sizes are proportional to the average relative abundance of the microorganism across samples. Capital letters inside square brackets represent the lowest taxonomy level identified for each OTU: G (Genus), F (Family), O (Order), C (Class) and P (Phyla). Color of edges indicates relationship type: blue with black arrow (one-way relations), red (co-exclusion), and green (co-presence). See Table S5.xslx for pattern types, scores, OTU taxonomic labels, etc.

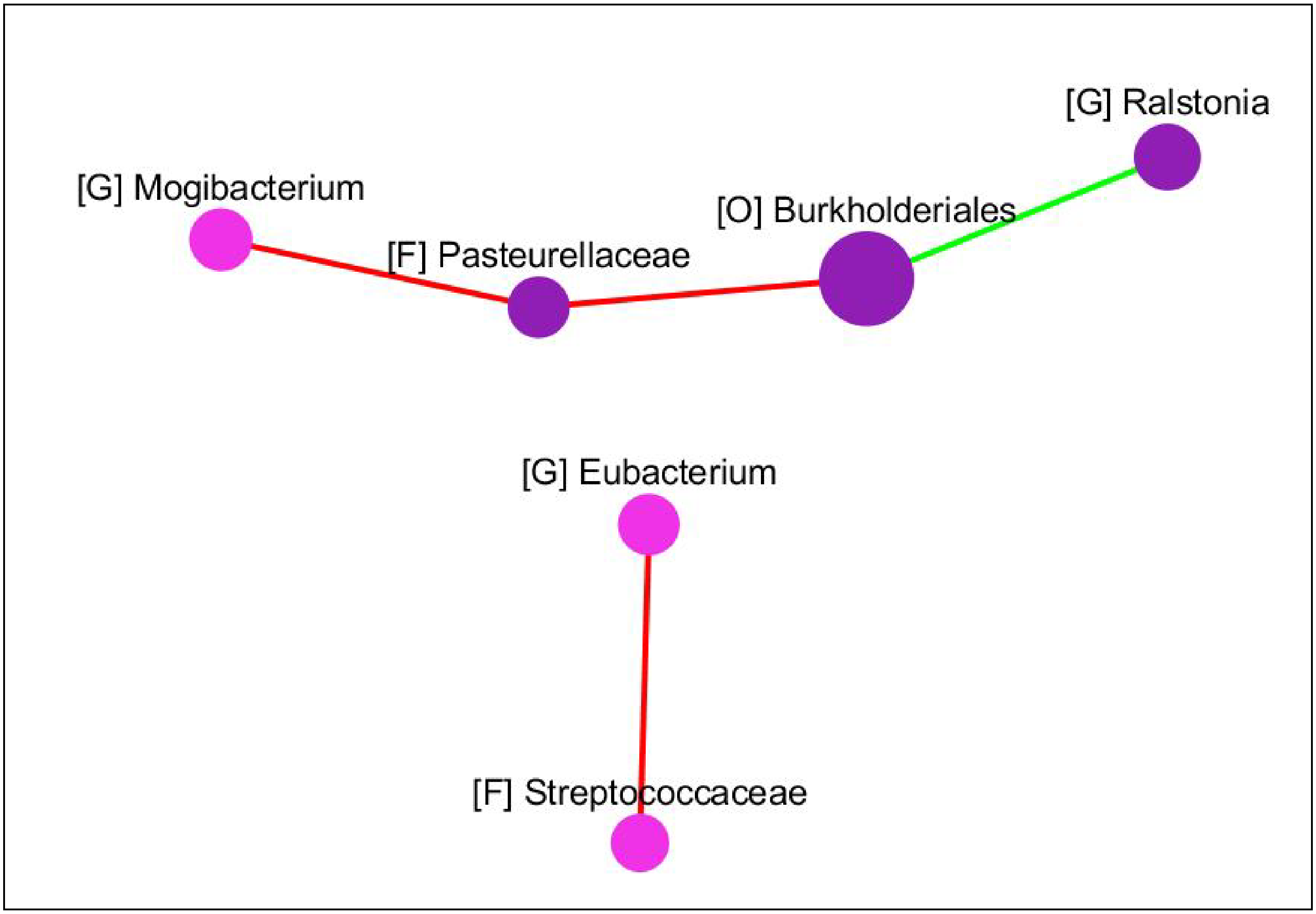

**6. Throat**

**Data summary:** 115 OTUs present across 140 samples were used in the analysis. 4 OTUs involved in 2 statistically significant relationships were observed in the network.

**Figure 6:** 2D patterns identified with minimum pattern score 0.95. Node colors reflect different taxonomy assignments at Phylum level and node sizes are proportional to the average relative abundance of the microorganism across samples. Capital letters inside square brackets represent the lowest taxonomy level identified for each OTU: G (Genus), F (Family), O (Order), C (Class) and P (Phyla). Color of edges indicates relationship type: blue with black arrow (one-way relations), red (co-exclusion), and green (co-presence). See Table S6.xslx for pattern types, scores, OTU taxonomic labels, etc.

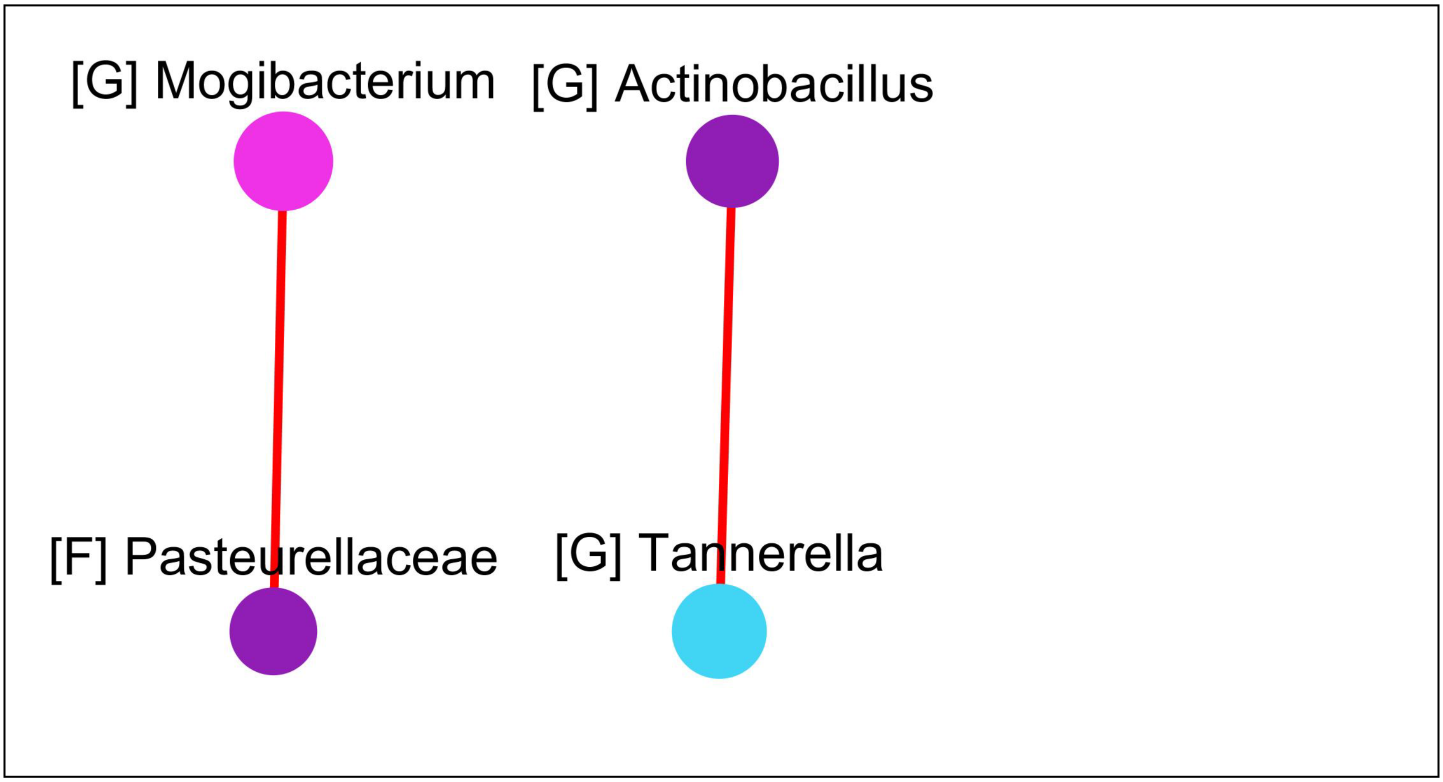

### 7. Tongue Dorsum

**Data summary:** 81 OTUs present across 159 samples were used in the analysis. 35 OTUs involved in 43 statistically significant relationships were observed in the network.

**Figure 7:** 2D patterns identified with minimum pattern score 0.95. Node colors reflect different taxonomy assignments at Phylum level and node sizes are proportional to the average relative abundance of the microorganism across samples. Capital letters inside square brackets represent the lowest taxonomy level identified for each OTU: G (Genus), F (Family), O (Order), C (Class) and P (Phyla). Color of edges indicates relationship type: blue with black arrow (one-way relations), red (co-exclusion), and green (co-presence). See Table S7.xslx for pattern types, scores, OTU taxonomic labels, etc.

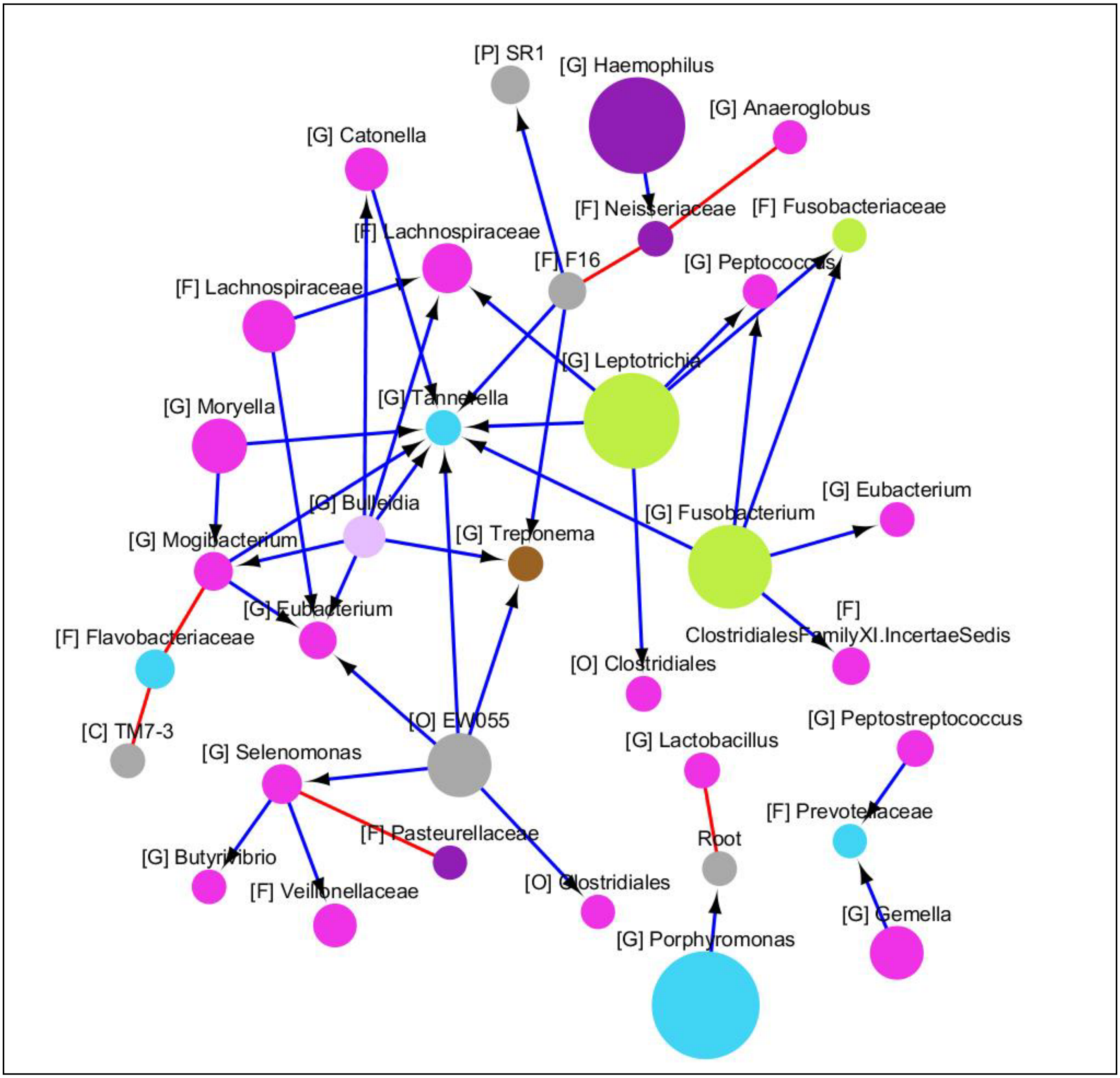

**8. Supragingival Plaque**

**Data summary:** 90 OTUs present across 155 samples were used in the analysis. 35 OTUs involved in 58 statistically significant relationships were observed in the network.

**Figure 8:** 2D patterns identified with minimum pattern score 0.95. Node colors reflect different taxonomy assignments at Phylum level and node sizes are proportional to the average relative abundance of the microorganism across samples. Capital letters inside square brackets represent the lowest taxonomy level identified for each OTU: G (Genus), F (Family), O (Order), C (Class) and P (Phyla). Color of edges indicates relationship type: blue with black arrow (one-way relations), red (co-exclusion), and green (co-presence). See Table S8.xslx for pattern types, scores, OTU taxonomic labels, etc.

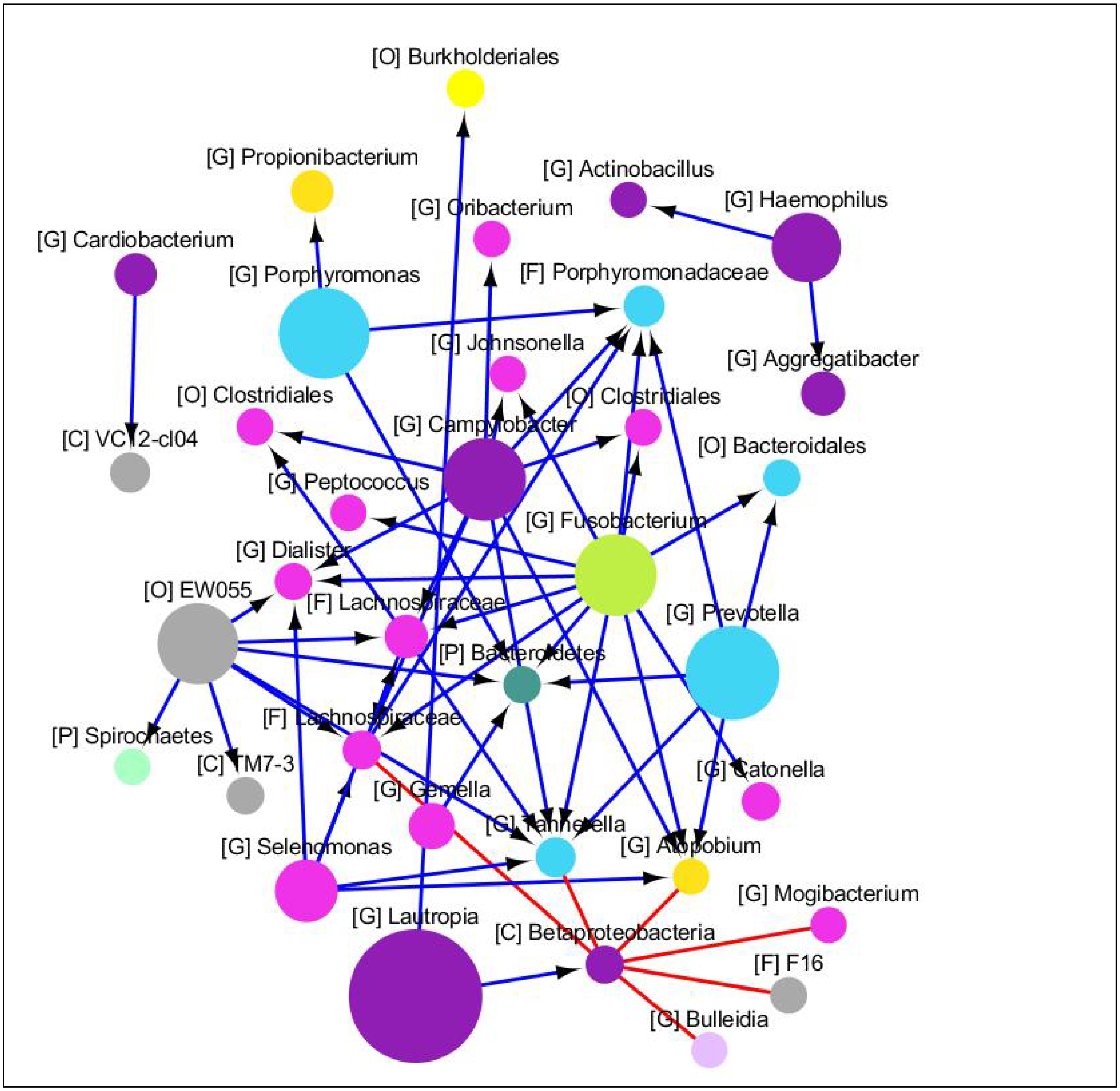

### 9. Subgingival Plaque

**Data summary:** 94 OTUs present across 155 samples were used in the analysis. 33 OTUs involved in 33 statistically significant relationships were observed in the network.

**Figure 9:** 2D patterns identified with minimum pattern score 0.95. Node colors reflect different taxonomy assignments at Phylum level and node sizes are proportional to the average relative abundance of the microorganism across samples. Capital letters inside square brackets represent the lowest taxonomy level identified for each OTU: G (Genus), F (Family), O (Order), C (Class) and P (Phyla). Color of edges indicates relationship type: blue with black arrow (one-way relations), red (co-exclusion), and green (co-presence). See Table S9.xslx for pattern types, scores, OTU taxonomic labels, etc.

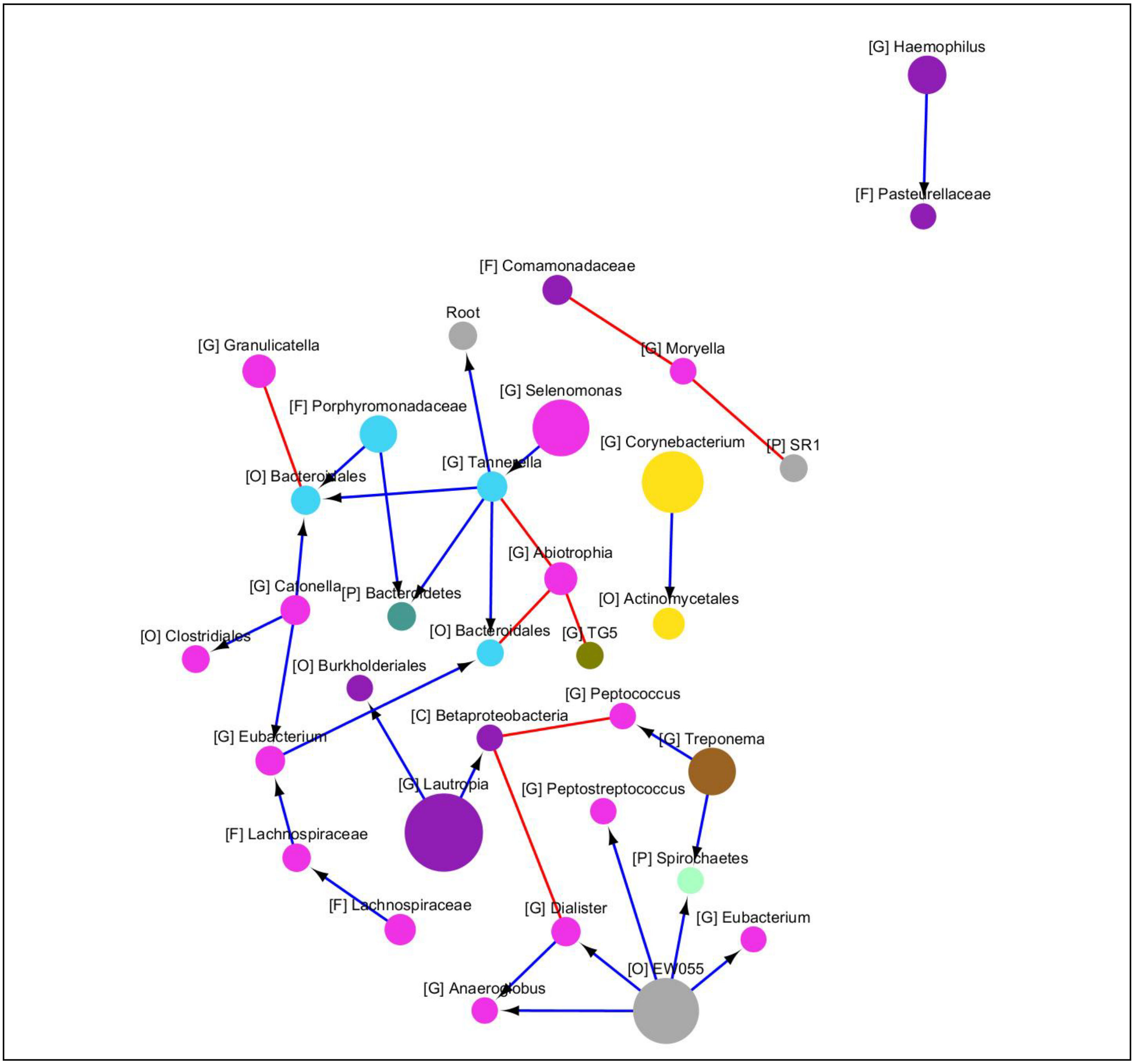

**10. Buccal Mucosa**

**Data summary:** 105 OTUs present across 154 samples were used in the analysis. 29 OTUs involved in 32 statistically significant relationships were observed in the network.

**Figure 10:** 2D patterns identified with minimum pattern score 0.95. Node colors reflect different taxonomy assignments at Phylum level and node sizes are proportional to the average relative abundance of the microorganism across samples. Capital letters inside square brackets represent the lowest taxonomy level identified for each OTU: G (Genus), F (Family), O (Order), C (Class) and P (Phyla). Color of edges indicates relationship type: blue with black arrow (one-way relations), red (co-exclusion), and green (co-presence). See Table S10.xslx for pattern types, scores, OTU taxonomic labels, etc.

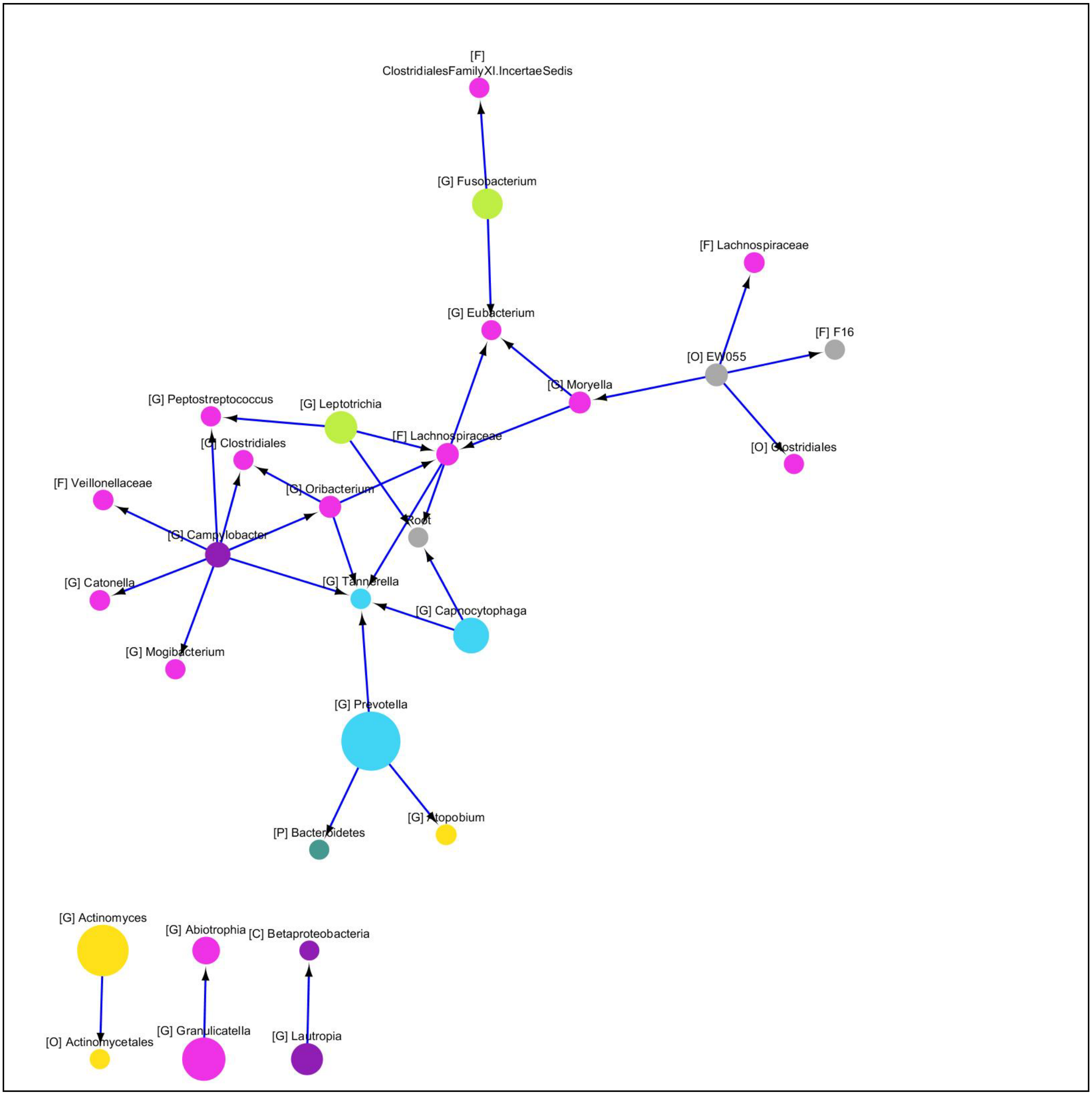

**11. Hard Palate**

**Data summary:** 117 OTUs present across 147 samples were used in the analysis. 2 OTUs involved in 1 statistically significant relationships were observed in the network.

**Figure 11:** 2D patterns identified with minimum pattern score 0.95. Node colors reflect different taxonomy assignments at Phylum level and node sizes are proportional to the average relative abundance of the microorganism across samples. Capital letters inside square brackets represent the lowest taxonomy level identified for each OTU: G (Genus), F (Family), O (Order), C (Class) and P (Phyla). Color of edges indicates relationship type: blue with black arrow (one-way relations), red (co-exclusion), and green (co-presence). See Table S11.xslx for pattern types, scores, OTU taxonomic labels, etc.

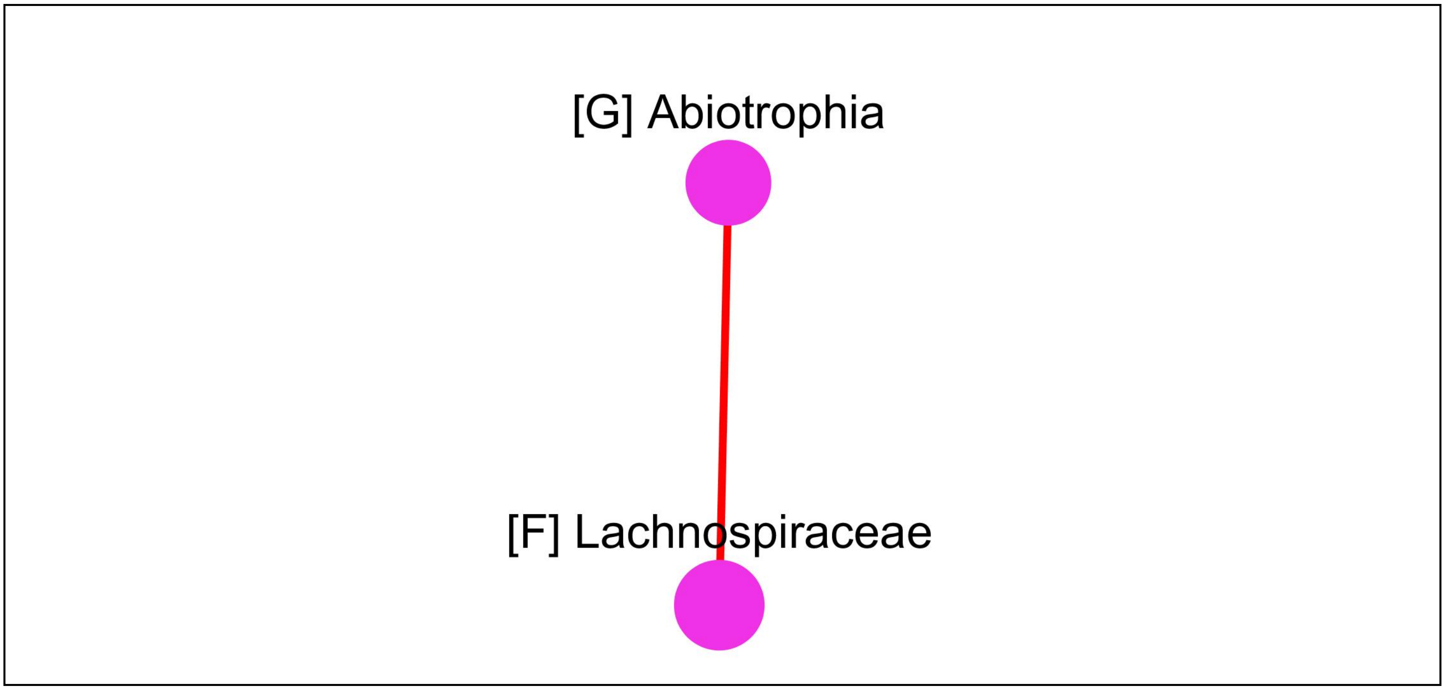

**12. Attached Keratinized Gingiva**

**Data summary:** 74 OTUs present across 161 samples were used in the analysis. 20 OTUs involved in 19 statistically significant relationships were observed in the network.

**Figure 12.** 2D patterns identified with minimum pattern score 0.95. Node colors reflect different taxonomy assignments at Phylum level and node sizes are proportional to the average relative abundance of the microorganism across samples. Capital letters inside square brackets represent the lowest taxonomy level identified for each OTU: G (Genus), F (Family), O (Order), C (Class) and P (Phyla). Color of edges indicates relationship type: blue with black arrow (one-way relations), red (co-exclusion), and green (co-presence). See Table S12.xslx for pattern types, scores, OTU taxonomic labels, etc.

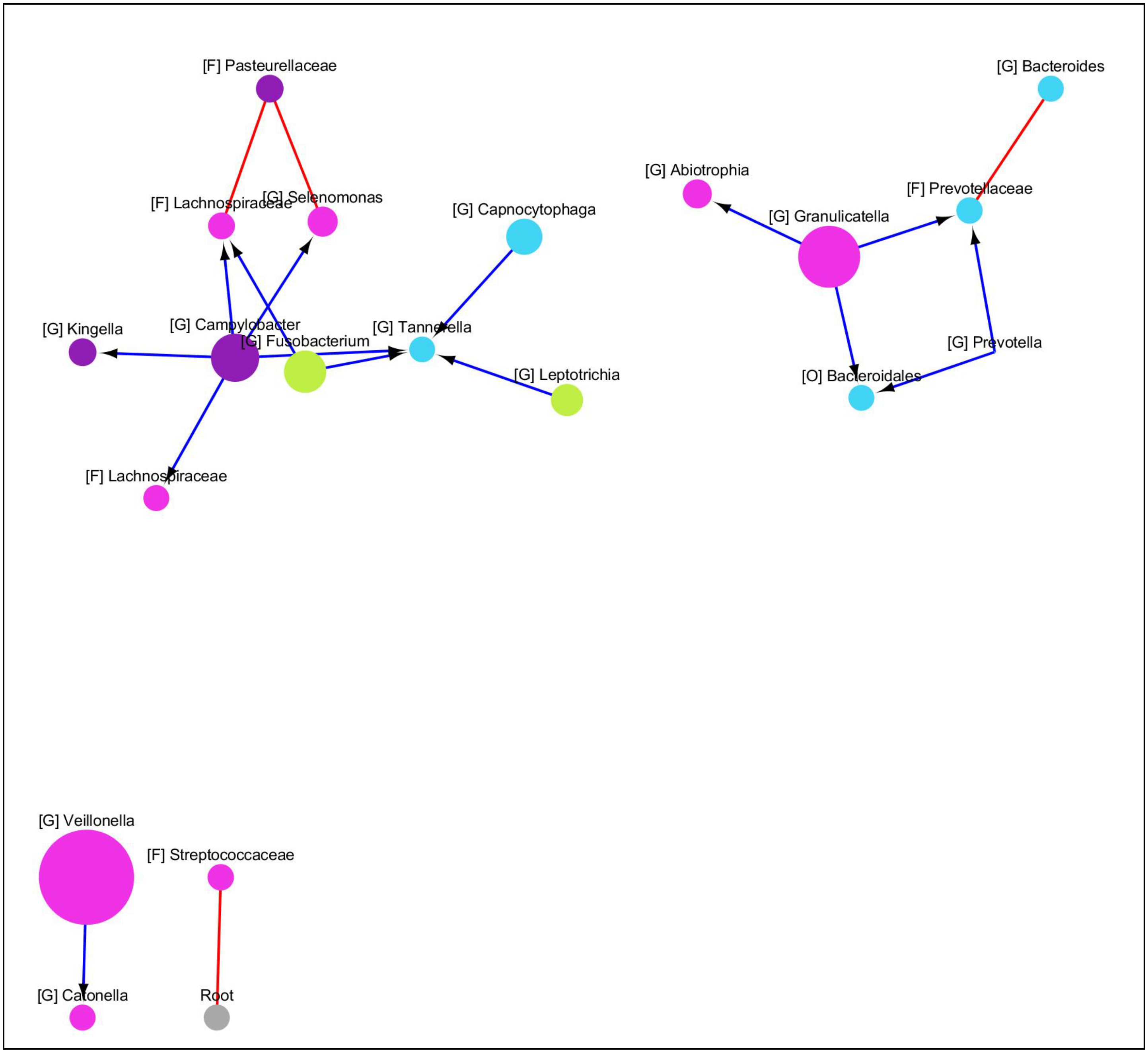

**13. Anterior Nares**

**Data summary:** 129 OTUs present across 127 samples were used in the analysis. 4 OTUs involved in 2 statistically significant relationships were observed in the network.

**Figure 13:** 2D patterns identified with minimum pattern score 0.95. Node colors reflect different taxonomy assignments at Phylum level and node sizes are proportional to the average relative abundance of the microorganism across samples. Capital letters inside square brackets represent the lowest taxonomy level identified for each OTU: G (Genus), F (Family), O (Order), C (Class) and P (Phyla). Color of edges indicates relationship type: blue with black arrow (one-way relations), red (co-exclusion), and green (co-presence). See Table S13.xslx for pattern types, scores, OTU taxonomic labels, etc.

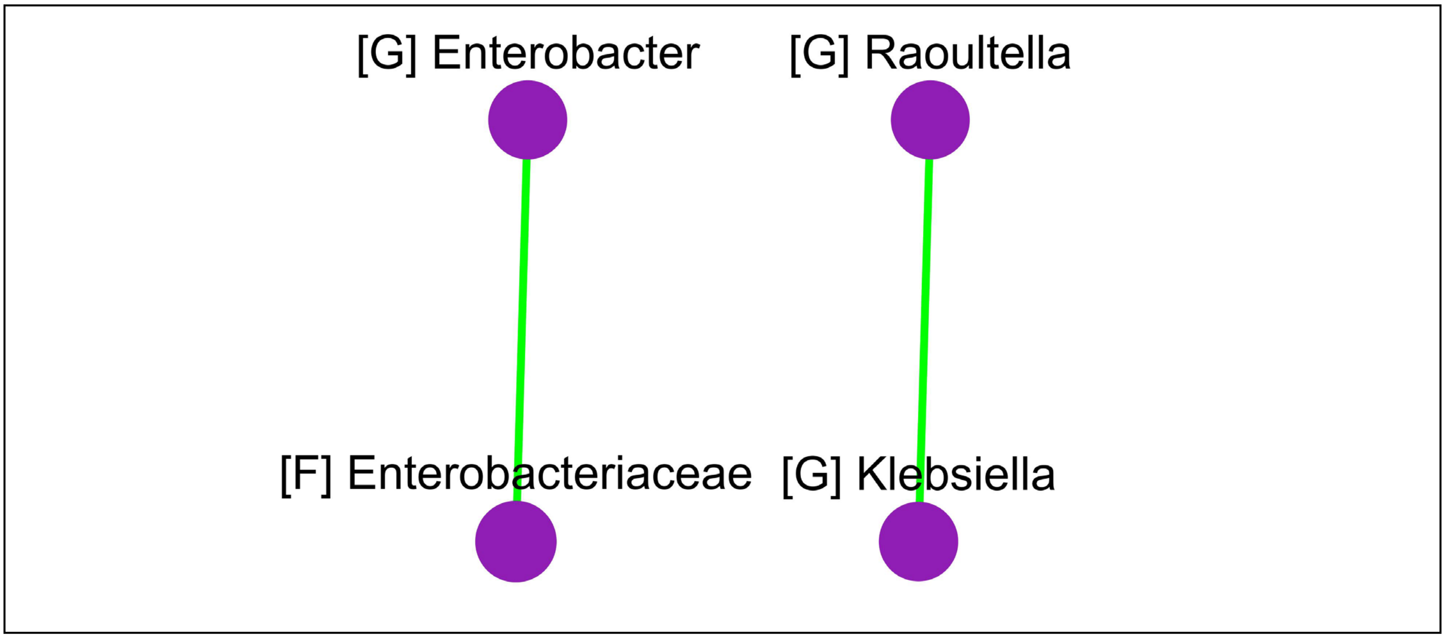

**14. Antecubital Fossa**

**Data summary:** 200 OTUs present across 171 samples were used in the analysis. 25 OTUs involved in 20 statistically significant relationships were observed in the network.

**Figure 14:** 2D patterns identified with minimum pattern score 0.95. Node colors reflect different taxonomy assignments at Phylum level and node sizes are proportional to the average relative abundance of the microorganism across samples. Capital letters inside square brackets represent the lowest taxonomy level identified for each OTU: G (Genus), F (Family), O (Order), C (Class) and P (Phyla). Color of edges indicates relationship type: blue with black arrow (one-way relations), red (co-exclusion), and green (co-presence). See Table S14.xslx for pattern types, scores, OTU taxonomic labels, etc.

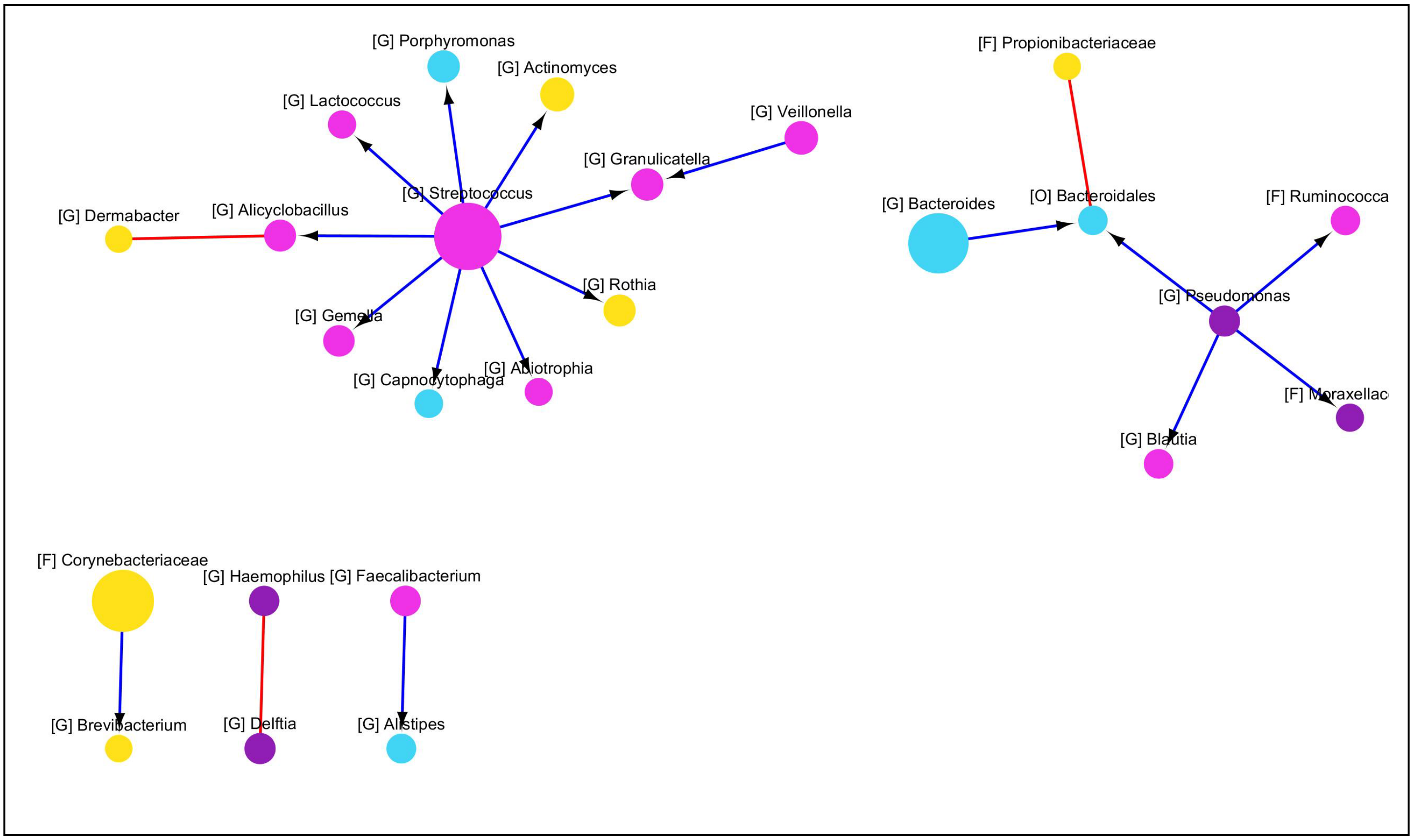

**15. Retroauricular Crease**

**Data summary:** 108 OTUs present across 309 samples were used in the analysis. 18 OTUs involved in 16 statistically significant relationships were observed in the network.

**Figure 15:** 2D patterns identified with minimum pattern score 0.95. Node colors reflect different taxonomy assignments at Phylum level and node sizes are proportional to the average relative abundance of the microorganism across samples. Capital letters inside square brackets represent the lowest taxonomy level identified for each OTU: G (Genus), F (Family), O (Order), C (Class) and P (Phyla). Color of edges indicates relationship type: blue with black arrow (one-way relations), red (co-exclusion), and green (co-presence). See Table S15.xslx for pattern types, scores, OTU taxonomic labels, etc.

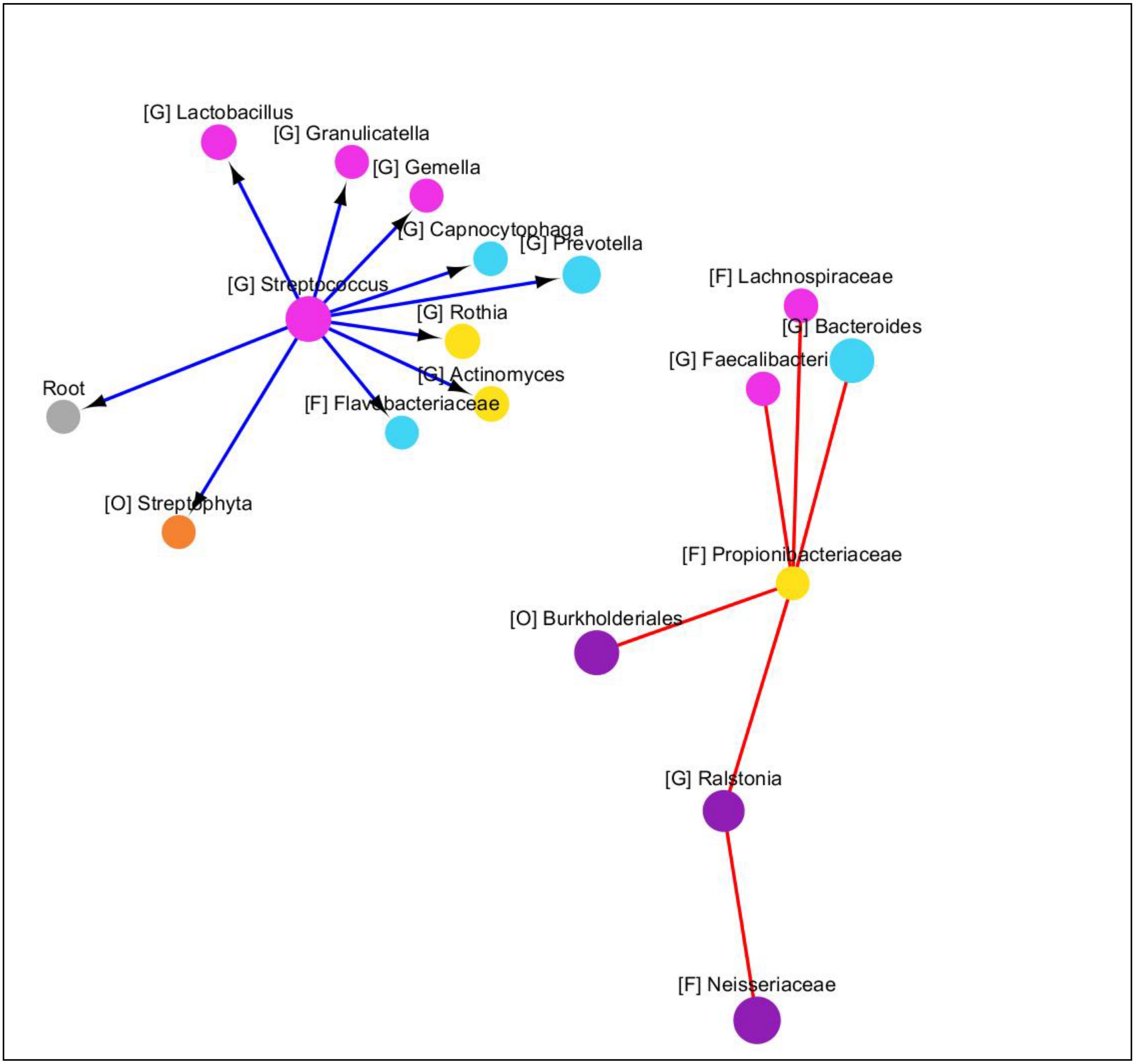

### 16. Palatine Tonsils

**Data summary:** 99 OTUs present across 156 samples were used in the analysis. 23 OTUs involved in 22 statistically significant relationships were observed in the network.

**Figure 16:** 2D patterns identified with minimum pattern score 0.95. Node colors reflect different taxonomy assignments at Phylum level and node sizes are proportional to the average relative abundance of the microorganism across samples. Capital letters inside square brackets represent the lowest taxonomy level identified for each OTU: G (Genus), F (Family), O (Order), C (Class) and P (Phyla). Color of edges indicates relationship type: blue with black arrow (one-way relations), red (co-exclusion), and green (co-presence). See Table S16.xslx for pattern types, scores, OTU taxonomic labels, etc.

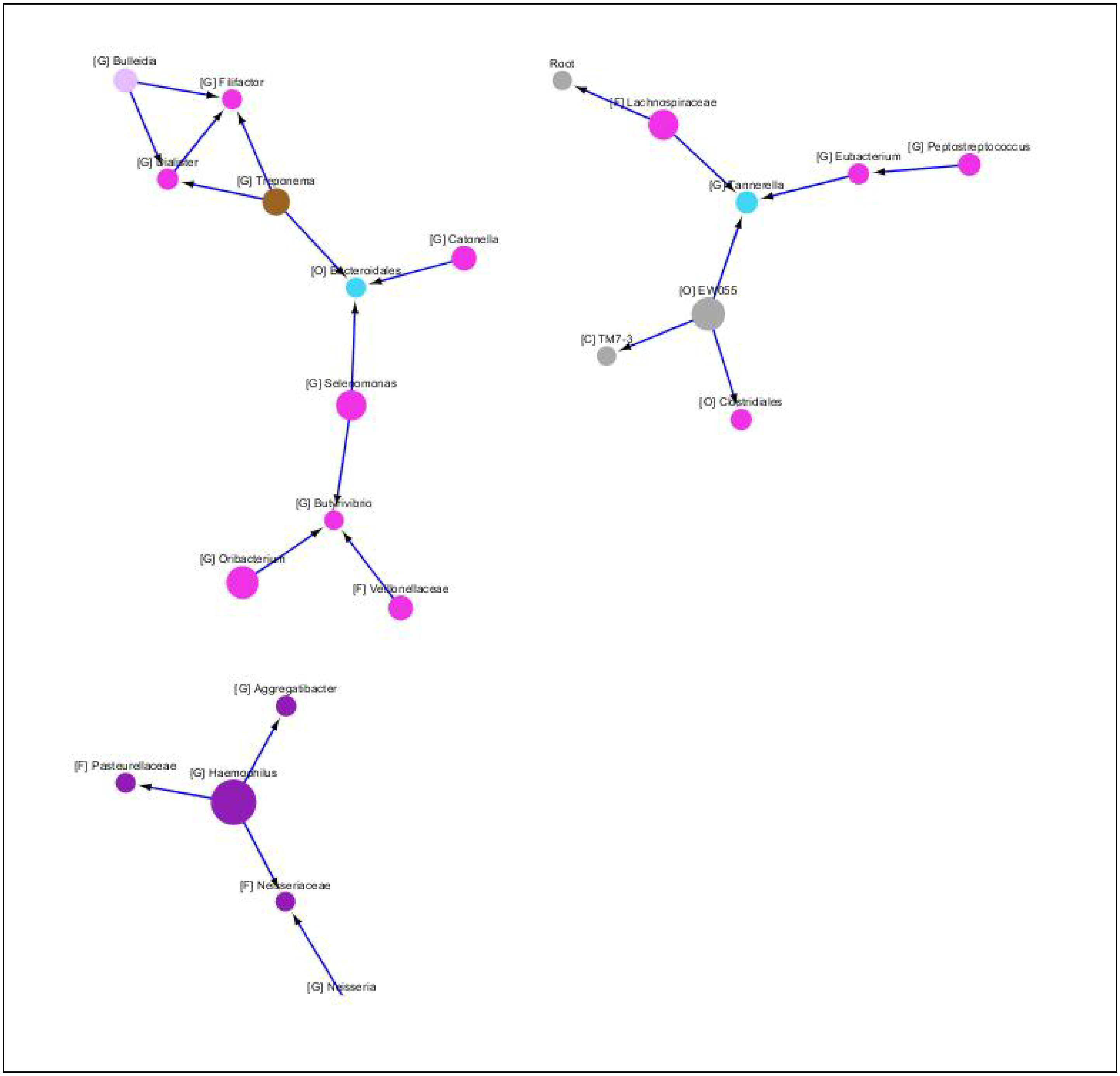
